# Comprehensive longitudinal microbiome analysis of the chicken cecum reveals a shift from competitive to environmental drivers and a window of opportunity for *Campylobacter*

**DOI:** 10.1101/341842

**Authors:** Umer Zeeshan Ijaz, Lojika Sivaloganathan, Aaron Mckenna, Anne Richmond, Carmel Kelly, Mark Linton, Alexandros Stratakos, Ursula Lavery, Abdi Elmi, Brendan Wren, Nick Dorrell, Nicolae Corcionivoschi, Ozan Gundogdu

## Abstract

Chickens are a key food source for humans yet their microbiome contains bacteria that can be pathogenic to humans, and indeed potentially to chickens themselves. *Campylobacter* is present within the chicken gut and is the leading cause of bacterial foodborne gastroenteritis within humans worldwide. Infection can lead to secondary sequelae such as Guillain-Barré syndrome and stunted growth in children from low-resource areas. Despite the global health impact and economic burden of *Campylobacter*, how and when *Campylobacter* appears within chickens remains unclear. As such, there has been a motivation to decrease the number of *Campylobacter* within chickens and thus reduce the risk of infection to humans. The lack of day-to-day microbiome data with replicates, relevant metadata, and a lack of natural infection studies have delayed our understanding of the chicken gut microbiome and *Campylobacter*. Here, we performed a comprehensive day-to-day microbiome analysis of the chicken cecum from day 3 to 35 (12 replicates each day; n=396) combining metadata such as chicken weight and feed conversion rates to investigate what the driving forces are for the microbial changes within the chicken gut over time, and how this relates to *Campylobacter* appearance within a natural habitat setting. We found a rapidly increasing microbial diversity up to day 12 with variation observed both in terms of genera and abundance, before a stabilisation of the microbial diversity after day 20. In particular, we identified a shift from competitive to environmental drivers of microbial community from days 12 to 20 creating a window of opportunity whereby *Campylobacter* appears. *Campylobacter* was identified at day 16 which was one day after the most substantial changes in metabolic profiles observed. In addition, microbial variation over time is most likely influenced by the diet of the chickens whereby significant shifts in OTU abundances and beta dispersion of samples often corresponded with changes in feed. This study is unique in comparison to the most recent studies as neither sampling was sporadic nor *Campylobacter* was artificially introduced, thus the experiments were performed in a natural setting. We believe that our findings can be useful for future intervention strategies and can help elucidate the mechanism through which *Campylobacter* within chickens can be reduced.

## Introduction

Chickens (*Gallus gallus domesticus*) are an important food source for humans with over 50 billion reared annually for meat and eggs (Part et al., 2016). Feed conversion and the health of chickens is heavily dependent on the largely unexplored complex gut microbial community which plays a role in nutrient assimilation, vitamin and amino acid production and prevention of pathogen colonization (Sergeant et al., 2014, Apajalahti, 2005, McNab, 2007, Józefiak et al., 2004). In chickens, the organ with the highest number and variety of bacteria is the cecum (10^10^-10^11^ cells/g) which plays an essential role in the digestion of non-starch polysaccharides (NSPs) found in chicken feed (Jozefiak et al., 2004, Barnes et al., 1972, Bjerrum et al., 2006). The importance of this organ is demonstrated when up to 10% of energy needs can be recovered from a well-functioning cecum (Jozefiak et al., 2004, Hegde et al., 1982), and also that the cecum remains a source of bacterial human infection and a reservoir of antibiotic resistance determinants.

The chicken cecum contracts several times a day releasing contents towards the ileum and the cloaca (Pauwels et al., 2015). Notably the cecal drop contains *Campylobacter*, a Gram-negative spiral shaped bacterium which causes an estimated 400 million human infections each year (Friedman et al., 2000, Walker, 2005). *Campylobacter* causes bloody diarrhoea, fever and abdominal pains in humans and can also cause post infectious sequelae such as Guillain-Barré syndrome which is a potentially fatal paralytic autoimmune illness. In low-resource areas, asymptomatic and occasionally persistent *Campylobacter* infections are common in children younger than one year and correlate with stunted growth and therefore life-long physical and cognitive deficits (Amour et al., 2016). Approximately 80-90% of these infections are attributed to *Campylobacter jejuni*, with poultry as the most important source of human campylobacteriosis within industrialized countries (Humphrey et al., 2007, Mullner et al., 2009, Sheppard et al., 2009). *C*. *jejuni* colonizes the chicken cecum with relatively high numbers (10^9^ CFU per gram) and whereas traditionally was considered a commensal of the chicken gut, more recently has been demonstrated to be pathogenic to the chicken, with this dependent on the genetics of the host and the strain of infection (Van Deun et al., 2008, Hermans et al., 2012, Wigley, 2015, Humphrey et al., 2014, Humphrey et al., 2015). Natural colonisation of chickens is reported to be typically at approximately day 14 of the chicken life cycle, although we do not know how and why this occurs, and what the impact of *Campylobacter* is on the microbiome (Kalupahana et al., 2013, Neill et al., 1984, Thibodeau et al., 2015).

The microbiome of chickens develop rapidly from days 1-3 where *Enterobacteriaceae* dominate, with *Firmicutes* increasing in abundance and taxonomic diversity from approximately day 7 onwards (Ballou et al., 2016, Mancabelli et al., 2016, Danzeisen et al., 2011). Bacterial populations within the chicken gut are subsequently driven by the rearing environment and from the bacteria present in food and water (Connerton et al., 2018). How and when *Campylobacter* appears and the impact on the chicken gut microbiome remains unanswered. The presence of *Campylobacter* has been noted to prompt an increase in *Bifidobacterium* and modify abundances of *Clostridia* and *Mollicutes* (Thibodeau et al., 2015). The identification of a number of hydrogenases within the ceca may lead to a potential hydrogen sink and provide an explanation as to the high abundance of genera such as *Campylobacter* (Sergeant et al., 2014). Comparison of broilers not exposed and exposed to *C. jejuni* at day 6 or day 20 revealed reductions in the relative abundance of operational taxonomic units (OTUs) within the taxonomic family *Lactobacillaceae* and the *Clostridium* cluster XIVa, with specific members of the *Lachnospiraceae* and *Ruminococcaceae* families exhibiting transient shifts in microbial community populations dependent upon the age at which the birds become colonized by *C. jejuni* (Connerton et al., 2018). These studies have enhanced our understanding of the chicken cecal microbiome, however the lack of day-to-day microbiome data, suitable replicate numbers, relevant metadata, and lack of natural infectivity studies have not allowed us to fully appreciate what is occurring in a natural habitat in relation to how and when *Campylobacter* appears within the chicken gut. To answer these questions, in this study we have performed a comprehensive analysis of the chicken cecal microbiome from days 3 to 35, with 12 replicates per day (n=396), correlating additional metadata such as chicken weight and feed conversion rates with *Campylobacter* detection in a natural environmental setting.

## Experimental procedures

### Experimental design, broilers and sample collection

This study was performed using a total of 396 Ross-308 male broiler chickens provided by Moy Park (39 Seagoe Industrial Estate, Portadown, Craigavon, Co. Armagh, BT63 5QE, UK). The birds were divided into 12 pens; each pen contained 33 chickens (Supplementary Figure 1). Birds were raised on three phase diets from day 0 to day 35. Starter diets were offered to the birds from days 0-10, grower diets from days 11-25 and finisher diets from days 26-35. Every 24 hours, a single chicken from each of the 12 pens was removed at random, euthanized by cervical dislocation or anaesthesia combined with cervical dislocation, followed by genomic DNA (gDNA) extracted from the chicken cecum. A total of 16 samples were removed from the final analysis due to poor gDNA quality.

### Poultry growth and performance measurements

The performance parameters investigated were mean body weight (BW_mean), body weight gain (Gain), feed intake (FI) and feed conversion ratio (FCR). Measurements were taken at time points 3-7 days, 8-14 days, 15-24 days and 25-35 days. These variables were then correlated with the microbial community’s composition in various statistical analyses.

### DNA extraction, 16S rRNA amplification and sequencing

Cecal gDNA was extracted using the QIAamp DNA Stool Mini Kit according to the manufacturer’s instructions and stored at −20°C. 16S metagenomic sequencing library construction was performed using Illumina guidelines (Illumina, U.S.A). The 16S ribosomal primers used were V3 (tcgtcggcagcgtcagatgtgtataagagacagcctacgggnggcwgcag) and V4 (gtctcgtgggctcggagatgtgtataagagacaggactachvgggtatctaatcc) (Klindworth et al., 2013, D’Amore et al., 2016). A second PCR step was performed to attach dual indices and Illumina sequencing adapters using the Nextera XT Index kit. Sequencing was performed on the Illumina MiSeq at LSHTM using a v3 300 bp paired-end kit.

### Bioinformatics

Abundance tables were obtained by constructing OTUs (a proxy for species) as follows. Paired-end reads were trimmed and filtered using Sickle v1.200 (Joshi and Fass, 2011) by applying a sliding window approach and trimming regions where the average base quality drops below 20. Following this we applied a 10 bp length threshold to discard reads that fall below this length. We then used BayesHammer (Nikolenko et al., 2013) from the Spades v2.5.0 assembler to error correct the paired-end reads followed by pandaseq (v2.4) with a minimum overlap of 20 bp to assemble the forward and reverse reads into a single sequence spanning the entire V3-V4 region. The above choice of software was as a result of author’s recent work (Schirmer et al., 2015, D’Amore et al., 2016) where it was shown that the above strategy reduces the substitution rates (main form of error) significantly. After having obtained the consensus sequences from each sample, we used the VSEARCH (v2.3.4) pipeline (all these steps are documented in https://github.com/torognes/vsearch/wiki/VSEARCH-pipeline) for OTU construction. The approach is as follows: we pool the reads from different samples together and add barcodes to keep an account of the samples these reads originate from. We then dereplicate the reads and sort them by decreasing abundance and discard singletons. In the next step, the reads are clustered based on 97% similarity, followed by removing clusters that have chimeric models built from more abundant reads (--uchime_denovo option in vsearch). A few chimeras may be missed, especially if they have parents that are absent from the reads or are present with very low abundance. Therefore, in the next step, we use a reference-based chimera filtering step (--uchime_ref option in vsearch) using a gold database (https://www.mothur.org/w/images/f/f1/Silva.gold.bacteria.zip). The original barcoded reads were matched against clean OTUs with 97% similarity (a proxy for species level separation) to generate OTU table (a total of 18,588 unique sequences) for n=382 samples.

The representative OTUs were then taxonomically classified against the SILVA SSU Ref NR database release v123 database with assign_taxonomy.py script from the Qiime (Caporaso et al., 2010) workflow. To find the phylogenetic distances between OTUs, we first multisequence aligned the OTUs against each other using Kalign v2.0.4 (Lassmann and Sonnhammer, 2005) (using the options -gpo 11 -gpe 0.85) and then used FastTree v2.1.7 (Price et al., 2010) to generate the phylogenetic tree in NEWICK format. Finally make_otu_table.py from Qiime workflow was employed to combine abundance table with taxonomy information to generate biome file for OTUs. Tax4Fun (Asshauer et al., 2015) was used to predict the functional capabilities of microbial communities based on 16S rRNA datasets (all prokaryotic KEGG organisms are available in Tax4Fun for SILVA v123 and KEGG database release 64.0) and then utilising ultrafast protein classification (UProC) tool (Meinicke, 2015) to generate metabolic functional profiles after normalising the data for 16S rRNA gene copy numbers. In Tax4Fun, we used MoP-Pro approach (Asshauer and Meinicke, 2013) to give pre-computed 274 KEGG Pathway reference profiles. Although Tax4Fun based metabolic prediction is constrained by the taxa available in the reference database, it gives a statistic called fraction-of-taxonomic-units-unexplained (FTU) which reflects the amount of sequences assigned to a taxonomic unit and not transferable to KEGG reference organisms. This can be used as a measure of confidence in trusting the predictions. Summary statistics of FTUs returned in this study are as follows: 1st Quantile:0.09129; Median:0.13995; Mean:0.14902; and 3^rd^ Quantile:0.19800 (Figure 1h). Thus, on average metabolic profiles of ∼86% of the taxa were present and therefore with this high representation, we used the pathways in the statistical analysis.

### Statistical analysis

Statistical analyses were performed in R using the tables and data generated as above as well as the meta data associated with the study. For community analysis (including alpha and beta diversity analyses) we used the vegan package (Oksanen et al., 2015). For alpha diversity measures, we calculated: *Richness*, estimated number of species/features per sample; and *Shannon* entropy: a commonly used index to characterise species diversity. To calculate Unifrac distances (that account for phylogenetic closeness), we used the phyloseq (McMurdie and Holmes, 2013) package. Nonmetric Distance Scaling (NMDS) plot of community data (OTUs) used different distance measures (Vegan’s metamds() function): *Bray-Curtis*, considers the species abundance count; *Unweighted Unifrac,* considers the phylogenetic distance between the branch lengths of OTUs observed in different samples without taking into account the abundances; and *Weighted Unifrac,* unweighted unifrac distance weighted by the abundances of OTUs. The samples are grouped for different treatments as well as the mean ordination value and spread of points (ellipses were drawn using Vegan’s ordiellipse() function that represent the 95% confidence interval of the standard errors of the groups).

To understand multivariate homogeneity of groups dispersion (variances) between multiple conditions, we used Vegan’s betadisper() function in which the distances between objects and group centroids are handled by reducing the original distances (BrayCurtis, Unweighted Unifrac, or Weighted Unifrac) to principal coordinates and then performing ANOVA on them. We used Vegan’s adonis() for analysis of variance using distance matrices (BrayCurtis/Unweighted Unifrac/Weighted Unifrac) i.e., partitioning distance matrices among sources of variation (Grouping type i.e., weeks, body weight, feed intake, feed conversion ratio etc.). This function, henceforth referred to as PERMANOVA, fits linear models to distance matrices and uses a permutation test with pseudo-F ratios.

To find OTUs that are significantly different between multiple conditions (days/weeks), we used DESeqDataSetFromMatrix() function from DESeq2 (Love et al., 2014) package with the adjusted p-value significance cut-off of 0.05 and log2 fold change cut-off of 2. This function uses negative binomial GLM to obtain maximum likelihood estimates for OTUs log fold change between two conditions. Then Bayesian shrinkage is applied to obtain shrunken log fold changes subsequently employing the Wald test for obtaining significances. To find KEGG pathways significantly up/down-regulated between multiple conditions (days/weeks), the Kruskal-Wallis test was used with p-values adjusted for multiple comparisons using the fdrtool package (Klaus and Strimmer, 2013, Klaus and Strimmer, 2015).

We performed Local Contribution to Beta Diversity (LCBD) analysis (Legendre and De Caceres, 2013) by using LCBD.comp() from adespatial package (Dray et al., 2018). We used the Hellinger distance (abundances), unweighted (phylogenetic distance) and weighted Unifrac (phylogenetic distance weighted by abundance) dissimilarities. LCBD gives the sample-wise local contributions to beta diversity that could be derived as a proportion of the total beta diversity. In the context of this longitudinal study, it provides a mean to show how markedly different the microbial community structure of a single sample is from the average (with higher LCBD values representing outliers), and also provides a mean to show when the community structure has stabilised in a temporal setting.

To characterise the phylogenetic community composition within each sample whether the microbial community structure is stochastic (driven by competition among taxa) or deterministic (driven by strong environmental pressure i.e. host environment), we quantified: mean-nearest-taxon-distance (MNTD) and the nearest-taxon-index (NTI) using mntd(), and ses.mntd(); and mean-phylogenetic-diversity (MPD) and nearest-relative-index (NRI) using mpd() and ses.mpd() function from the picante (Kembel et al., 2010) package. NTI and NRI represent the negative of the output from ses.mntd() and ses.mpd(), respectively. They also quantify the number of standard deviations that the observed MNTD/MPD is from the mean of the null distribution (999 randomization by using null.model=’richness’ in the ses.mntd() and ses.mpd() functions and only considering the taxa as present/absent without taking their abundances). We used the top 1000 most abundant OTUs for calculation of these measures based on the recommendations given in (Stegen et al., 2012).

We used the “BVSTEP” routine (Clarke and Ainsworth, 1993), an algorithm that searches for highest correlation (Mantel test) between dissimilarities of a fixed and variable multivariate datasets using bvStep() from sinkr package (Taylor, 2014) by permuting through 2^n^-1 possible combinations of features in the variable dataset. Testing all feature combinations is unrealistic and computationally intractable when the feature space is high (18,588 OTUs in our case). Thus, we used the abundance table with 1000 most abundant OTUs (with the premise that the most abundant species that may have a significant role to play) to best correlate with the overall similarities given all the OTUs (18,588 in our case). This analysis is complimentary to the differential analysis and identified the OTUs that were causing the major shifts in beta diversity. The phylogenetic tree and annotations summarizing the findings of this study were drawn using Evolview (http://www.evolgenius.info/evolview/).

We considered analyses on two different groupings of the sample data, comparison of microbial profiles on a daily basis to reveal temporal patterns, and on a weekly basis (4 weeks), primarily because the poultry growth and performance parameters were recorded on a weekly basis. The statistical scripts and workflows for all above can be found at http://userweb.eng.gla.ac.uk/umer.ijaz#bioinformatics

## Results

### Daily diversity patterns converge to a stable community as we go forward in time

Although alpha diversity (Shannon) on microbial counts (Figure 1a) shows a rapid increase over the first ten days, it follows a plateauing effect where the microbiome normalises at approximately day 12. This is in line with previous reports whereby the gastrointestinal (GI) tract of poultry comes into contact with exogenous microorganisms immediately after hatch and as the host grows, this microbiome becomes highly diverse until it reaches a relatively stable yet dynamic state (Pan and Yu, 2014). The same temporal phenomenon can be observed when considering local contributions to beta diversity based on abundance count (Hellinger distance; Figure 1b). When considering phylogenetic distances only (Unweighted Unifrac; Figure 1c), although the decrease in beta diversity contributions is marginally slower than the abundance counts counterpart, there is a sudden increase around day 20. Using both abundances and phylogenetic distances this seems to disappear (Weighted Unifrac; Figure 1d). It should be noted that a higher LCBD value suggests the diversity patterns of a sample is markedly different from the rest of the samples in an average sense. In contrast, the level of microbial diversity between the different pens was relatively stable (results not significant and thus not shown) suggesting less or no variability amongst pens. *Campylobacter* was detected in three chickens from the 12 pens at day 16 (Figure 1a). This is in line with previous reports where natural colonisation of chickens has been reported at approximately day 14 of the chicken life cycle (Kalupahana et al., 2013, Neill et al., 1984, Thibodeau et al., 2015, Hermans et al., 2011). *Campylobacter* was also identified in one of the chickens at day 3 and previously it has also been reported that chickens between 0 and 3 days of age can become infected with *Campylobacter* (Cawthraw et al., 1996).

**Figure 1:**
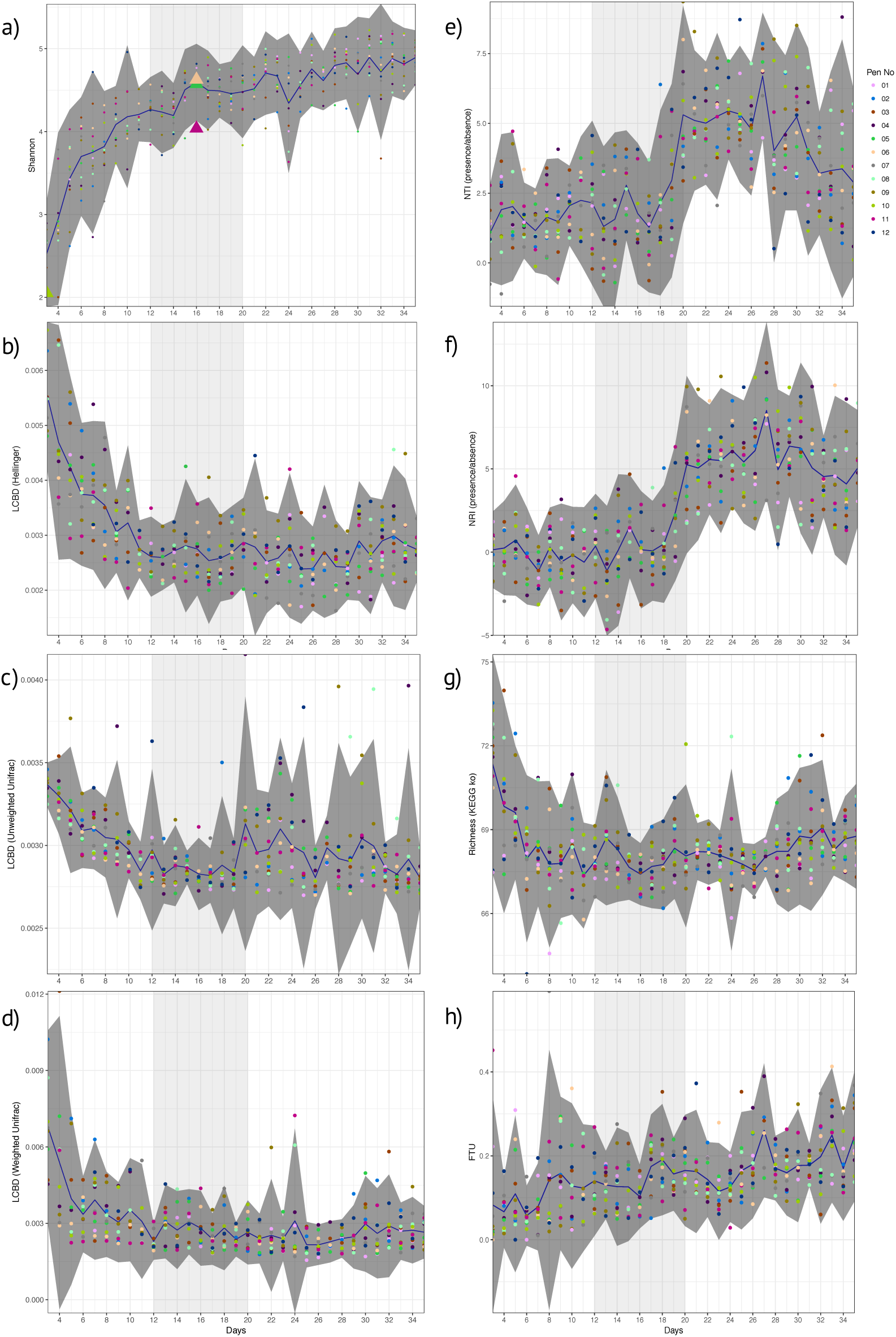
Day-wise statistical measures calculated on the microbiome data. a) Shannon entropy with first appearance of *Campylobacter* (>=5 sequences) highlighted as triangles. b), c), and d) Local contribution to beta diversity (LCBD) calculated by using Hellinger transform on the microbial counts, Unweighted Unifrac dissimilarity (phylogenetic distances only), and Weighted Unifrac dissimilarity (phylogenetic distances weighted with abundance counts) respectively e) and f) Nearest-Taxon-Index (NTI) and nearest-relative-index (NRI) considering presence/absence of OTUs in samples g) Richness calculated as exponentiation of Shannon entropy on the proportional representation of KEGG pathways on samples, and h) fraction-of-taxonomic-units-unexplained (FTU) calculated on each sample. In all subfigures, the mean value is represented by solid blue line with 95% confidence interval of standard deviation given as dark shaded region around the mean. The samples are coloured with respect to the pens they originate from. Based on the analysis given in this study, we have identified days 12 to 20 of importance and are thus highlighted as lighter shaded regions.

### Window of opportunity for Campylobacter between day 12 and day 20

Next, we explored ecological drivers of microbial community to determine whether there is any environmental pressure (host environment) responsible for assemblage of microbial community or if it is driven purely by competition. Using NTI and NRI (Figure 1e, f), one can observe a step function response around day 12. For a single community, NTI/NRI greater than +2 indicates strong phylogenetic clustering (driven by environmental filtering) and less than −2 indicates phylogenetic overdispersion (environment has little or no role to play). Since chicken ceca are already a constrained environment to begin with (as opposed to real environmental datasets), the lower bound of −2 may not be feasible and hence the values should be taken relatively with an increasing value implying increasing host environmental pressure. It should be noted that whilst NRI reflects the phylogenetic clustering in a broad sense (whole phylogenetic tree) with the negative values representing evenly spread community, NTI focuses more on the tips of the tree with positive values of NTI indicating that species co-occur with more closely related species than expected, and negative values indicate that closely related species do not co-occur. We have chosen presence/absence of species while calculating these measures without taking into account the abundances as they mask the phenomenon similar to LCBD profiles (Figure 1c, d). When we consider differential analysis of OTUs (Supplementary Table 1), we can notice that between days 9 and 11 there is a high proportion of OTUs that were log2 fold different. After day 20, we also observe the same between days 26 and 28 with the changes in phylogenetic structure responsible for peaks in NTI/NRI. Interestingly, chickens were raised on three phase diets; starter diets (days 0-10), grower diets (days 11-25) and finisher diets (days 26-35). The high proportion of OTUs that were log2 fold different between days 26 and 28 may be attributed to the change in feed from grower to finisher feed. Since the NTI/NRI are already significantly higher than 2, we do not consider this as an upper bound and revert back to day 20 as an upper bound for the window. Based on beta dispersion analysis (Table 1), we observe days 11 to 13 and days 19 to 21 when the dispersions of the microbial communities are changing significantly. The alteration in the chicken feed from starter diet (days 0-10) to grower diets (days 11-25) may also play a role in the significant beta dispersion between days 11 to 13, although the feed change does not seem a likely explanation for days 19 to 21. For completeness we also generated differential analysis of genus level where *Campylobacter* was identified as being significantly down-regulated between day 16 to day 17 (Supplementary Table 2).

**Table 1:**
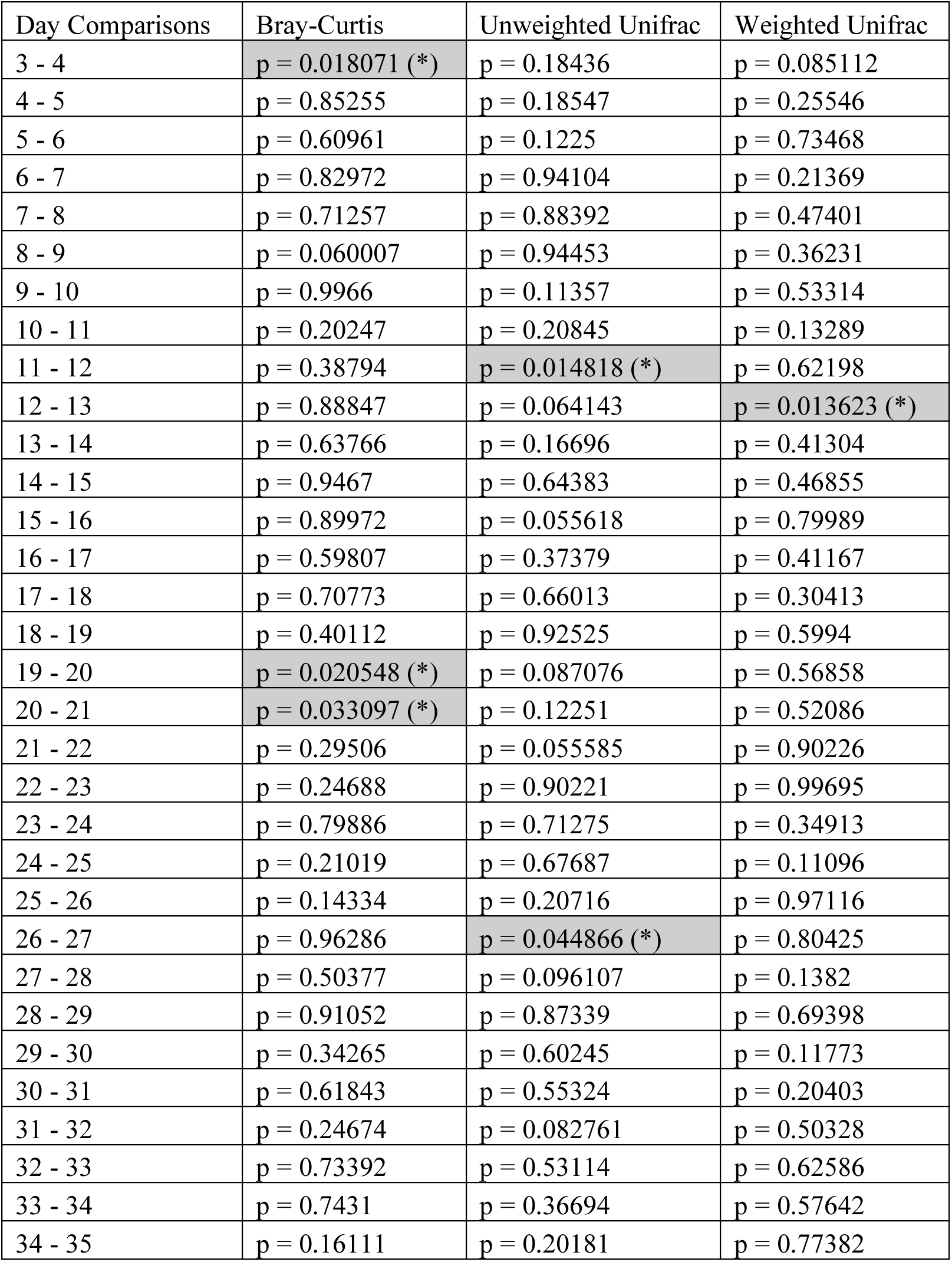
Statistics for beta dispersion comparison on daily microbiome data.

If we consider the richness of metabolic pathways (Figure 1g), we notice that they achieve stability before the microbial community at around day 6 with no obvious patterns to suggest anything apparent between day 12 and day 20 other than a marginal decrease to day 16 and increasing again onwards. However, if we consider the differential expression analysis of pathways (Supplementary Table 3), we can notice a large proportion of these pathways changing between day 14 and 15, a day before *Campylobacter* was first observed. We identified a reduction in lysine degradation (ko00310) from day 14 to day 15, and an increase in D-Alanine metabolism (ko00473) from day 14 to day 15. *C. jejuni* typically cannot utilize sugars as a carbon source as it lacks the glycolytic enzyme phosphofructokinase and so depends on the availability of free amino and keto acids scavenged from the host or from the intestinal microbiome (Parkhill et al., 2000, Velayudhan and Kelly, 2002, Lee and Newell, 2006). *C. jejuni* utilizes serine, aspartate, glutamate and proline preferentially as nutritional substrates *in vitro* with serine catabolism required for colonization of the intestinal tract (Leach et al., 1997, Elharrif and Mégraud, 1986, Velayudhan et al., 2004, Hendrixson and DiRita, 2004). Amino acids can also potentially be deaminated to a small number of intermediates that can directly feed into the central metabolism, including pyruvate (from serine and alanine), oxaloacetate (from aspartate), and 2-oxoglutarate (from glutamate) (Velayudhan et al., 2004). The variation of such metabolic pathways may give an indication as to the appearance of *Campylobacter* at this time point. We also identified a reduction from days 14 to day 15 of a number of pathways relating to specific bacteria; *Vibrio cholerae* pathogenic cycle (ko05111; Biofilm formation - *Vibrio cholerae*), *Escherichia coli* (ko05130; Pathogenic *Escherichia coli* infection), *Salmonella* species (ko05132; *Salmonella* infection). In addition, we identified a reduction from day 14 to 15 of Bacterial secretion systems (ko03070). Future studies are needed to elucidate and confirm the predicted pathways. In view of these findings, *Camplyobacter* appears in day 16 within this window of opportunity (Figure 1) where there exists a shift from competitive to environmental drivers of microbial community, with day 16 lying immediately after the most substantial changes in metabolic profiles observed over the whole period.

### Analysis of dominant bacterial group over time

Analysis of the 50 most abundant genera (Supplementary Figure 2) identified trends that have been reported previously in that the chicken microbiome contains *Enterobacteriaceae* at early days of development, and that *Firmicutes* increase in abundance and taxonomic diversity over time (Ballou et al., 2016, Mancabelli et al., 2016, Danzeisen et al., 2011). *Escherichia.Shigella* (Phylum *Proteobacteria*; Family *Enterobacteriaceae*) was identified as being highly abundant at day 3 and showed a general reduction up to approximately day 7. *Escherichia.Shigella* was also noted to be present after day 28. This pattern was observed for *Eisenbergiella* (Phylum *Firmicutes*; Family *Lachnospiraceae*) which displayed a decrease from early time points, but remained present throughout. This pattern was also observed for *Ruminiclostridium* (Phylum *Firmicutes*; Family *Ruminococcaceae*) which however was not in the abundant genera after day 23. *Flavonifractor* (Phylum *Firmicutes*; Family -) was identified consistently at early time points but was rarely abundant after day 19. *Enterobacter* (Phylum *Proteobacteria*; Family *Enterobacteriaceae)* was only observed at days 3 and 4 and was not abundant at any other time points. Here we identified that *Ruminiclostridium.5* and *Ruminiclostridium.9* (Phylum *Firmicutes*; Family *Ruminococcaceae*) were consistently present throughout at a relatively significant level of abundance. This was also the case for *Anaerotruncus* (Phylum *Firmicutes*; Family *Clostridiaceae*), but at a lower level of abundance, especially before day 7. *Faecalibacterium* (Phylum *Firmicutes*; Family *Clostridiaceae*) was rarely abudant at early time points, however was observed consistently at a relative high abundance after day 14. *Lachnoclostridium* (Phylum *Firmicutes*; Family *Lachnospiraceae*) was found to be present throughout with a relatively high level of fluctuation. Certain genera such as *Ruminococcaceae. UCG.005* and *Ruminococcaceae. UCG.014* (Phylum *Firmicutes*; Family *Ruminococcaceae*) were not abundant at high levels at early time points however increased significantly at approximately days 16-19. Finally, *Megamonas* (Phylum *Firmicutes*; Family *Veillonellaceae*) and *Intestinimonas* (Phylum *Firmicutes*; Family -) were not abundant throughout most time points, before appearing post day 22-25 onwards.

### Weekly microbial profiles and analysis of poultry performance metadata

The metadata collected here included Bird Weight (BW_Mean; grams), Body Weight Gain (Gain; g/bird), Feed Intake (FI), Feed Conversion Ratio (FCR), and was recorded on a weekly basis where we have considered grouping the microbiome samples accordingly; days 03-07 (week 1), days 08-14 (week 2), days 15-24 (week 3), and days 25-35 (week 4). As is the case with the daily microbiome profile, alpha diversity (rarefied richness and Shannon; Figure 2a) increases over time, however, due to the nature of this grouping, we lose the plateauing effect over time. In accordance with daily analysis, we can see a major shift in the parameters as we transition from days 08-14 to days 15-24 (Figure 2b). FCR in particular increases substantially in this period remaining stable for week 4 (days 25-35). Gain is also significantly elevated in this transition period (days 08-14 to days 15-24) when compared to other periods. In terms of beta diversity (Figure 2c), we observe the samples more sparsely spread in the first week (days 03-07) as compared to other weeks on abundance (Bray-Curtis) alone. The phylogenetic dispersion (Unweighted Unifrac) on the other hand is more preserved. We can also notice a gradient forming with later weeks more or less close to suggest convergence as we established in the case of daily profiles. Based on beta dispersion analysis (Table 2), we can notice that the dispersion in week 1 is significantly different to other weeks with 16%, 6%, and 17% variability in microbial community explained by PERMANOVA using counts alone (Bray-Curtis), phylogenetic distance alone (Unweighted Unifrac), and combination of two (Weighted Unifrac), respectively. With this grouping, main sources of variation are then the distribution of species rather than their phylogenetic relatedness. The metadata explains 10-12% variability (all significant) in terms of counts alone (Bray-Curtis) with 3-6% in terms of phylogeny (Unweighted Unifrac). For the sake of completeness, we also performed differential analysis of OTUs and pathways on a consecutive weekly basis (lower halves of Supplementary Tables 1, 2 and 3); however, these should be interpreted with great care as main source of variability are the daily changes and grouping samples on weekly basis will always return more significant OTUs and pathways.

**Figure 2:**
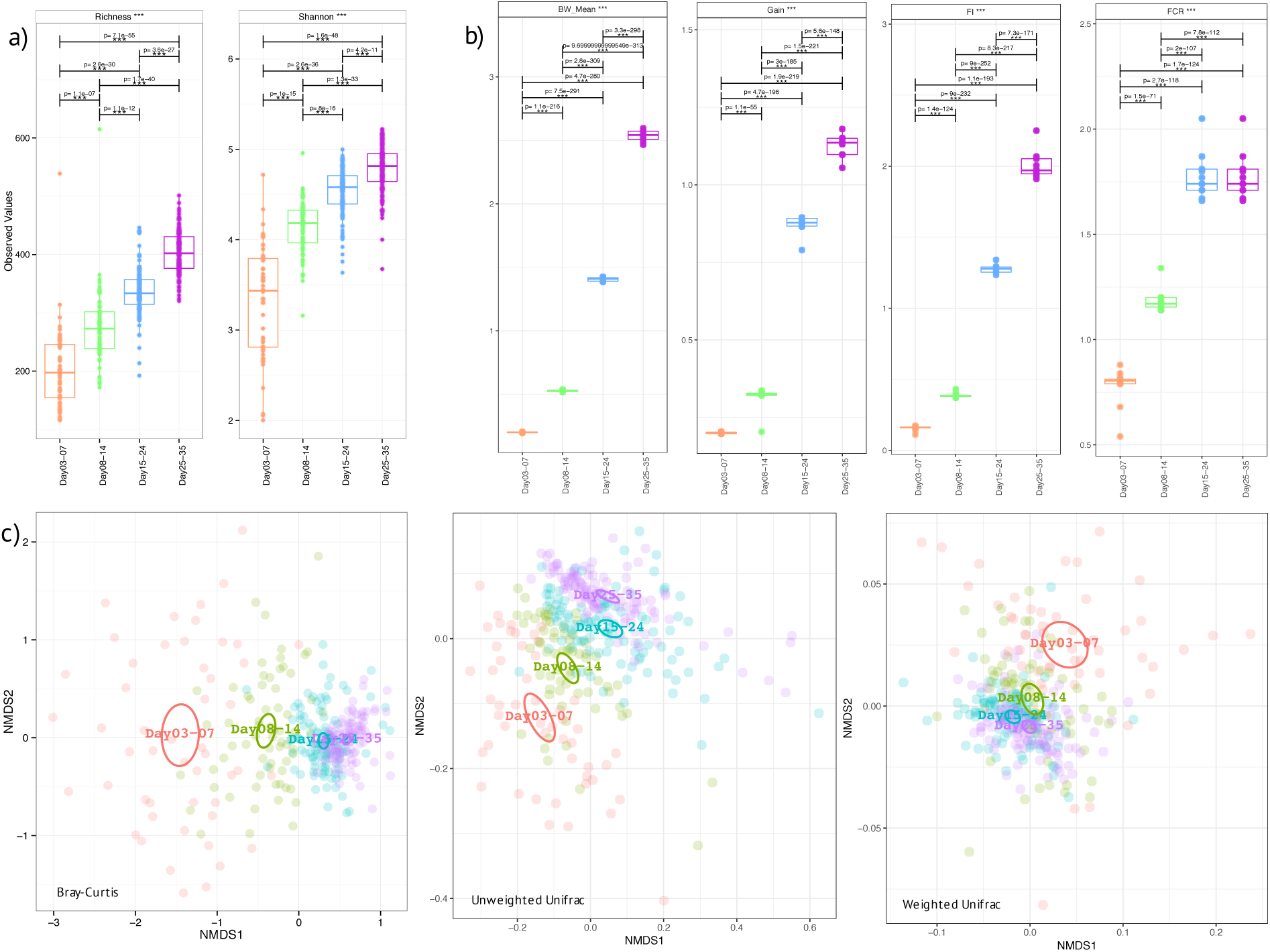
Week-wise measures calculated on the microbiome data a) Alpha diversity measures: Richness (after rarefying the samples to minimum library size) and Shannon entropy b) Extrinsic parameters calculated on weekly basis were mean body weight (BW_mean), body weight gain (Gain), feed intake (FI), feed conversion ratio (FCR), and c) Beta diversity measures using Bray-Curtis (counts), Unweighted Unifrac (phylogenetic distance), and Weighted Unifrac (phylogenetic distance weighted by abundance counts). In a) and b) we have performed pair-wise ANOVA and where significant the pairs were connected with p-values drawn on top. In c) the ellipses represent the 95% confidence interval of the standard error of the ordination points of a given grouping with labels drawn at the centre (mean) of the ordination points.

**Table 2:** Statistics for pairwise beta dispersion and PERMANOVA when using different dissimilarity measures on weekly microbiome data. In beta dispersion analysis, the pair-wise differences in distances from group centre/mean were subjected to ANOVA after performing Principle Coordinate Analysis, and if significant (p≤0.05) the values are shown. In PERMANOVA analysis, R^2^ represents the proportion of variability explained, for example, using “Groups” and “Bray-Curtis” dissimilarity, the weeks explain 16.8% variability in microbial community structure.

| Beta dispersion |  | Bray-Curtis | Unweighted Unifrac | Weighted Unifrac |
| --- | --- | --- | --- | --- |
| Day03-07 | Day08-14 | $p = 0.0061142$ (**) | $p = 0.00014712$ (***) | $p = 9.6914e-05$ (***) |
| | Day15-24 | n.s. | $p = 0.010418$ (*) | $p = 2.5203e-09$ (***) |
| | Day25-35 | $p = 0.042066$ (*) | $p = 0.00015112$ (***) | $p = 3.5789e-12$ (***) |
| Day08-14 | Day15-24 | $p = 0.00077017$ (***) | n.s. | $p = 0.019953$ (*) |
| | Day25-35 | n.s. | n.s. | $p = 0.0011717$ (**) |
| Day15-24 | Day25-35 | $p = 0.0075651$ (**) | $p = 0.020128$ * | n.s. |
| PERMANOVA |  |  |  |  |
| Groups | | $R^2 = 0.16763$ ($p = 0.001$) (***) | $R^2 = 0.06048$ ($p = 0.001$) (***) | $R^2 = 0.17577$ ($p = 0.001$) (***) |
| BW Mean | | $R^2 = 0.11721$ ($p = 0.001$) (***) | $R^2 = 0.03964$ ($p = 0.001$) (***) | $R^2 = 0.08723$ ($p = 0.001$) (***) |
| FI | | $R^2 = 0.11856$ ($p = 0.001$) (***) | $R^2 = 0.04069$ ($p = 0.001$) (***) | $R^2 = 0.09301$ ($p = 0.001$) (***) |
| FCR | | $R^2 = 0.1086$ ($p = 0.001$) (***) | $R^2 = 0.03842$ ($p = 0.001$) (***) | $R^2 = 0.11787$ ($p = 0.001$) (***) |
| Gain | | $R^2 = 0.11886$ ($p = 0.001$) (***) | $R^2 = 0.04146$ ($p = 0.001$) (***) | $R^2 = 0.0998$ ($p = 0.001$) (***) |

### Key species representing majority of the shift in community dynamics

In addition to differential analysis on OTUs (Supplementary Table 1) which returned OTUs that were log2 fold different between consecutive days, we also considered the subset analysis where we imploded the abundance table to the minimum set of OTUs, the resulting reduced-order abundance table correlated highly with the full table by preserving the beta diversity between the samples (Table 3). To see how much variability is lost, the PERMANOVA with full OTU table (18,588 OTUs) is provided as a reference. The 17 OTUs listed represent only ∼2% (subset S1) loss in variability and thus represent the main OTUs that are driving the community dynamics. In terms of metadata, the loss in variability is ∼1% (subset S1). The subset of the phylogenetic tree of these OTUs in addition to those selected in the differential analysis (daily comparisons), a total of 110 OTUs, was then extracted and annotated with these analysis in Figure 3 along with taxonomy information. It can be seen that majority of these (>50%) belong to *Firmicutes* (*Bacillaceae*, *Ruminococcaceae*, *Lachnospiracaeae*, *Lactobacillaceae*, *Peptostreptococcaceae*, and *Clostridiales vadin BB60 group*), with a small proportion belonging to *Actinobacteria* (*Coriobacteriacaea*), *Tenericutes* (*Mollicutes RF9*), and *Proteobacteria* (*Enterobacteriaceae* including *Escherichia.Shigella* as mentioned before).

**Table 3:**
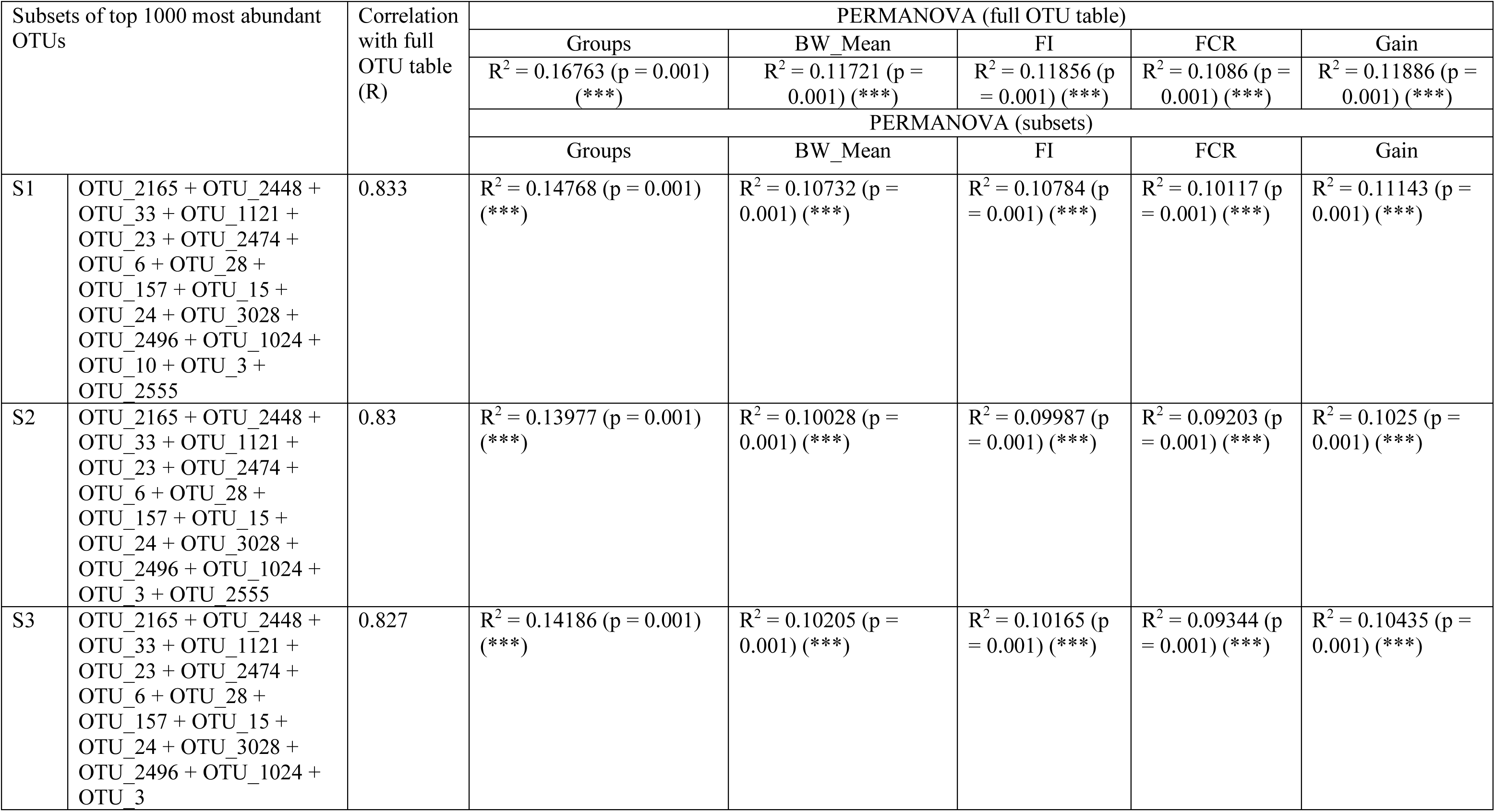
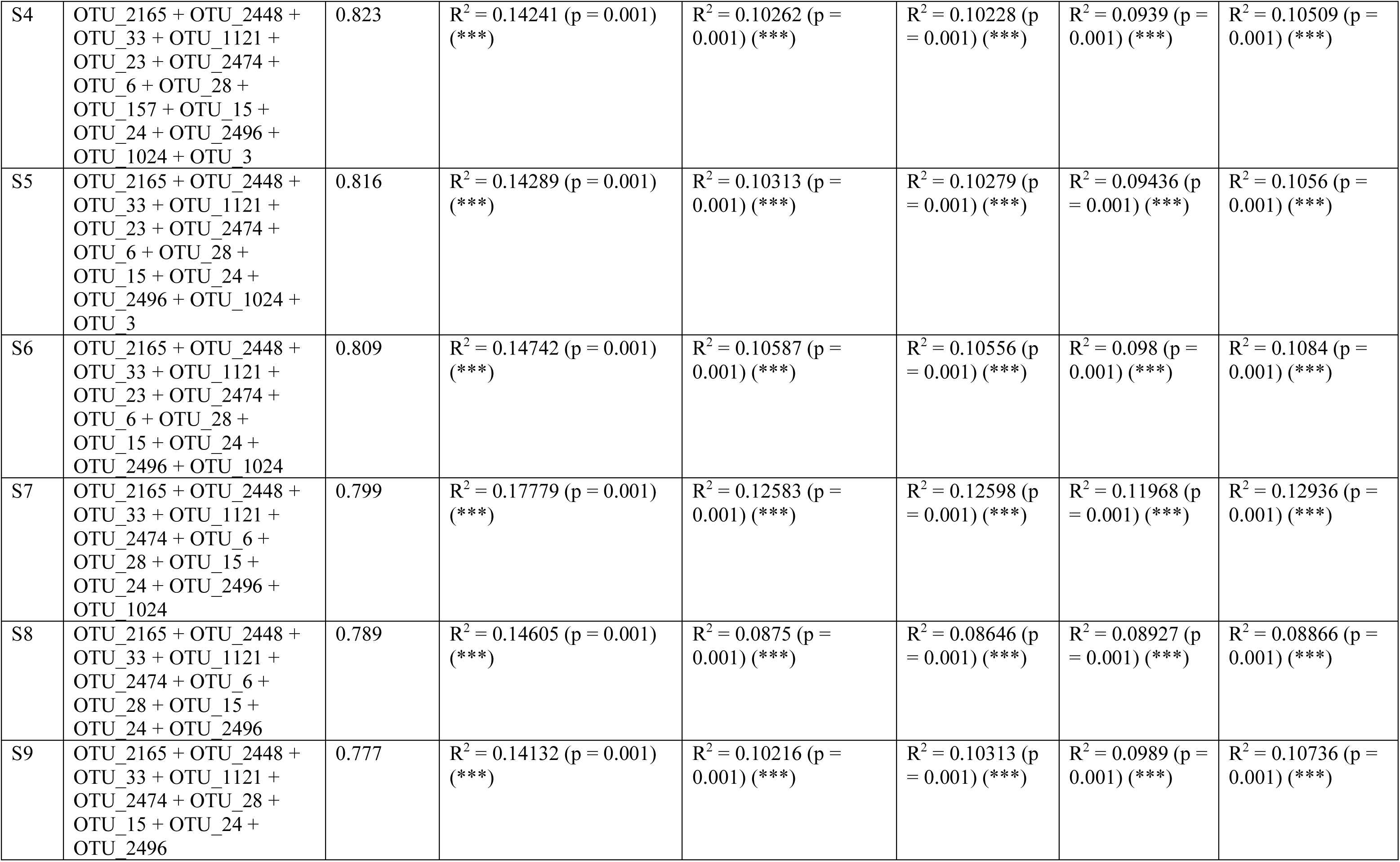
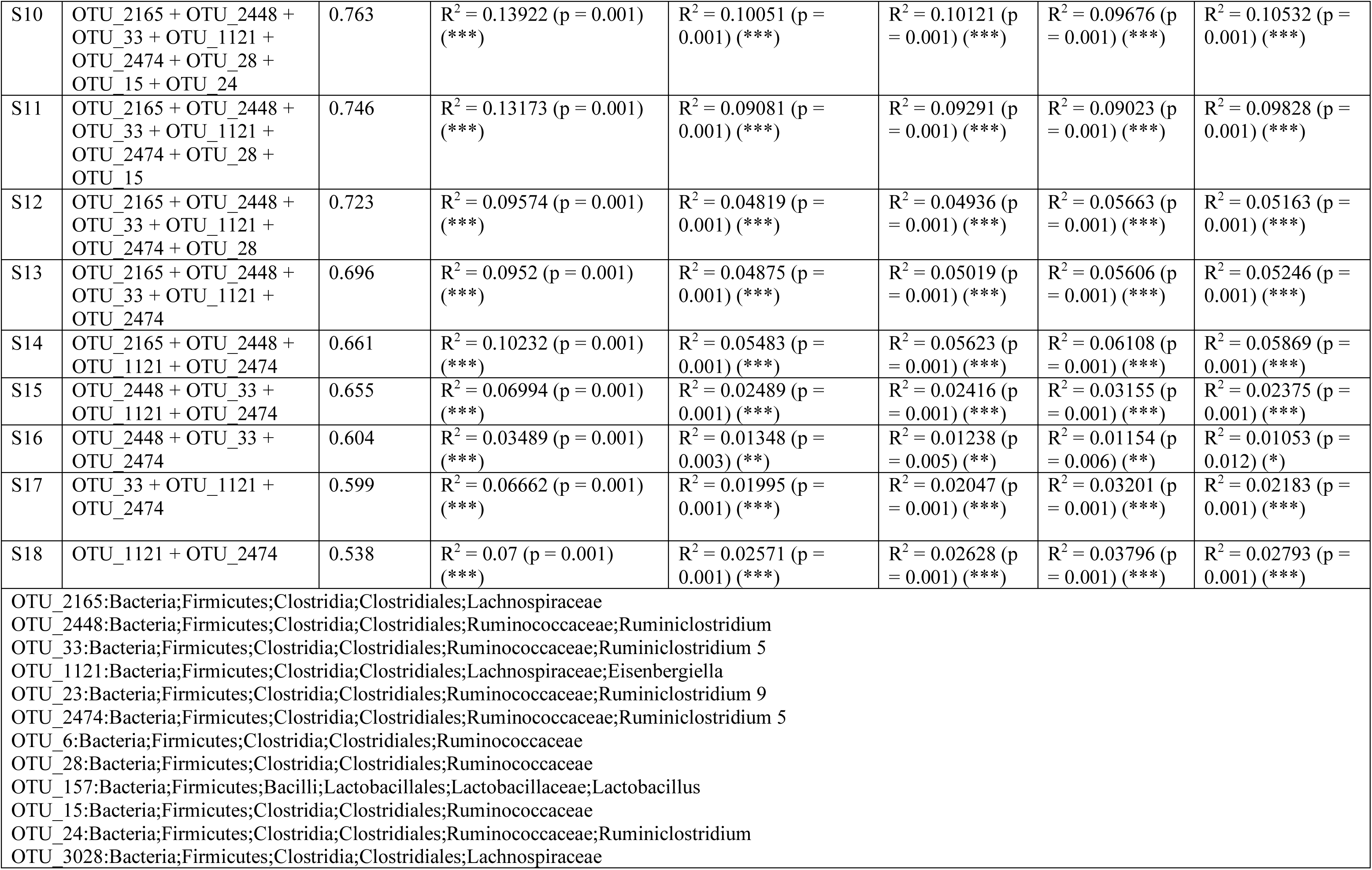
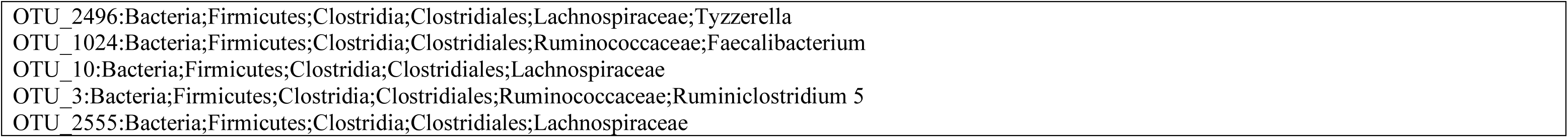
Subset analysis from BVSTEP routine listing top 18 subsets with highest correlation with the full OTU table considering Bray-Curtis distance done on weekly basis. For each subset, PERMANOVA was performed against different sources of variations.

| Subsets of top 1000 most abundant OTUs |  | Correlation with full OTU table (R) | PERMANOVA (full OTU table) |  |  |  |  |
| --- | --- | --- | --- | --- | --- | --- | --- |
|  |  |  | Groups | BW_Mean | FI | FCR | Gain |
| | | | $R^2 = 0.16763$ (p = 0.001) (***) | $R^2 = 0.11721$ (p = 0.001) (***) | $R^2 = 0.11856$ (p = 0.001) (***) | $R^2 = 0.1086$ (p = 0.001) (***) | $R^2 = 0.11886$ (p = 0.001) (***) |
|  |  |  | PERMANOVA (subsets) |  |  |  |  |
|  |  |  | Groups | BW_Mean | FI | FCR | Gain |
| S1 | OTU_2165 + OTU_2448 + OTU_33 + OTU_1121 + OTU_23 + OTU_2474 + OTU_6 + OTU_28 + OTU_157 + OTU_15 + OTU_24 + OTU_3028 + OTU_2496 + OTU_1024 + OTU_10 + OTU_3 + OTU_2555 | 0.833 | $R^2 = 0.14768$ (p = 0.001) (***) | $R^2 = 0.10732$ (p = 0.001) (***) | $R^2 = 0.10784$ (p = 0.001) (***) | $R^2 = 0.10117$ (p = 0.001) (***) | $R^2 = 0.11143$ (p = 0.001) (***) |
| S2 | OTU_2165 + OTU_2448 + OTU_33 + OTU_1121 + OTU_23 + OTU_2474 + OTU_6 + OTU_28 + OTU_157 + OTU_15 + OTU_24 + OTU_3028 + OTU_2496 + OTU_1024 + OTU_3 + OTU_2555 | 0.83 | $R^2 = 0.13977$ (p = 0.001) (***) | $R^2 = 0.10028$ (p = 0.001) (***) | $R^2 = 0.09987$ (p = 0.001) (***) | $R^2 = 0.09203$ (p = 0.001) (***) | $R^2 = 0.1025$ (p = 0.001) (***) |
| S3 | OTU_2165 + OTU_2448 + OTU_33 + OTU_1121 + OTU_23 + OTU_2474 + OTU_6 + OTU_28 + OTU_157 + OTU_15 + OTU_24 + OTU_3028 + OTU_2496 + OTU_1024 + OTU_3 | 0.827 | $R^2 = 0.14186$ (p = 0.001) (***) | $R^2 = 0.10205$ (p = 0.001) (***) | $R^2 = 0.10165$ (p = 0.001) (***) | $R^2 = 0.09344$ (p = 0.001) (***) | $R^2 = 0.10435$ (p = 0.001) (***) |
| S4 | OTU_2165 + OTU_2448 + OTU_33 + OTU_1121 + OTU_23 + OTU_2474 + OTU_6 + OTU_28 + OTU_157 + OTU_15 + OTU_24 + OTU_2496 + OTU_1024 + OTU_3 | 0.823 | $R^2 = 0.14241$ (p = 0.001) (***) | $R^2 = 0.10262$ (p = 0.001) (***) | $R^2 = 0.10228$ (p = 0.001) (***) | $R^2 = 0.0939$ (p = 0.001) (***) | $R^2 = 0.10509$ (p = 0.001) (***) |
| S5 | OTU_2165 + OTU_2448 + OTU_33 + OTU_1121 + OTU_23 + OTU_2474 + OTU_6 + OTU_28 + OTU_15 + OTU_24 + OTU_2496 + OTU_1024 + OTU_3 | 0.816 | $R^2 = 0.14289$ (p = 0.001) (***) | $R^2 = 0.10313$ (p = 0.001) (***) | $R^2 = 0.10279$ (p = 0.001) (***) | $R^2 = 0.09436$ (p = 0.001) (***) | $R^2 = 0.1056$ (p = 0.001) (***) |
| S6 | OTU_2165 + OTU_2448 + OTU_33 + OTU_1121 + OTU_23 + OTU_2474 + OTU_6 + OTU_28 + OTU_15 + OTU_24 + OTU_2496 + OTU_1024 | 0.809 | $R^2 = 0.14742$ (p = 0.001) (***) | $R^2 = 0.10587$ (p = 0.001) (***) | $R^2 = 0.10556$ (p = 0.001) (***) | $R^2 = 0.098$ (p = 0.001) (***) | $R^2 = 0.1084$ (p = 0.001) (***) |
| S7 | OTU_2165 + OTU_2448 + OTU_33 + OTU_1121 + OTU_2474 + OTU_6 + OTU_28 + OTU_15 + OTU_24 + OTU_2496 + OTU_1024 | 0.799 | $R^2 = 0.17779$ (p = 0.001) (***) | $R^2 = 0.12583$ (p = 0.001) (***) | $R^2 = 0.12598$ (p = 0.001) (***) | $R^2 = 0.11968$ (p = 0.001) (***) | $R^2 = 0.12936$ (p = 0.001) (***) |
| S8 | OTU_2165 + OTU_2448 + OTU_33 + OTU_1121 + OTU_2474 + OTU_6 + OTU_28 + OTU_15 + OTU_24 + OTU_2496 | 0.789 | $R^2 = 0.14605$ (p = 0.001) (***) | $R^2 = 0.0875$ (p = 0.001) (***) | $R^2 = 0.08646$ (p = 0.001) (***) | $R^2 = 0.08927$ (p = 0.001) (***) | $R^2 = 0.08866$ (p = 0.001) (***) |
| S9 | OTU_2165 + OTU_2448 + OTU_33 + OTU_1121 + OTU_2474 + OTU_28 + OTU_15 + OTU_24 + OTU_2496 | 0.777 | $R^2 = 0.14132$ (p = 0.001) (***) | $R^2 = 0.10216$ (p = 0.001) (***) | $R^2 = 0.10313$ (p = 0.001) (***) | $R^2 = 0.0989$ (p = 0.001) (***) | $R^2 = 0.10736$ (p = 0.001) (***) |
| S10 | OTU_2165 + OTU_2448 + OTU_33 + OTU_1121 + OTU_2474 + OTU_28 + OTU_15 + OTU_24 | 0.763 | $R^2 = 0.13922$ (p = 0.001) (***) | $R^2 = 0.10051$ (p = 0.001) (***) | $R^2 = 0.10121$ (p = 0.001) (***) | $R^2 = 0.09676$ (p = 0.001) (***) | $R^2 = 0.10532$ (p = 0.001) (***) |
| S11 | OTU_2165 + OTU_2448 + OTU_33 + OTU_1121 + OTU_2474 + OTU_28 + OTU_15 | 0.746 | $R^2 = 0.13173$ (p = 0.001) (***) | $R^2 = 0.09081$ (p = 0.001) (***) | $R^2 = 0.09291$ (p = 0.001) (***) | $R^2 = 0.09023$ (p = 0.001) (***) | $R^2 = 0.09828$ (p = 0.001) (***) |
| S12 | OTU_2165 + OTU_2448 + OTU_33 + OTU_1121 + OTU_2474 + OTU_28 | 0.723 | $R^2 = 0.09574$ (p = 0.001) (***) | $R^2 = 0.04819$ (p = 0.001) (***) | $R^2 = 0.04936$ (p = 0.001) (***) | $R^2 = 0.05663$ (p = 0.001) (***) | $R^2 = 0.05163$ (p = 0.001) (***) |
| S13 | OTU_2165 + OTU_2448 + OTU_33 + OTU_1121 + OTU_2474 | 0.696 | $R^2 = 0.0952$ (p = 0.001) (***) | $R^2 = 0.04875$ (p = 0.001) (***) | $R^2 = 0.05019$ (p = 0.001) (***) | $R^2 = 0.05606$ (p = 0.001) (***) | $R^2 = 0.05246$ (p = 0.001) (***) |
| S14 | OTU_2165 + OTU_2448 + OTU_1121 + OTU_2474 | 0.661 | $R^2 = 0.10232$ (p = 0.001) (***) | $R^2 = 0.05483$ (p = 0.001) (***) | $R^2 = 0.05623$ (p = 0.001) (***) | $R^2 = 0.06108$ (p = 0.001) (***) | $R^2 = 0.05869$ (p = 0.001) (***) |
| S15 | OTU_2448 + OTU_33 + OTU_1121 + OTU_2474 | 0.655 | $R^2 = 0.06994$ (p = 0.001) (***) | $R^2 = 0.02489$ (p = 0.001) (***) | $R^2 = 0.02416$ (p = 0.001) (***) | $R^2 = 0.03155$ (p = 0.001) (***) | $R^2 = 0.02375$ (p = 0.001) (***) |
| S16 | OTU_2448 + OTU_33 + OTU_2474 | 0.604 | $R^2 = 0.03489$ (p = 0.001) (***) | $R^2 = 0.01348$ (p = 0.003) (**) | $R^2 = 0.01238$ (p = 0.005) (**) | $R^2 = 0.01154$ (p = 0.006) (**) | $R^2 = 0.01053$ (p = 0.012) (*) |
| S17 | OTU_33 + OTU_1121 + OTU_2474 | 0.599 | $R^2 = 0.06662$ (p = 0.001) (***) | $R^2 = 0.01995$ (p = 0.001) (***) | $R^2 = 0.02047$ (p = 0.001) (***) | $R^2 = 0.03201$ (p = 0.001) (***) | $R^2 = 0.02183$ (p = 0.001) (***) |
| S18 | OTU_1121 + OTU_2474 | 0.538 | $R^2 = 0.07$ (p = 0.001) (***) | $R^2 = 0.02571$ (p = 0.001) (***) | $R^2 = 0.02628$ (p = 0.001) (***) | $R^2 = 0.03796$ (p = 0.001) (***) | $R^2 = 0.02793$ (p = 0.001) (***) |
| OTU_2165: Bacteria; Firmicutes; Clostridia; Clostridiales; Lachnospiraceae<br>OTU_2448: Bacteria; Firmicutes; Clostridia; Clostridiales; Ruminococcaceae; Ruminiclostridium<br>OTU_33: Bacteria; Firmicutes; Clostridia; Clostridiales; Ruminococcaceae; Ruminiclostridium 5<br>OTU_1121: Bacteria; Firmicutes; Clostridia; Clostridiales; Lachnospiraceae; Eisenbergiella<br>OTU_23: Bacteria; Firmicutes; Clostridia; Clostridiales; Ruminococcaceae; Ruminiclostridium 9<br>OTU_2474: Bacteria; Firmicutes; Clostridia; Clostridiales; Ruminococcaceae; Ruminiclostridium 5<br>OTU_6: Bacteria; Firmicutes; Clostridia; Clostridiales; Ruminococcaceae<br>OTU_28: Bacteria; Firmicutes; Clostridia; Clostridiales; Ruminococcaceae<br>OTU_157: Bacteria; Firmicutes; Bacilli; Lactobacillales; Lactobacillaceae; Lactobacillus<br>OTU_15: Bacteria; Firmicutes; Clostridia; Clostridiales; Ruminococcaceae<br>OTU_24: Bacteria; Firmicutes; Clostridia; Clostridiales; Ruminococcaceae; Ruminiclostridium<br>OTU_3028: Bacteria; Firmicutes; Clostridia; Clostridiales; Lachnospiraceae |  |  |  |  |  |  |  |

|  |
| --- |
| OTU_2496: Bacteria; Firmicutes; Clostridia; Clostridiales; Lachnospiraceae; Tyzzerella |
| OTU_1024: Bacteria; Firmicutes; Clostridia; Clostridiales; Ruminococcaceae; Faecalibacterium |
| OTU_10: Bacteria; Firmicutes; Clostridia; Clostridiales; Lachnospiraceae |
| OTU_3: Bacteria; Firmicutes; Clostridia; Clostridiales; Ruminococcaceae; Ruminiclostridium 5 |
| OTU_2555: Bacteria; Firmicutes; Clostridia; Clostridiales; Lachnospiraceae |

**Figure 3:**
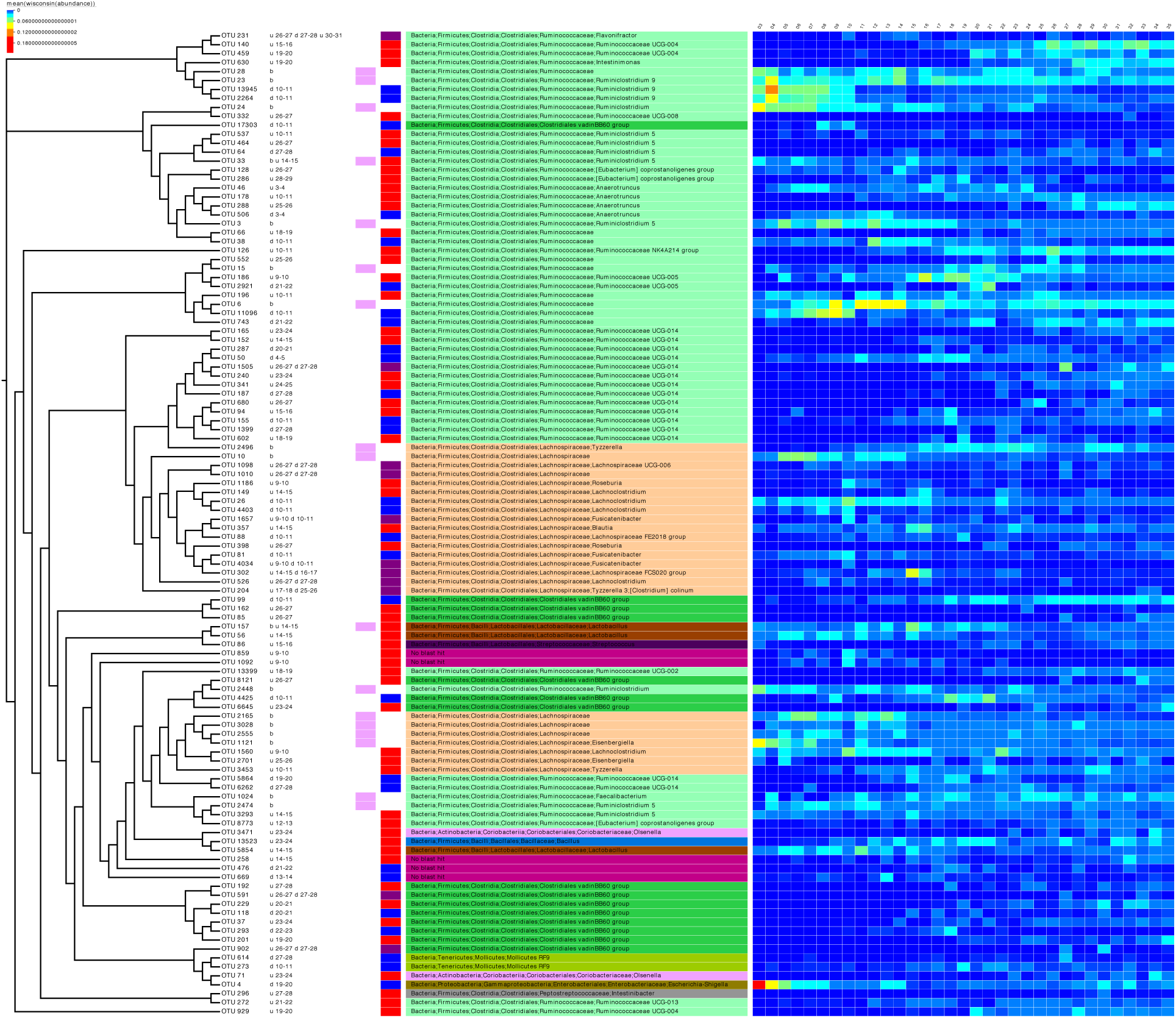
Phylogenetic tree of the subset of OTUs selected as significant on differential analysis (based on Table 3 and Supplementary Table 1). Next to the OTU labels are descriptive text representing where the OTUs were found to be significant, for example, the first entry for OTU 231, “u 26-27 d 27-28 u 30-31”, can be read as upregulated going from day 26 to 27 and then from day 30 to 31 and downregulated going from day 27 to 28. “b” represents the OTUs selected in the subset analysis. The next two columns are a pictorial representation of the above-mentioned descriptive text with pink colour representing OTUs selected in subset analysis, red colour for upregulated OTUs, blue for downregulated OTUs, and purple for OTUs which show the both trends (up/down regulation). The next column shows the taxonomy of the OTUs according to SILVA v123 with colouring at unique family level. The heatmap was drawn by collating the mean values of OTUs for samples from the same day after performing proportional standardization on the full OTU table using wisconsin() function.

## Discussion

Comprehensive investigation of the chicken cecal microbiome at a day-to-day level revealed a rapid increase in diversity up to day 12, with microbial variation observed both in terms of genera and abundance. We suspect this early variation is due to competitive factors determined by space and available food resources. Post day 20 there exists a considerable stabilisation of the chicken cecal microbiome where the relative microbial diversity and abundances are standardised, with environmental factors (in this case the host chicken) exerting a greater influence on any change in the microbial diversity. Between days 12 and 20 we observe a shift from competitive to environmental drivers of microbial community creating a window of opportunity whereby *Campylobacter* appears. We identified *Campylobacter* at day 16 with this day lying immediately after the most substantial changes in metabolic profiles observed over the whole period. Whilst we identified *Campylobacter* within 25% of the pens on day 16, we would naturally expect *Campylobacter* to spread to other chickens and pens and also be identified on subsequent days. We suspect that the experimental set-up here was such that following random selection of birds from each pen on each day, sacrificing the bird (to perform gDNA extraction from the ceca) did not allow for an opportunity for *Campylobacter* to spread to other chickens or pens. Clearly in a typical farm set-up this would not be the case and *Campylobacter* would spread naturally.

Microbial variation over time is most likely influenced by diet of the chickens whereby significant shifts in OTU abundances and beta dispersion of the samples often corresponded with changes in feed. Notably, the relatively high proportion of OTUs that were log2 fold different between days 9 and 11, and days 26 and 28, and beta dispersion for days 11-13 corresponded with changes in feed from grower to finisher. Further studies investigating different feed content is required to ascertain the complete impact on chicken cecal microbiome.

Previous microbiome studies of chicken ceca have often lacked the day-to-day sampling points, replicate numbers, relevant metadata and have often provided external *Campylobacter* infection that may potentially perturb the natural habitat. These have not allowed us to fully appreciate what is occurring in a natural environment in relation to how and when *Campylobacter* appears within the chicken gut. Thus, we believe the major strength of this study is that we achieve these missing points as we have performed the most comprehensive analysis of the chicken cecal microbiome to date, sampling from days 3 to 35, with 12 replicates per day (n=396), correlating additional metadata such as chicken weight and feed conversion rates and with *Campylobacter* detection in a natural environmental setting giving the most comparable experimental design to a farm set-up. As we were not able to sample the same chicken for all time points, future studies should investigate this further with added dietary information than what we have considered here, with experimental designs also to investigate and confirm the predicted pathways.

## Conclusions

Industry has endeavoured to reduce the burden of *Campylobacter* within chicken production lines with supplements often administered with the aim of performance enhancing and/or reducing bacteria such as *Campylobacter*, typically post day 25. The relative stability of the chicken cecal microbiota at this time point may explain the efficacy of such products, however the identification of a window of opportunity for bacteria such as *Campylobacter* may call for intervention strategies between days 12 to 20, or even earlier. This study can act as a baseline for future intervention strategies and help reduce the burden of *Campylobacter* within chickens.

## Availability of supporting data

The raw sequence files supporting the results of this article are available in the European Nucleotide Archive under the project accession number PRJEB25776.

## Author contributions

AM, AR, UL, BW, ND, NC and OG contributed to the study design. ND, NC and OG managed the study. CK, AM. AS and ML performed the sample collection and DNA extraction. LS, AM, AE and OG performed the library preparation and Illumina MiSeq sequencing at the LSHTM. UZI wrote the analysis scripts to generate the figures and tables in this paper. UZI and OG performed the bioinformatics and statistical analysis. UZI, ND, NC and OG drafted the initial version of the manuscript with all authors contributed to redrafting.

## Acknowledgements

The authors acknowledge research funding from Moy Park. UZI is funded by NERC Independent Research Fellowship (NE/L011956/1).

---

**Supplementary Figure 1:**
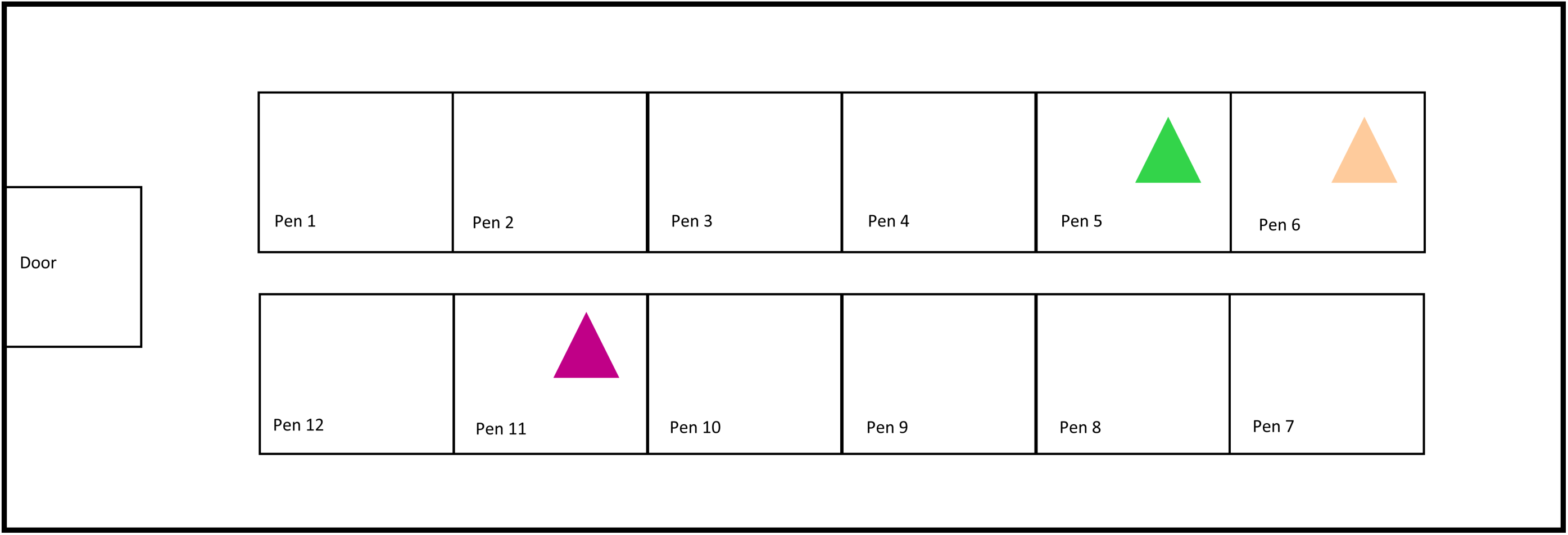
Spatial arrangement of pens. Triangles indicate pens with *Campylobacter*.

**Supplementary Figure 2:**
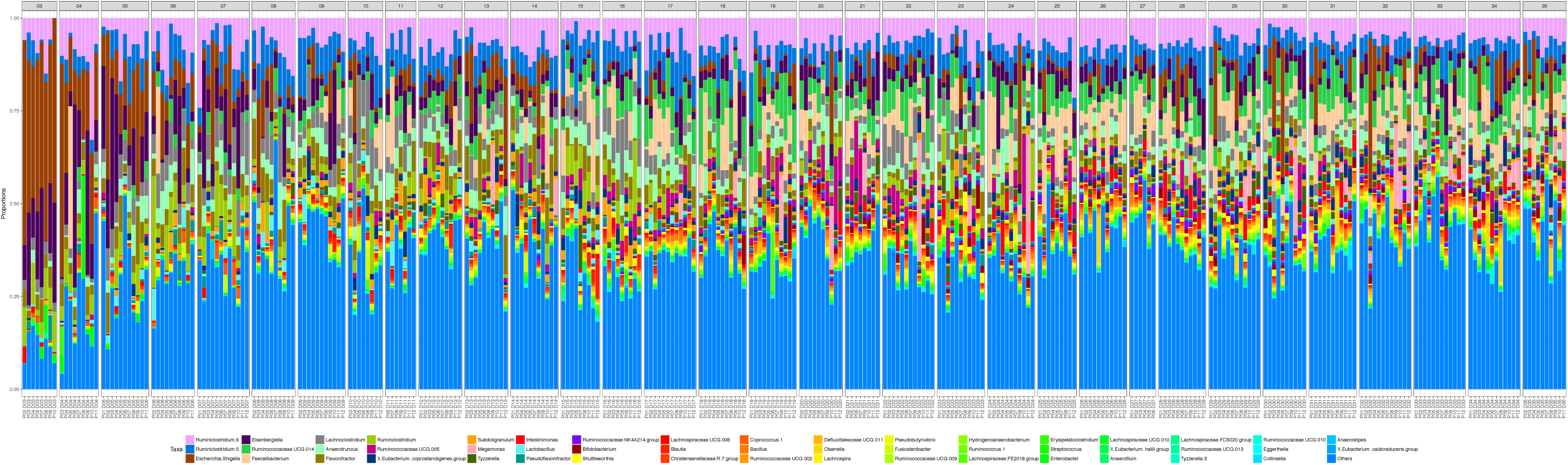
Relative abundance of 50 most abundant genera in this study.

**Supplementary Table 1:**
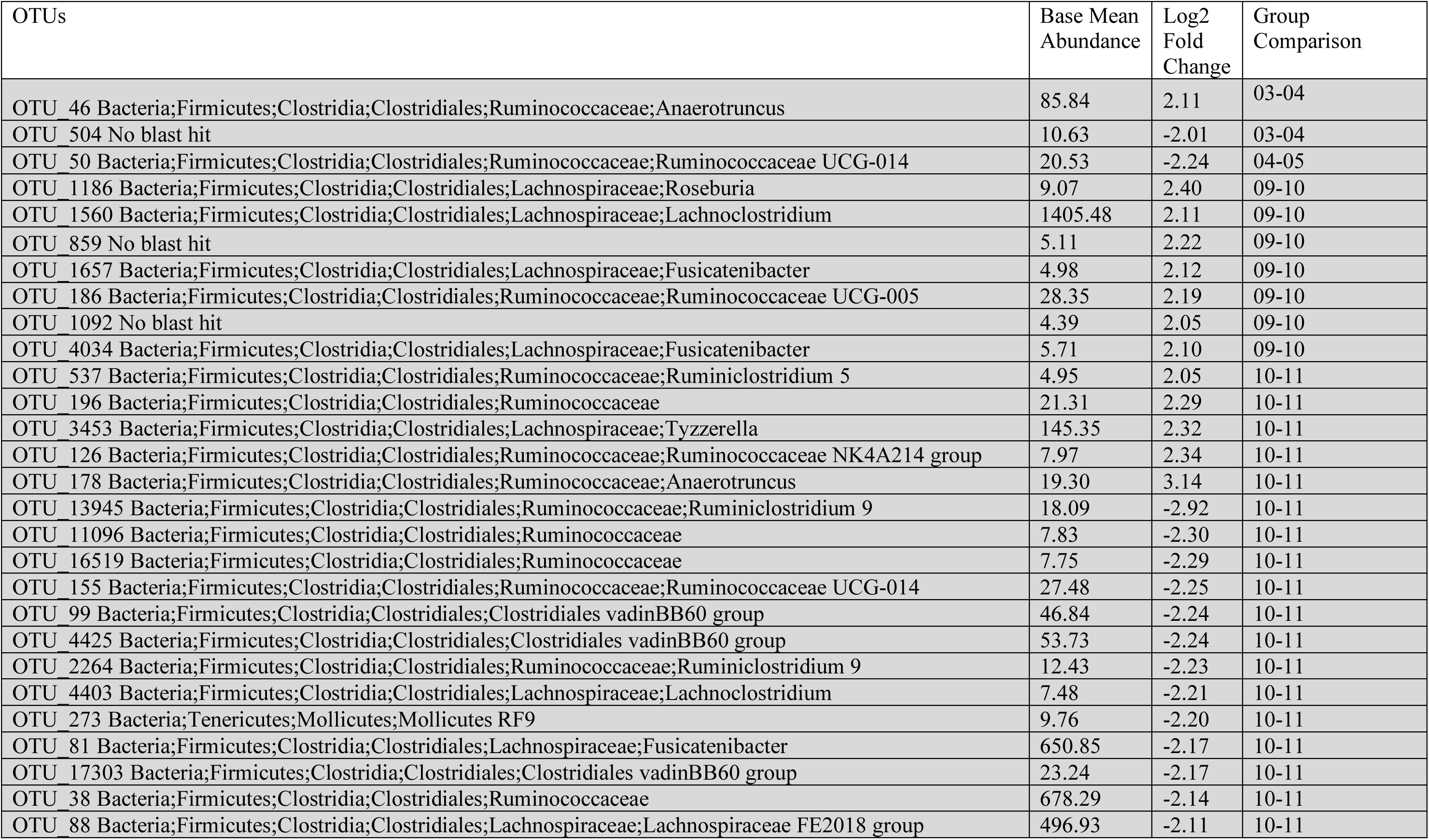
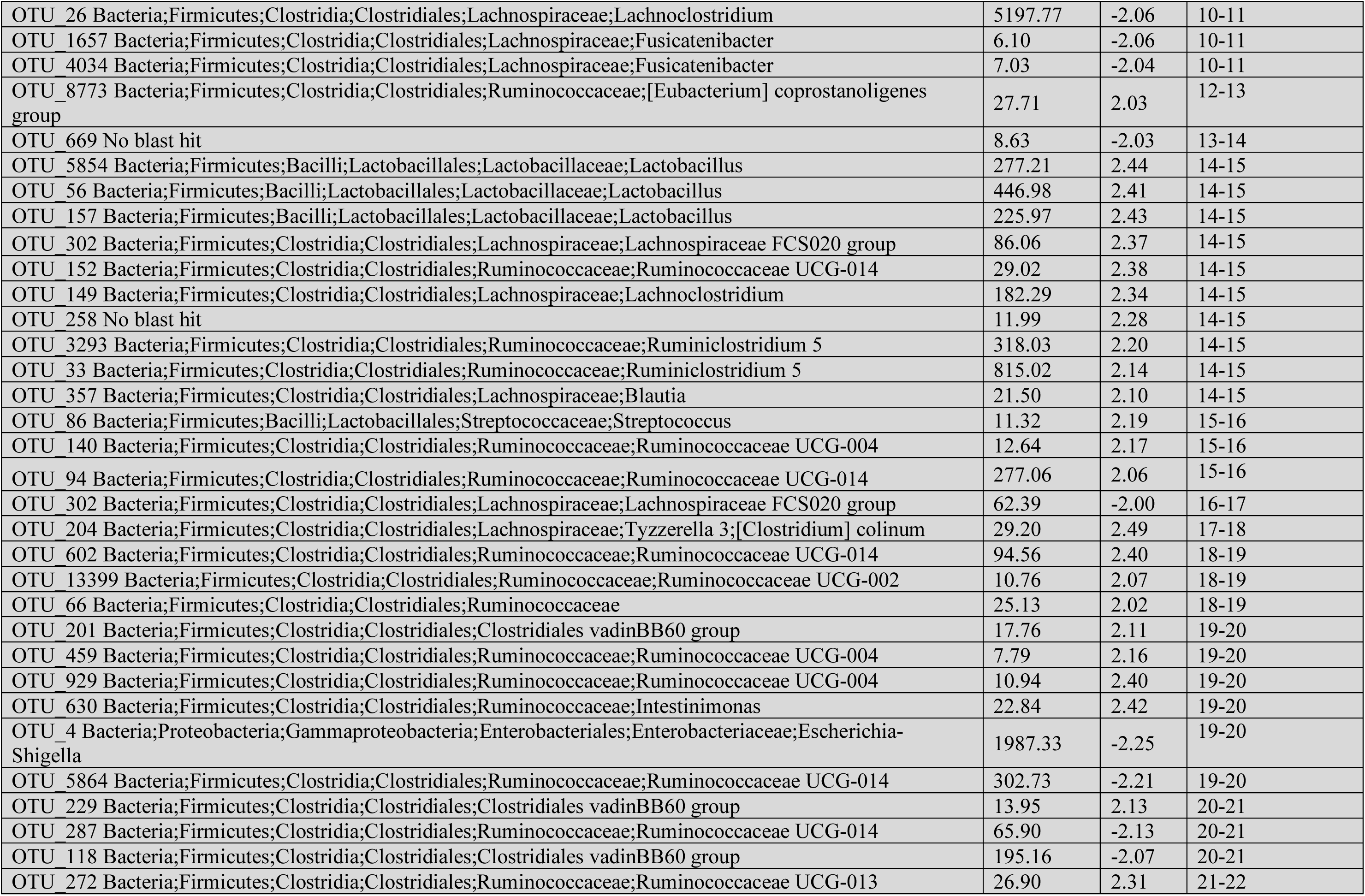
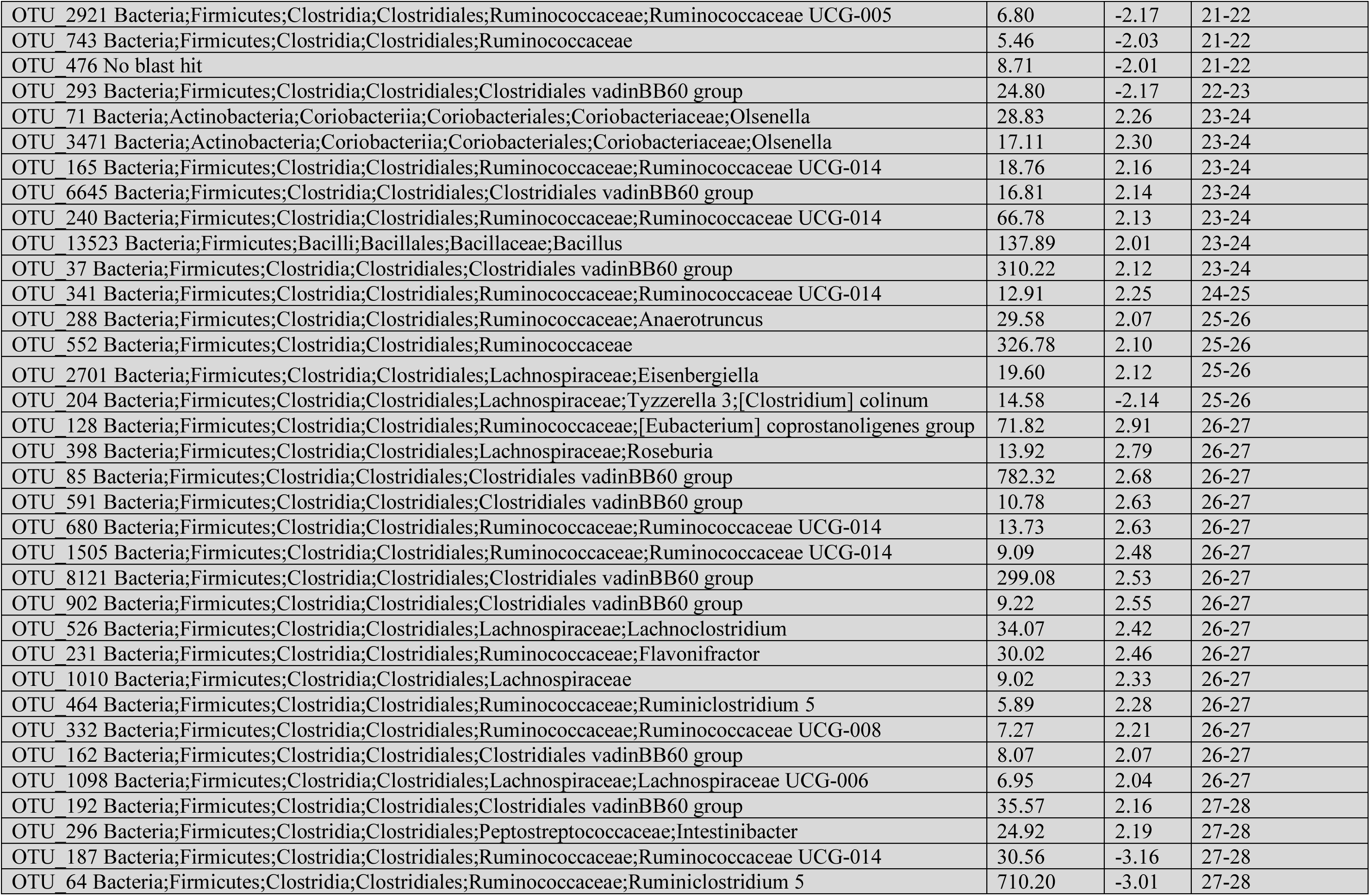
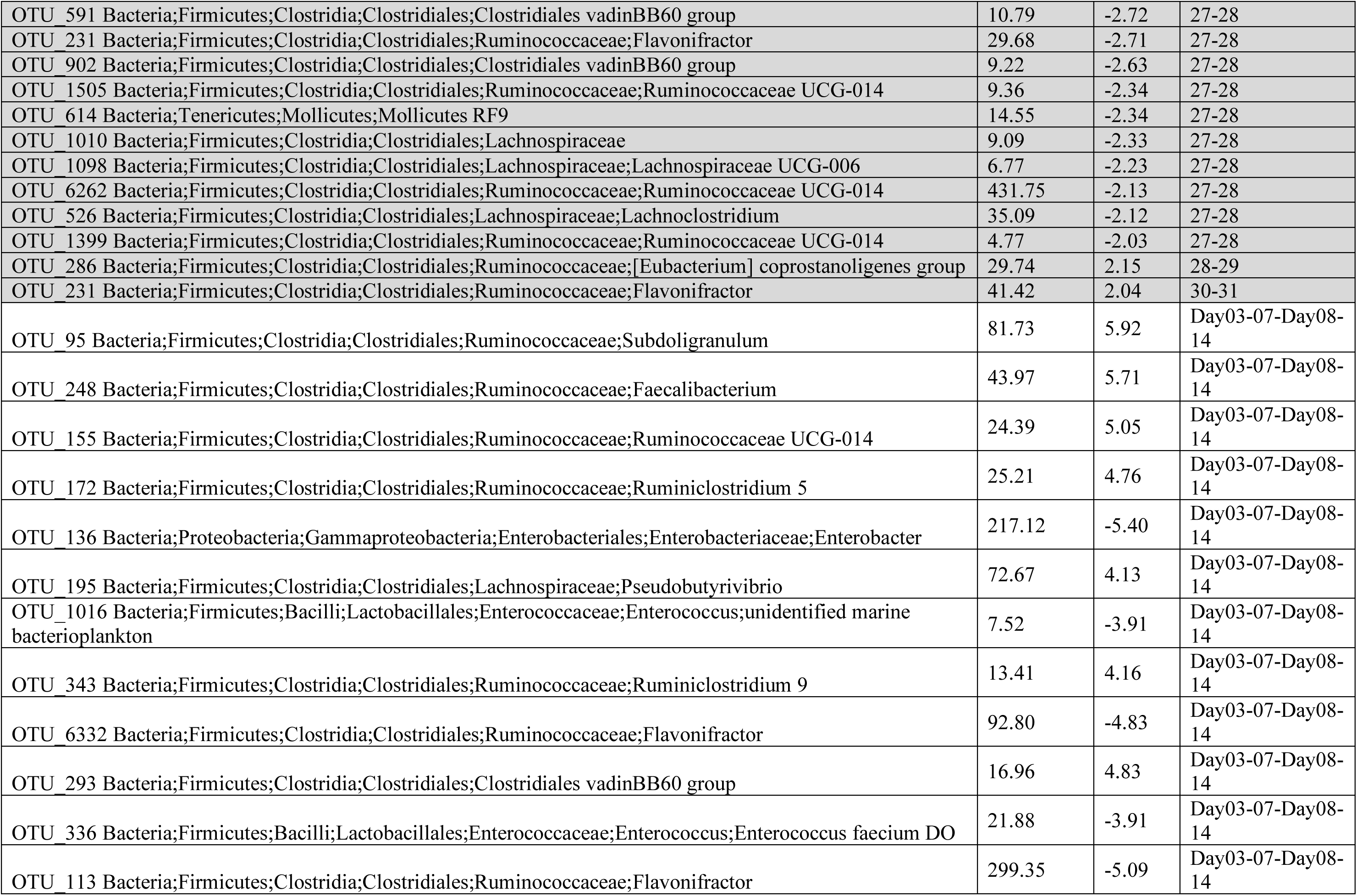
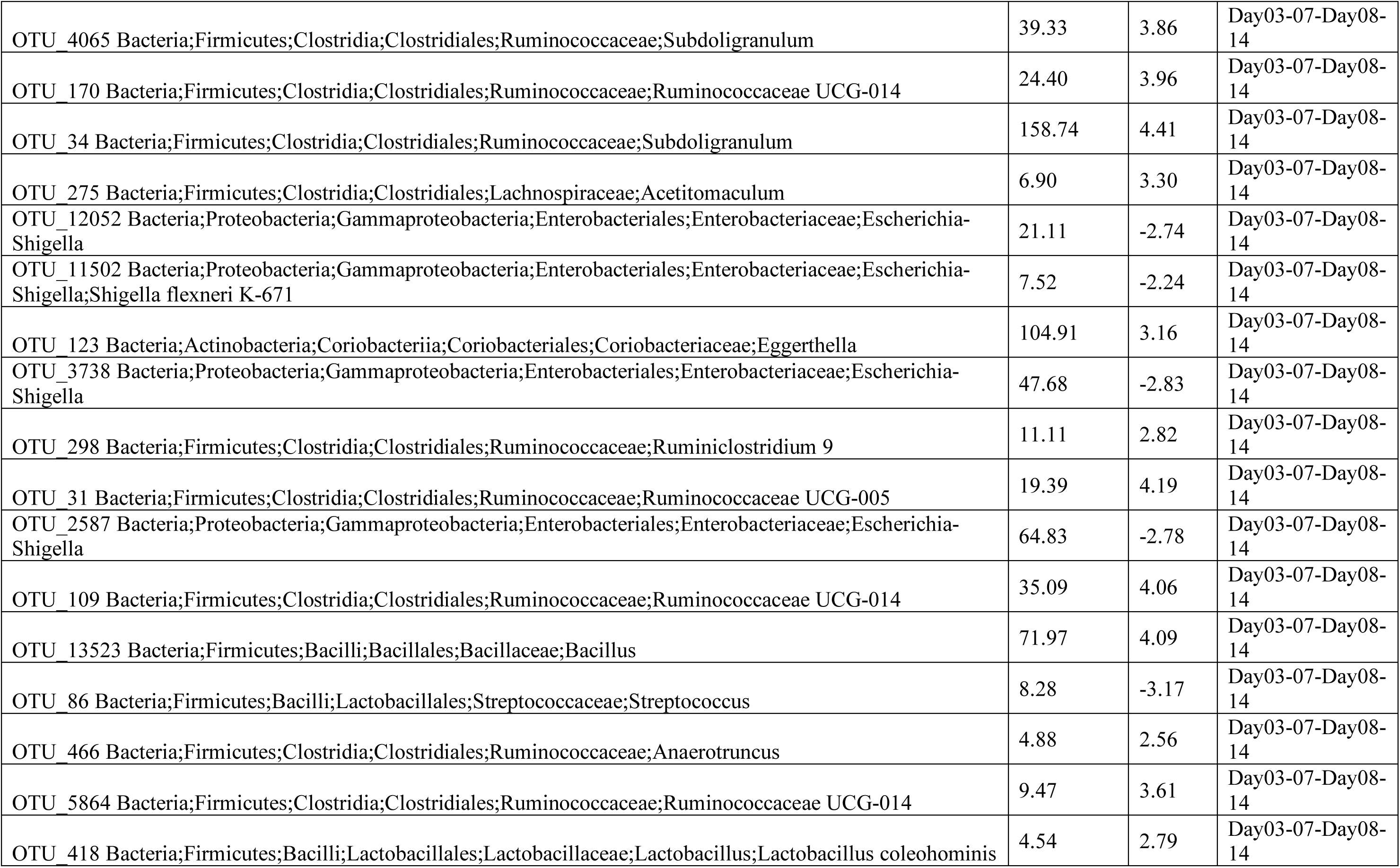
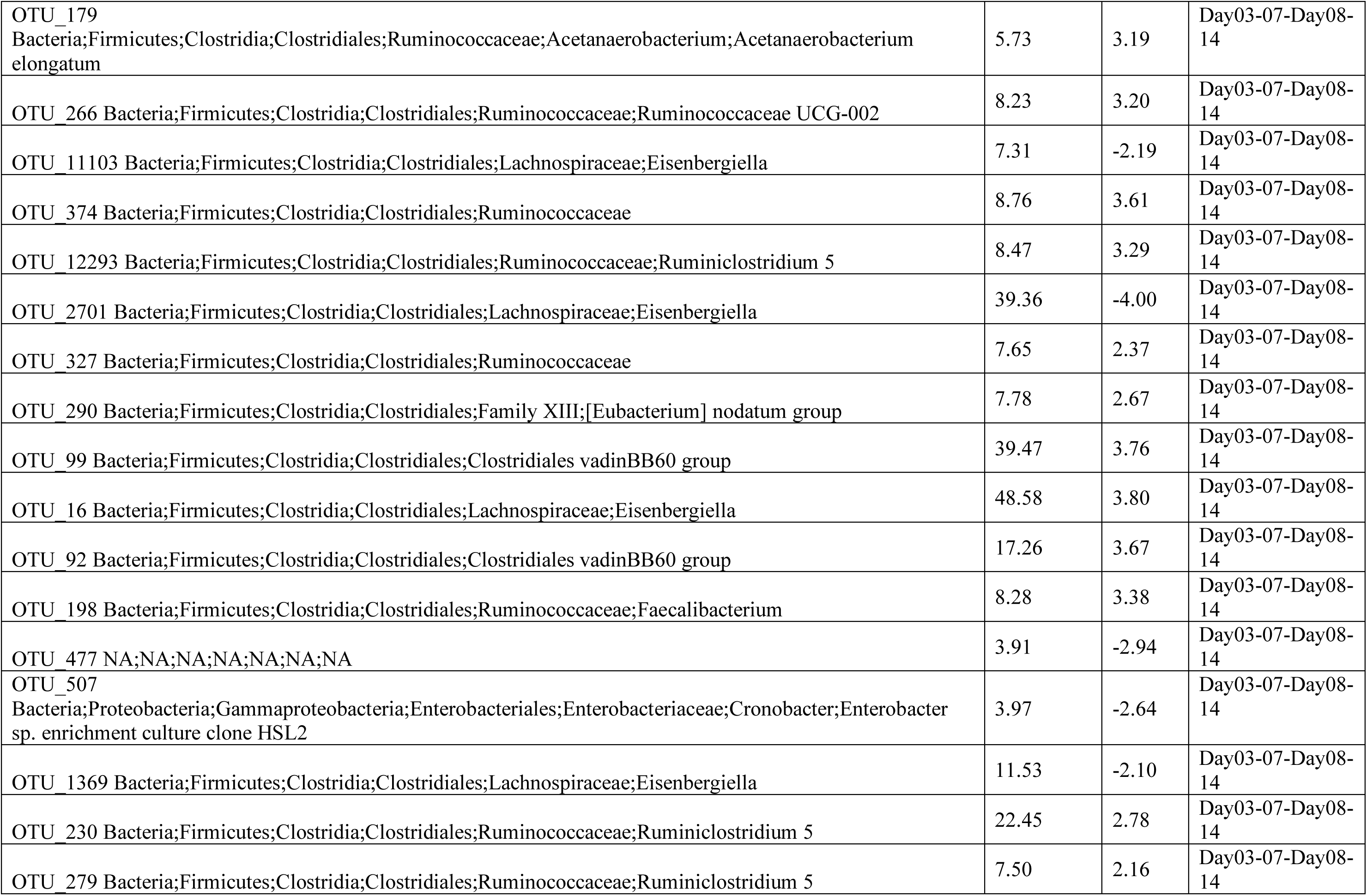
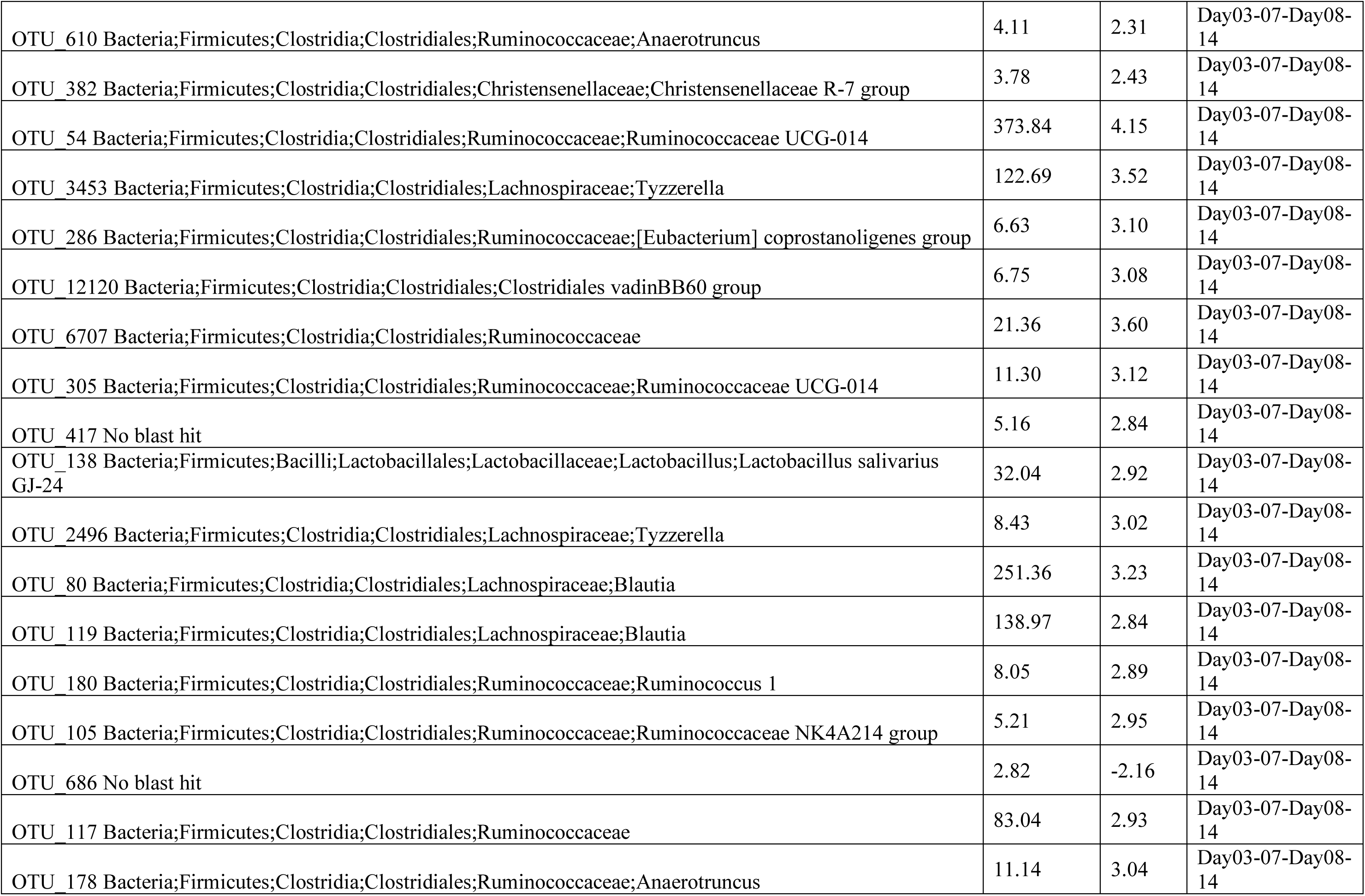
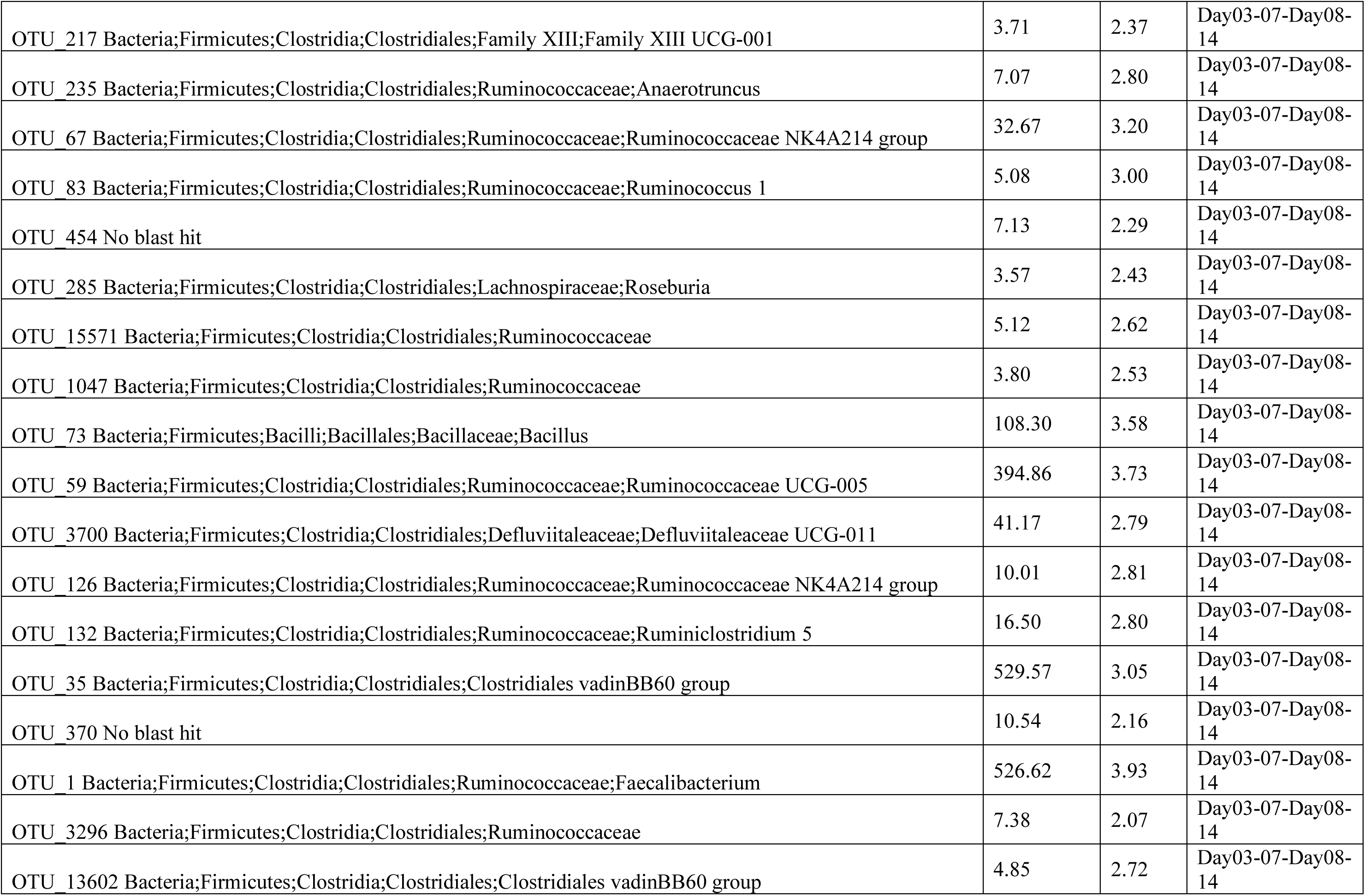
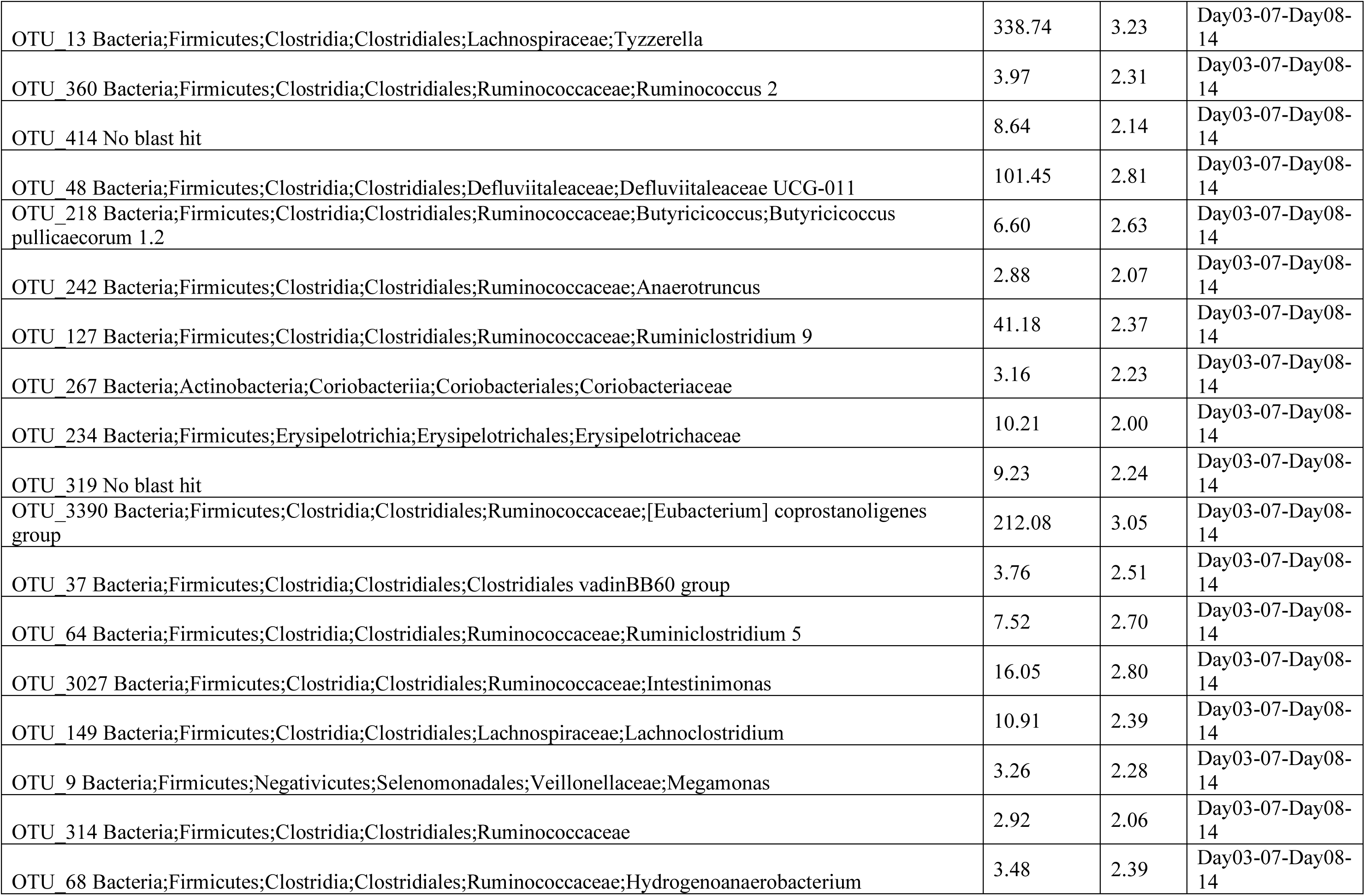
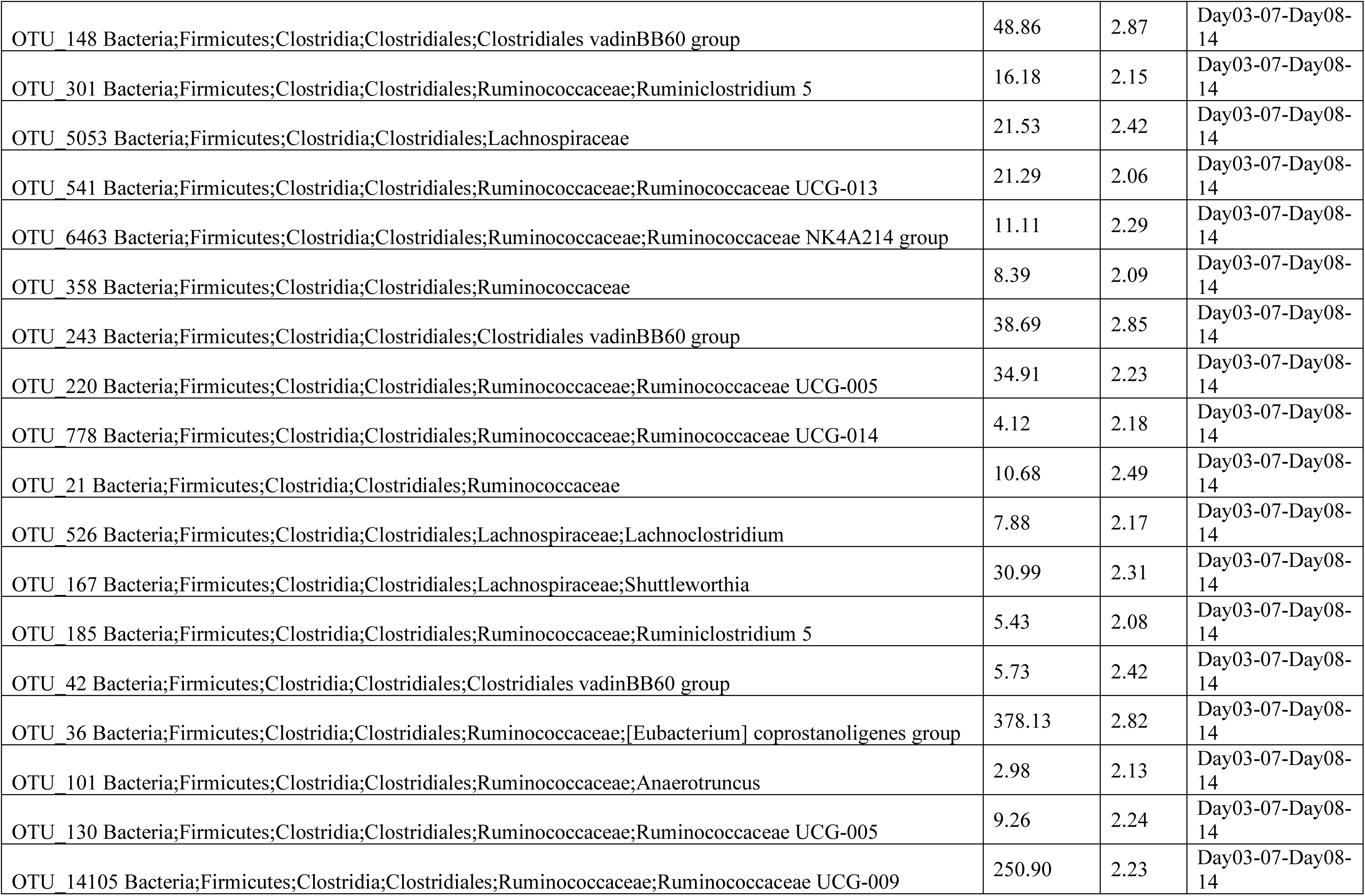
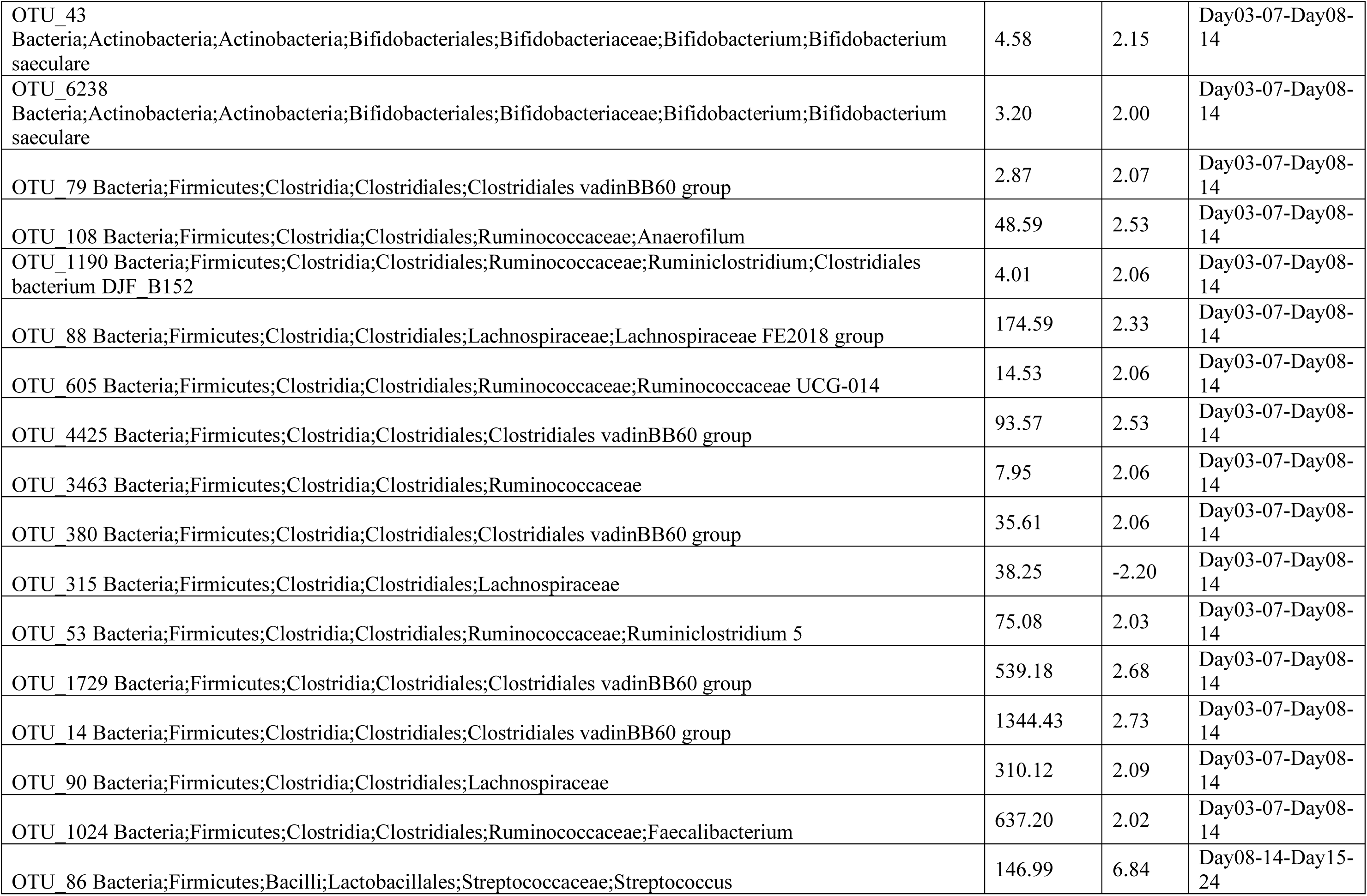
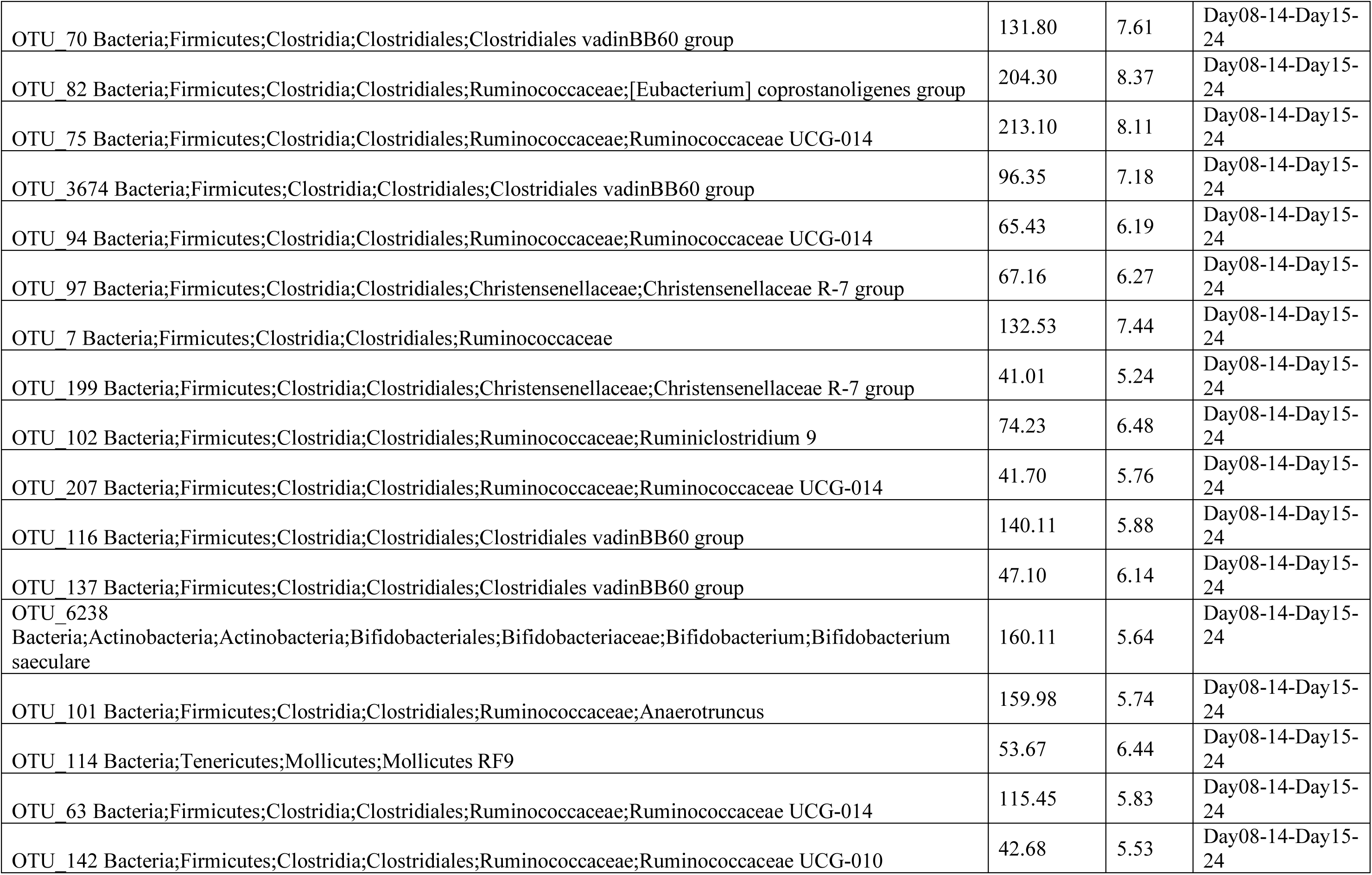
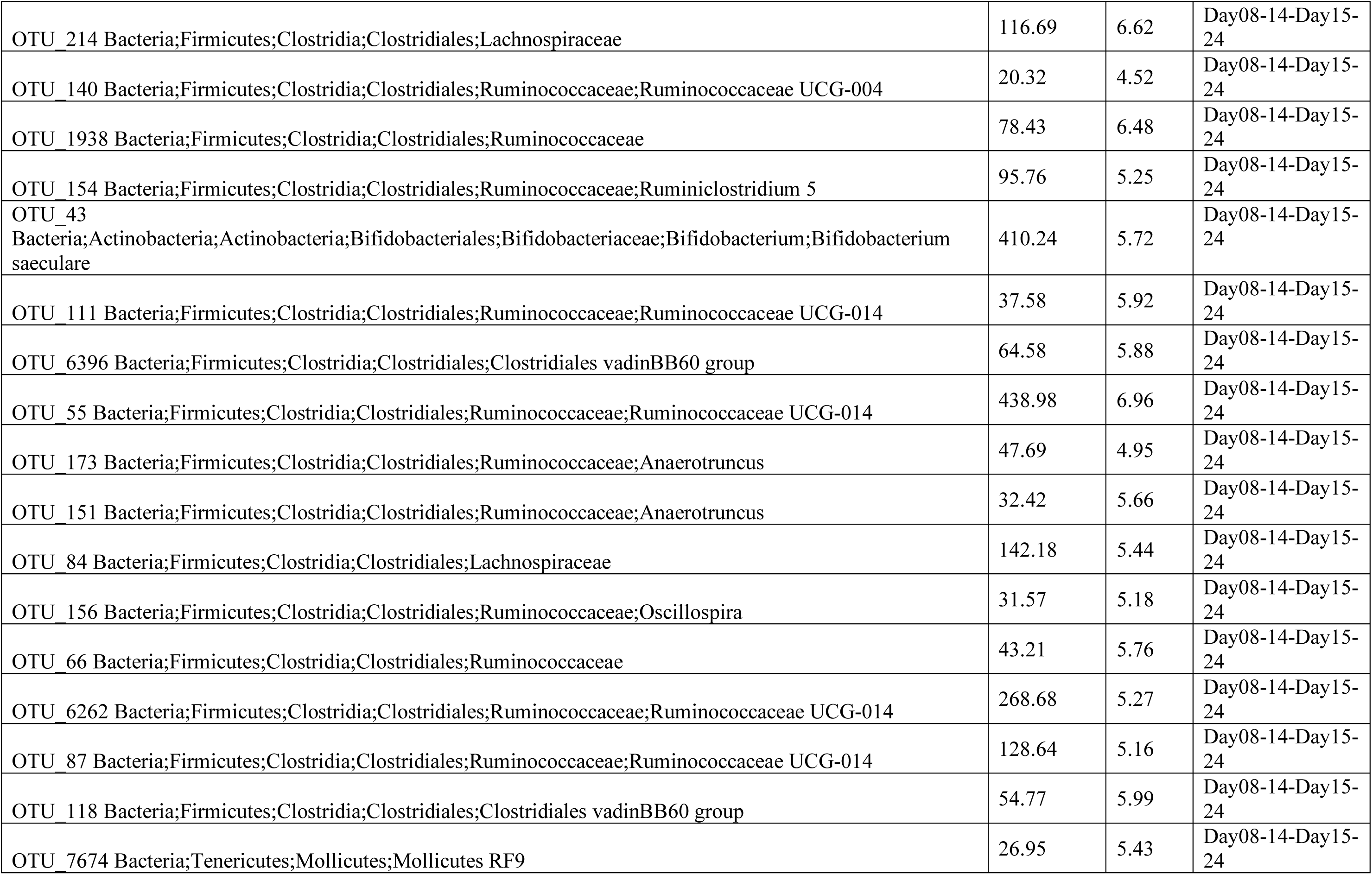
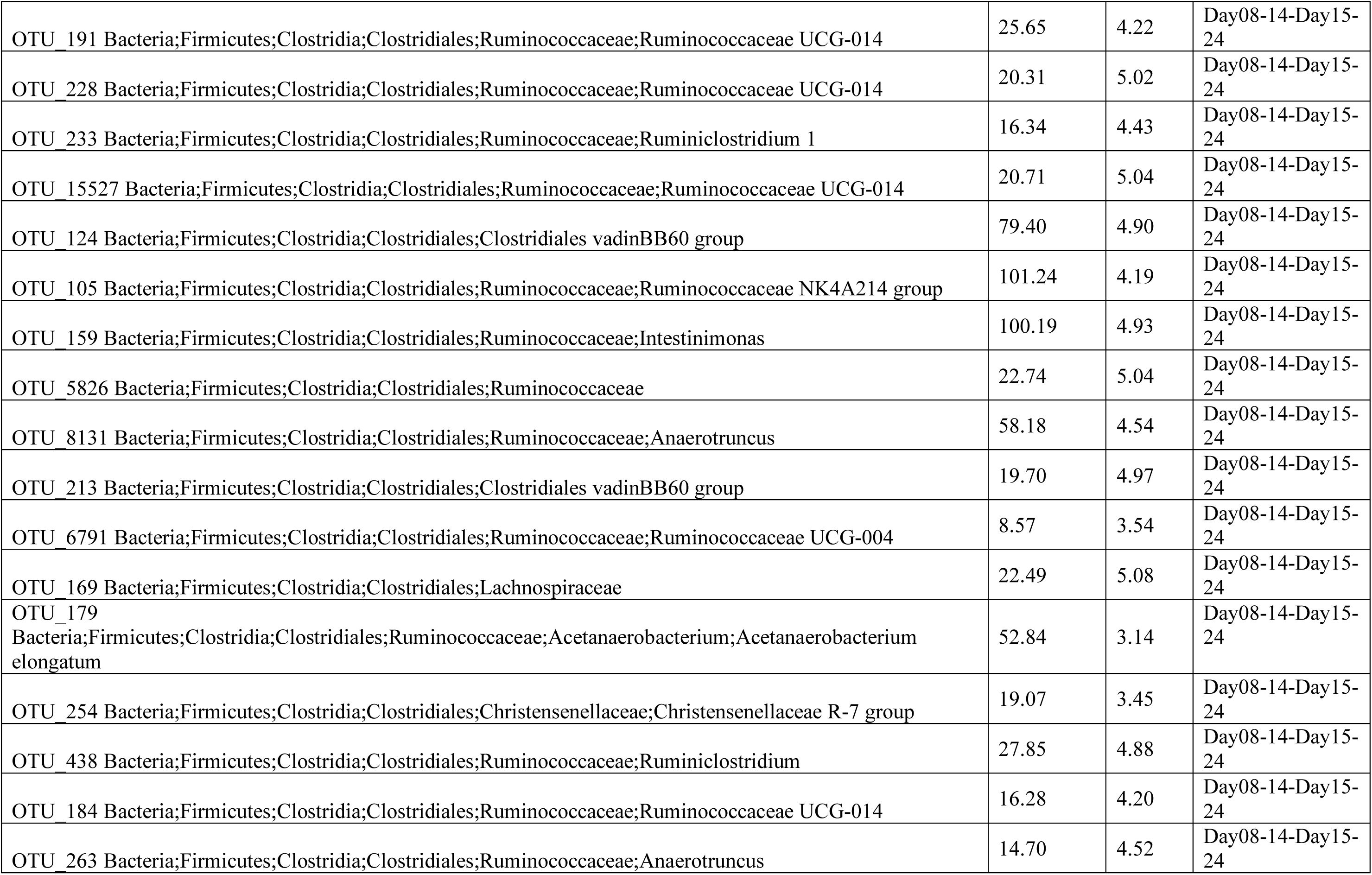
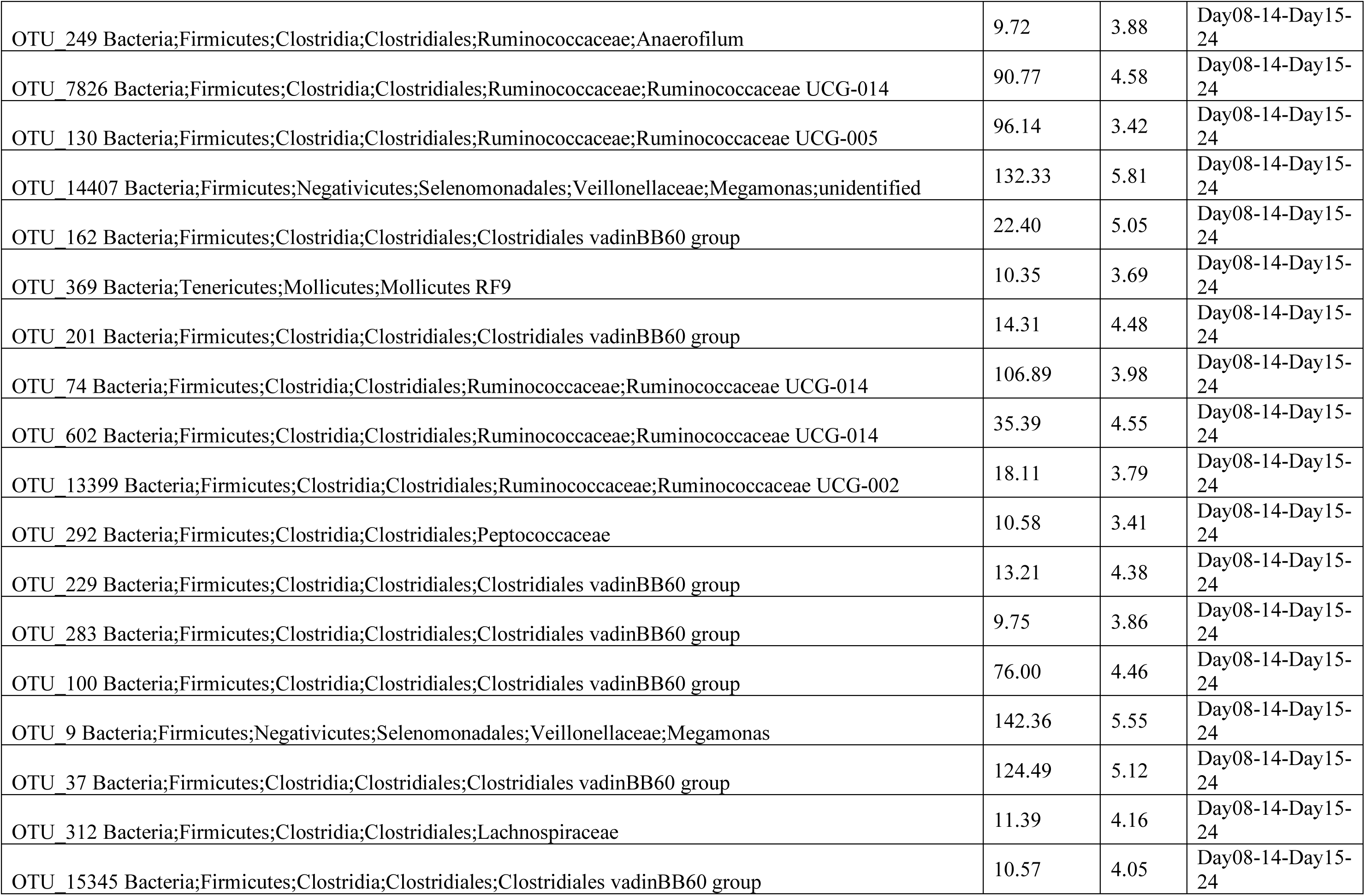
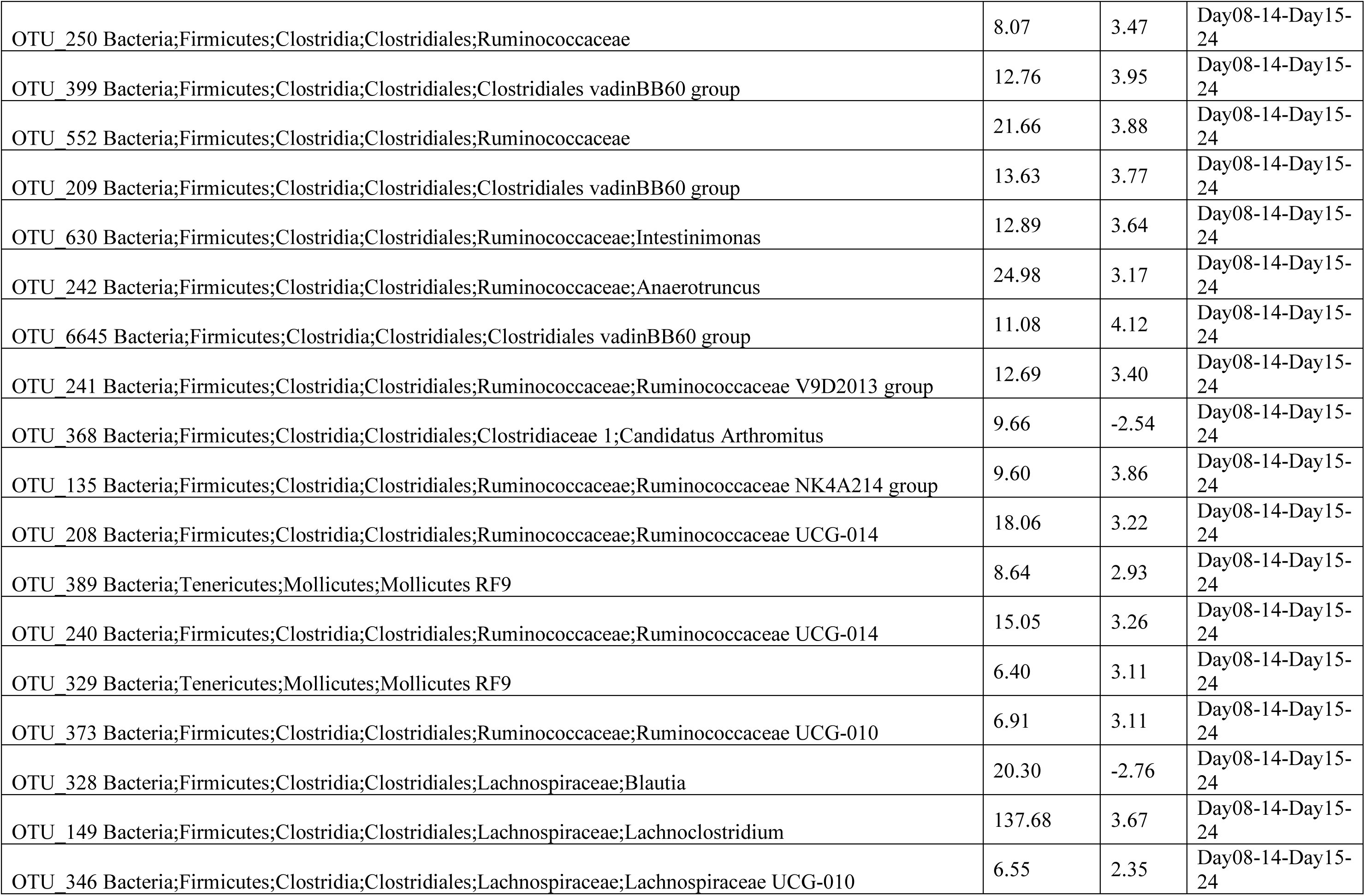
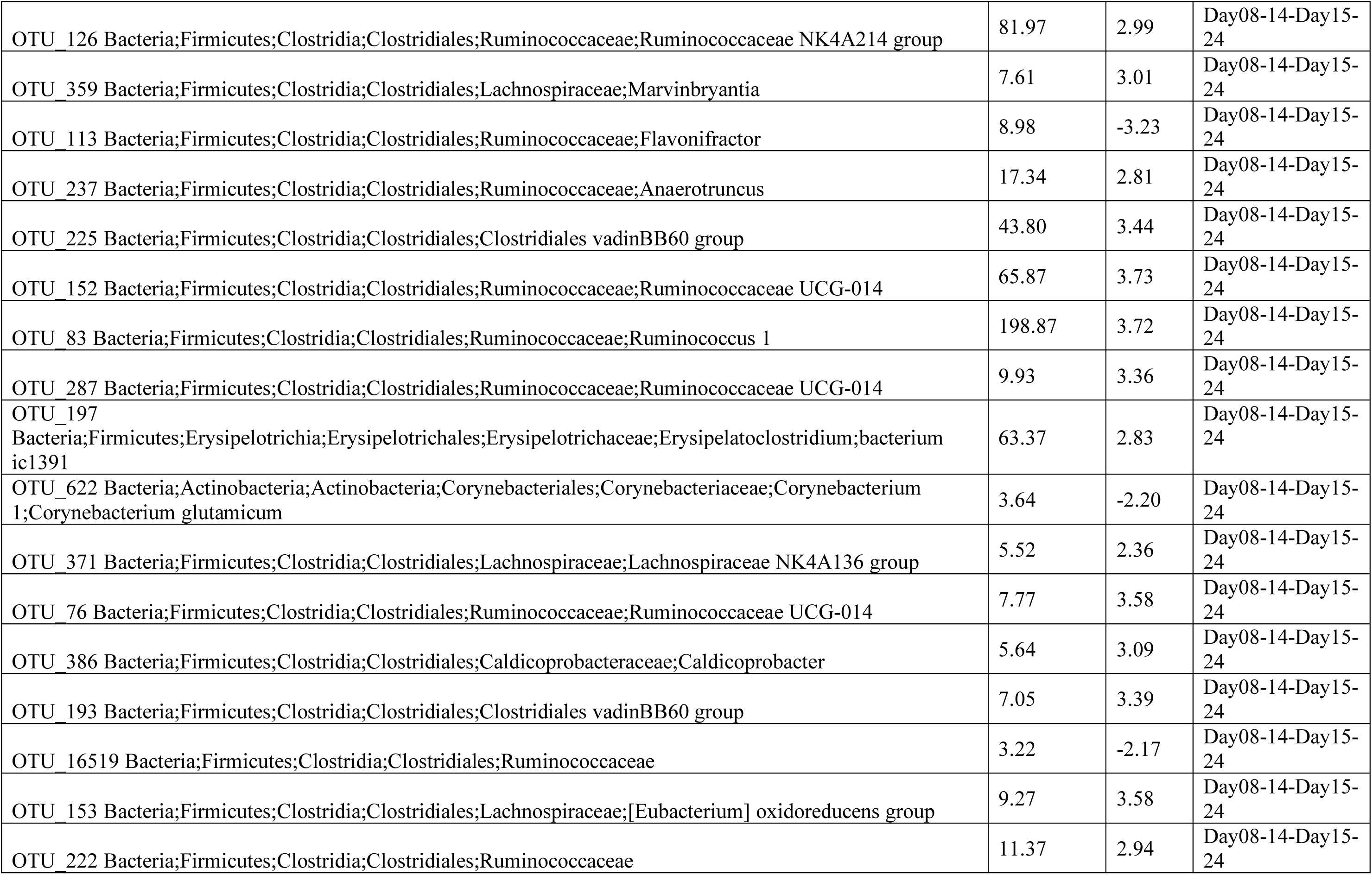
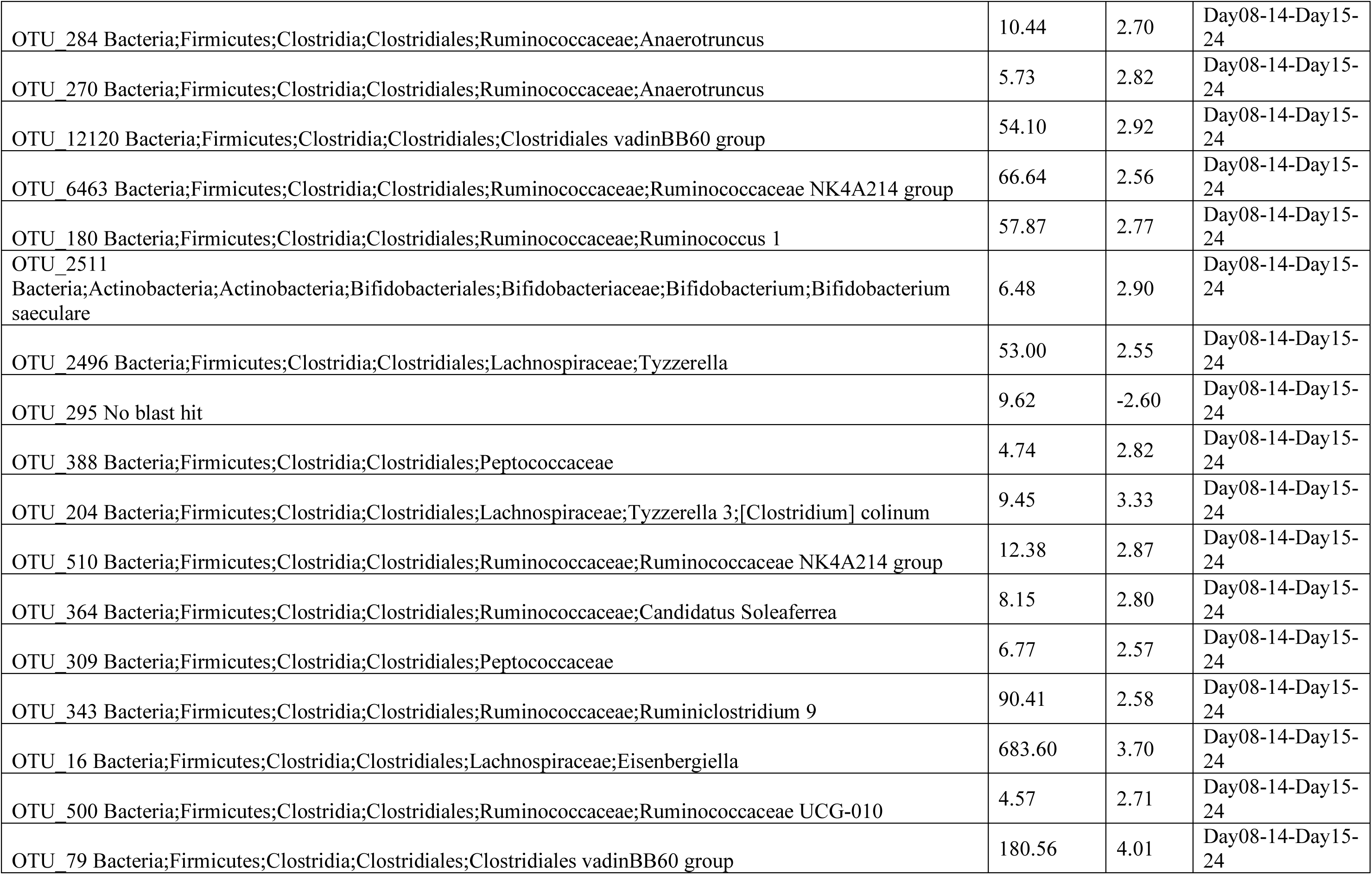
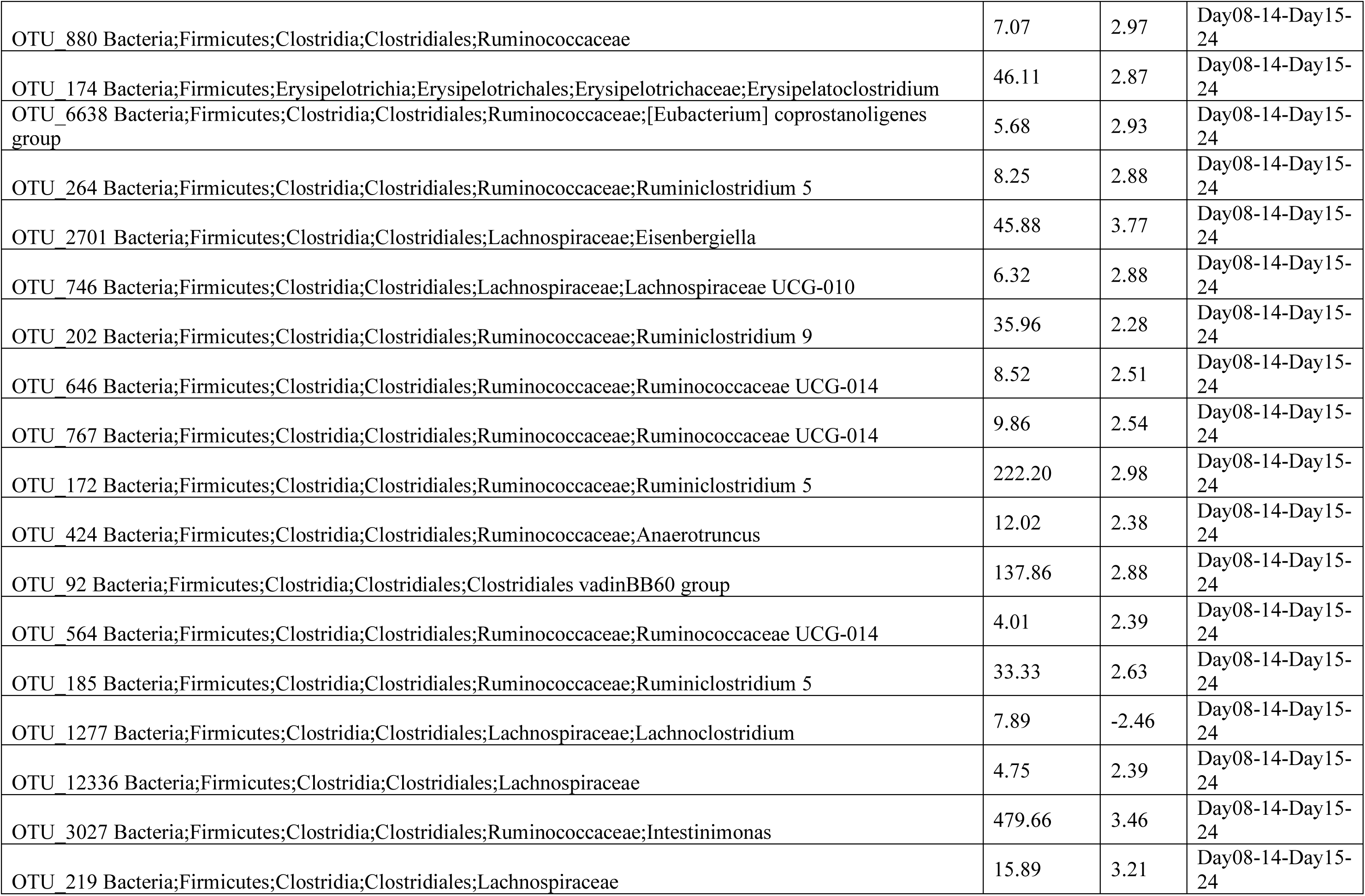
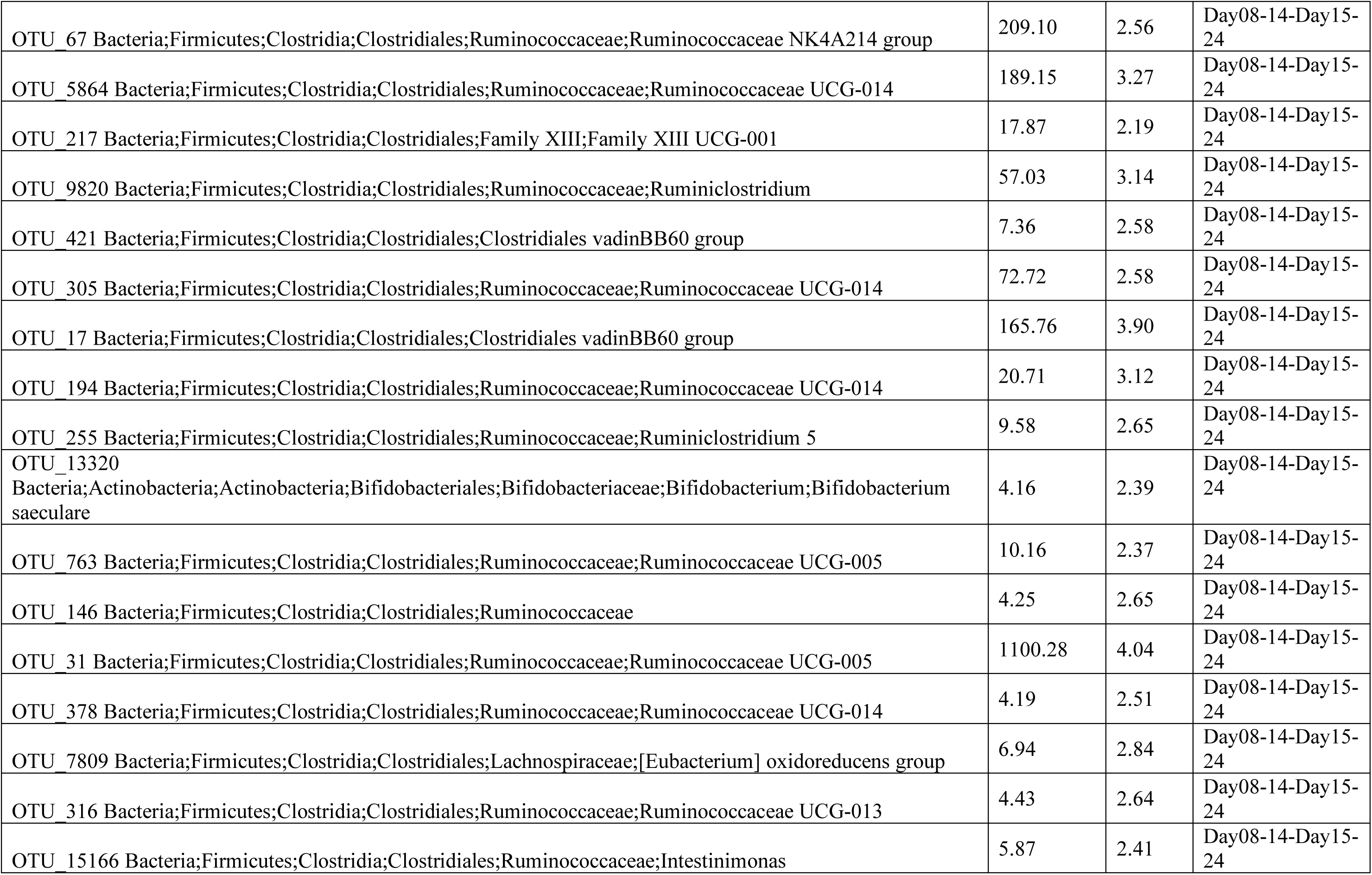
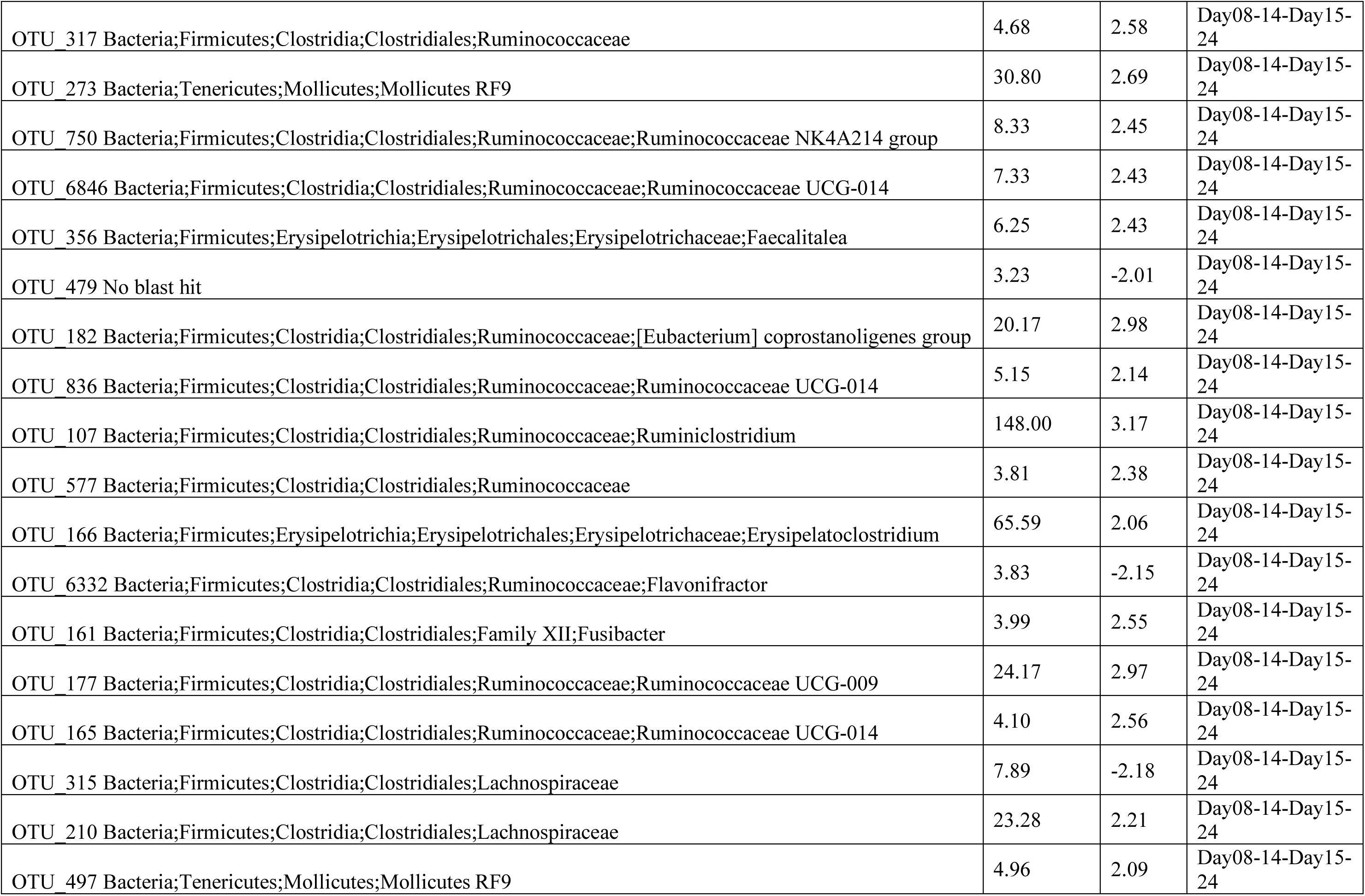
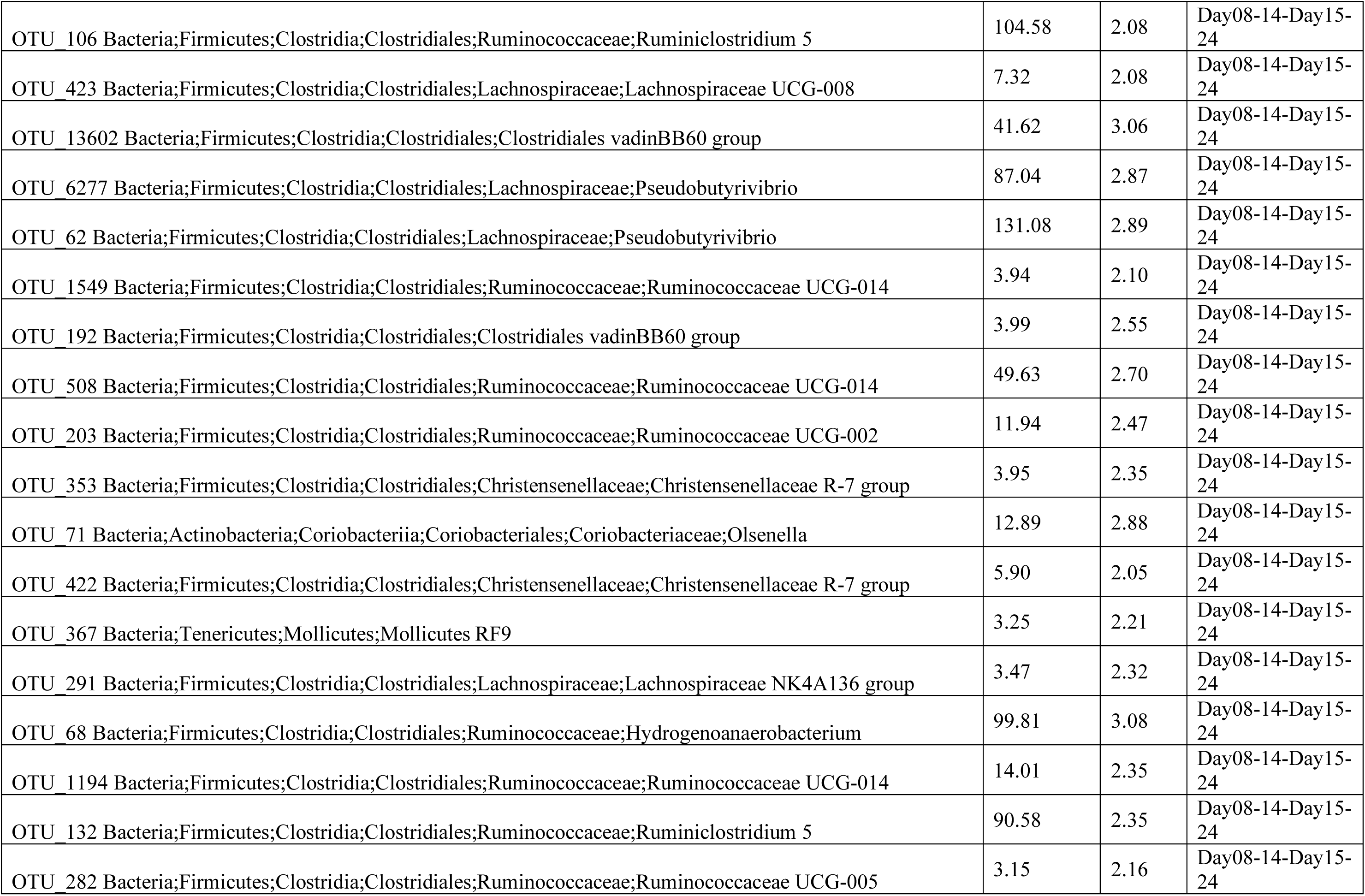
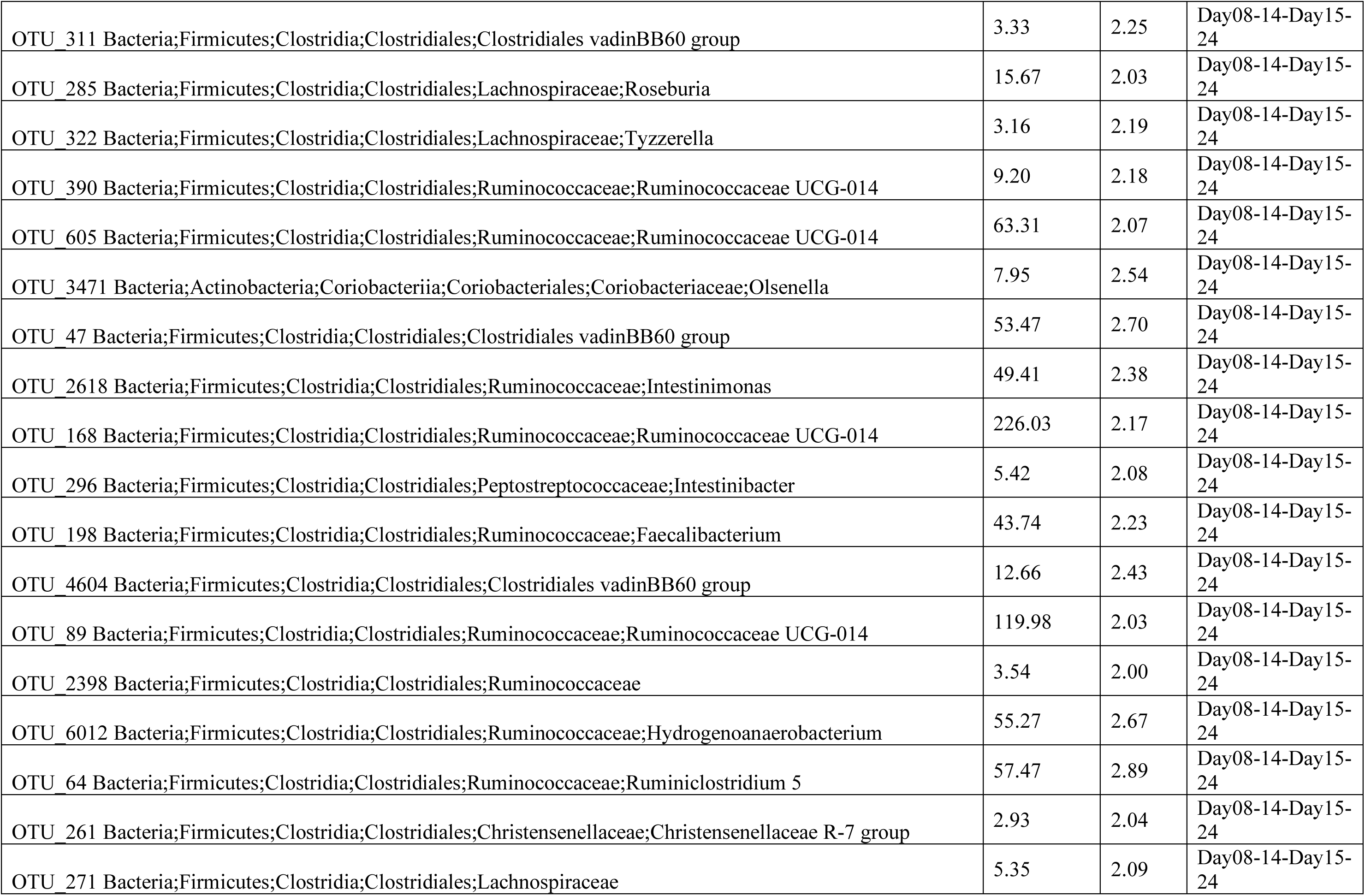
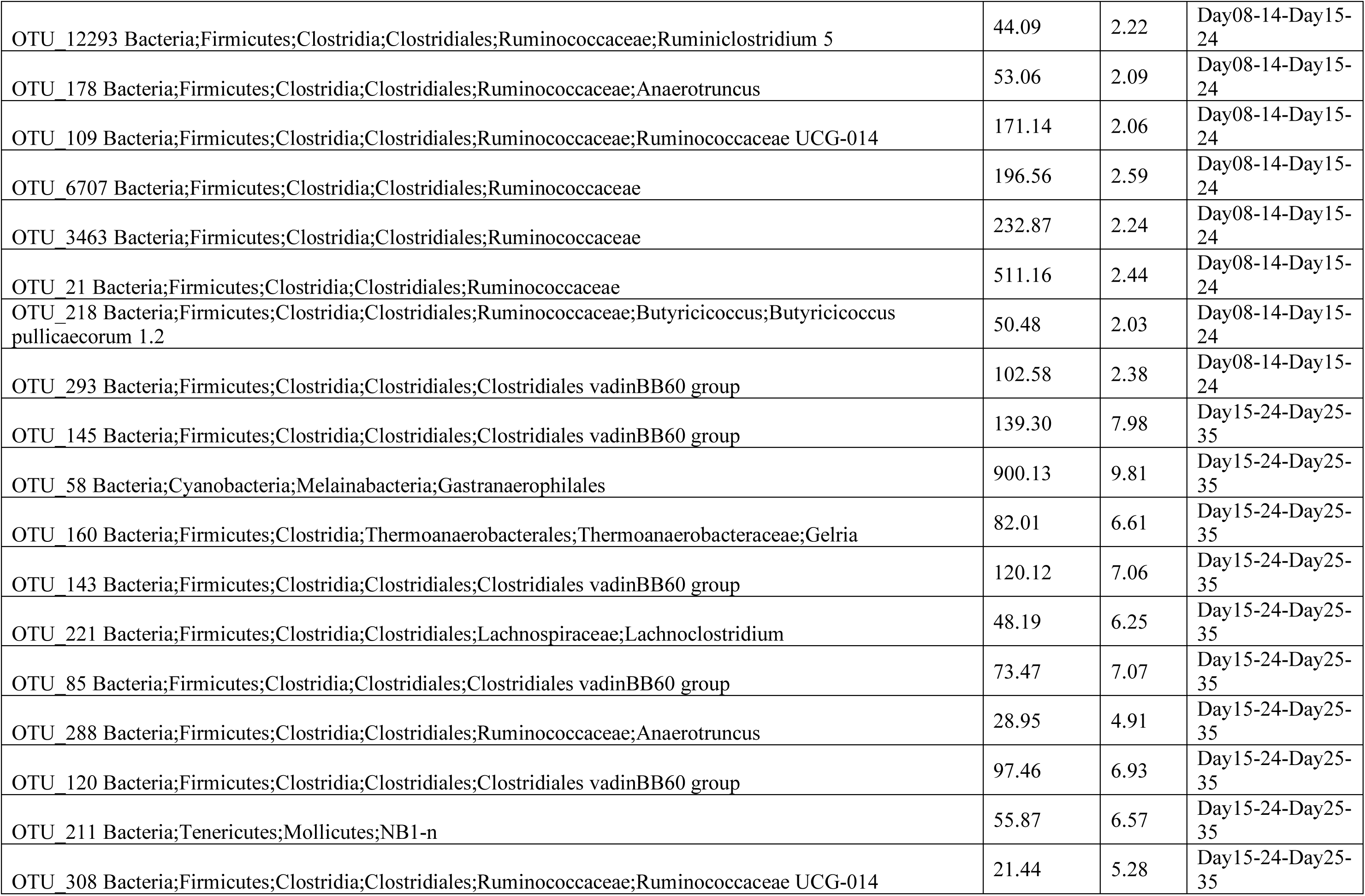
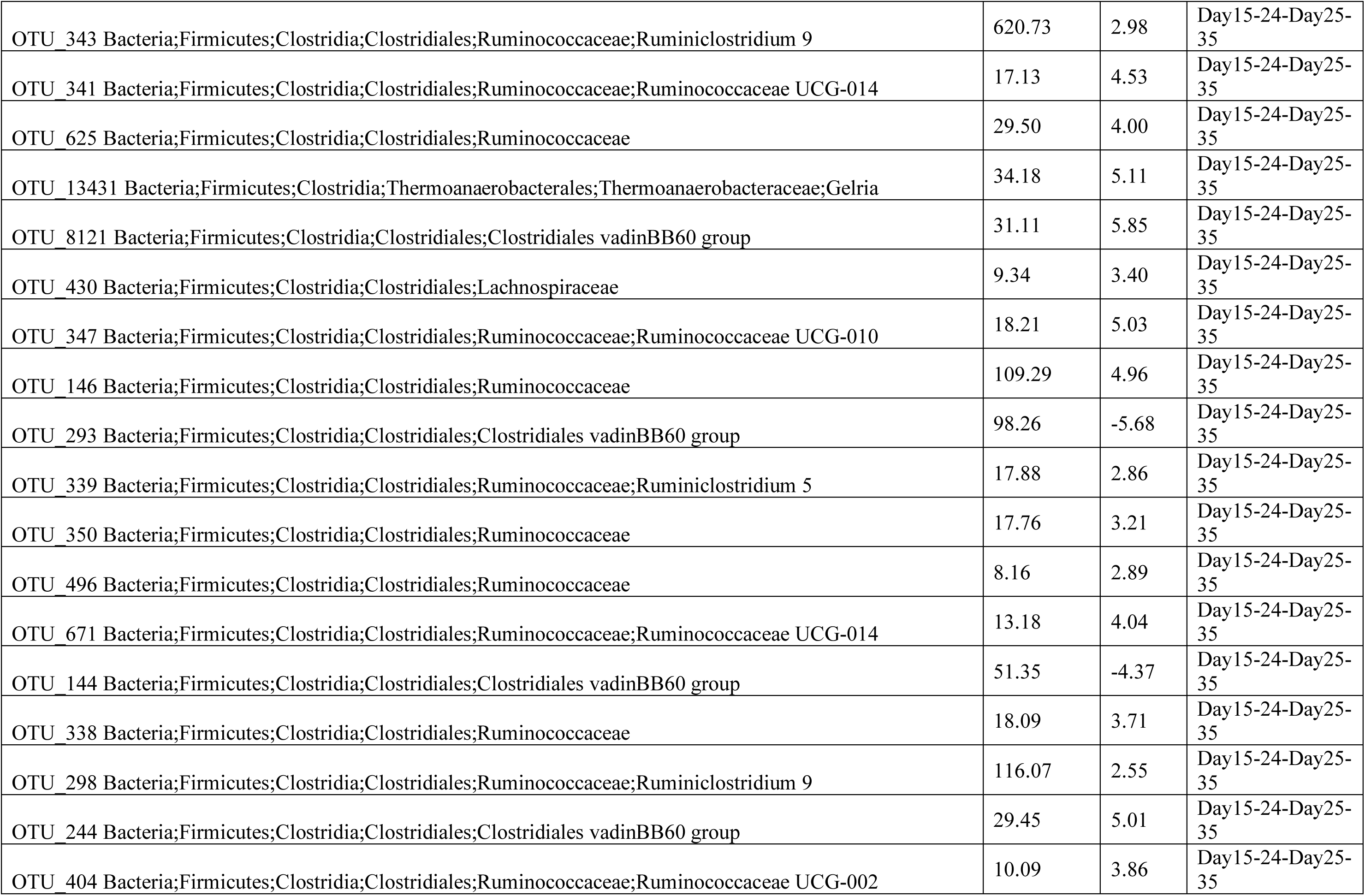
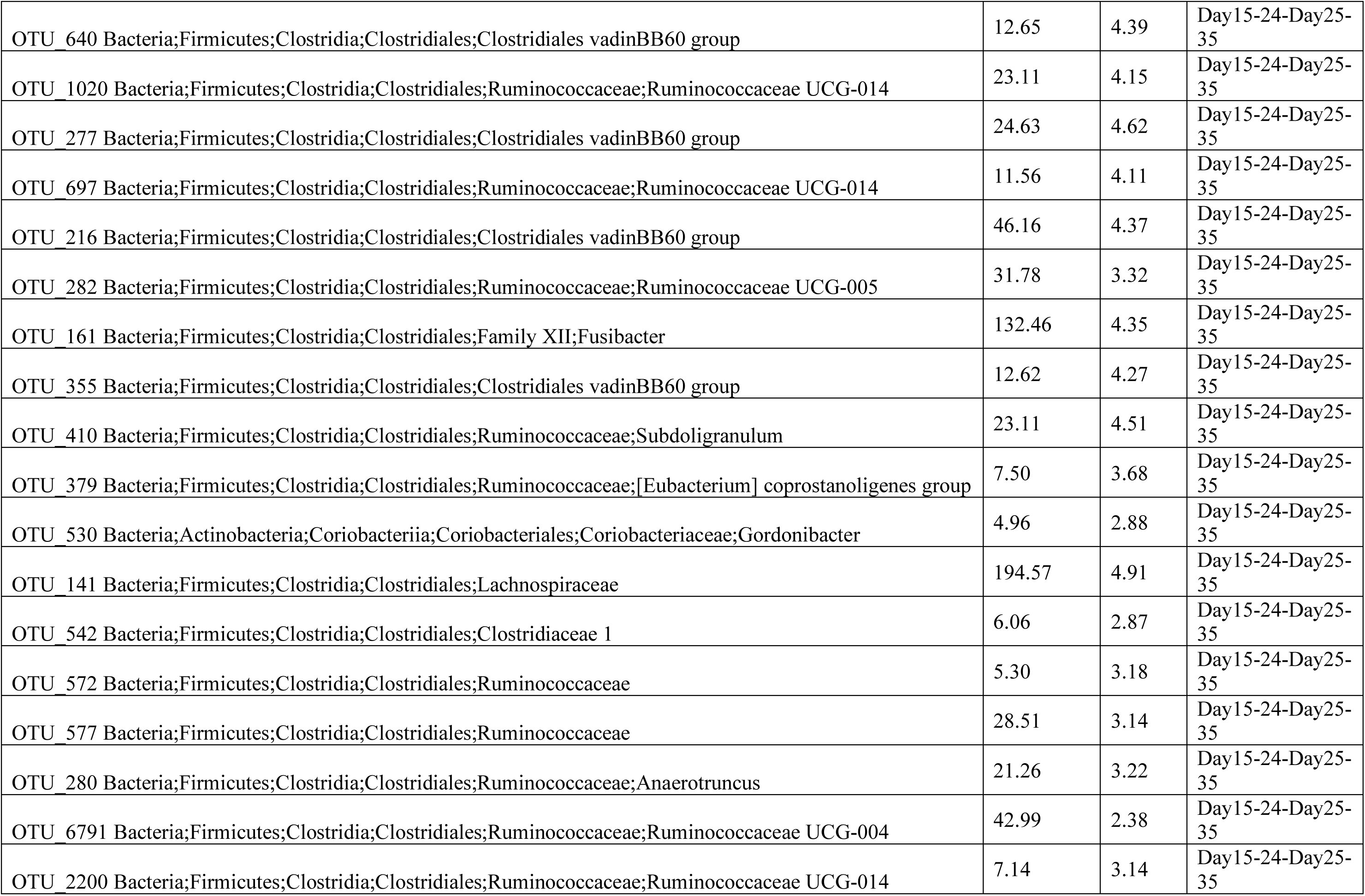
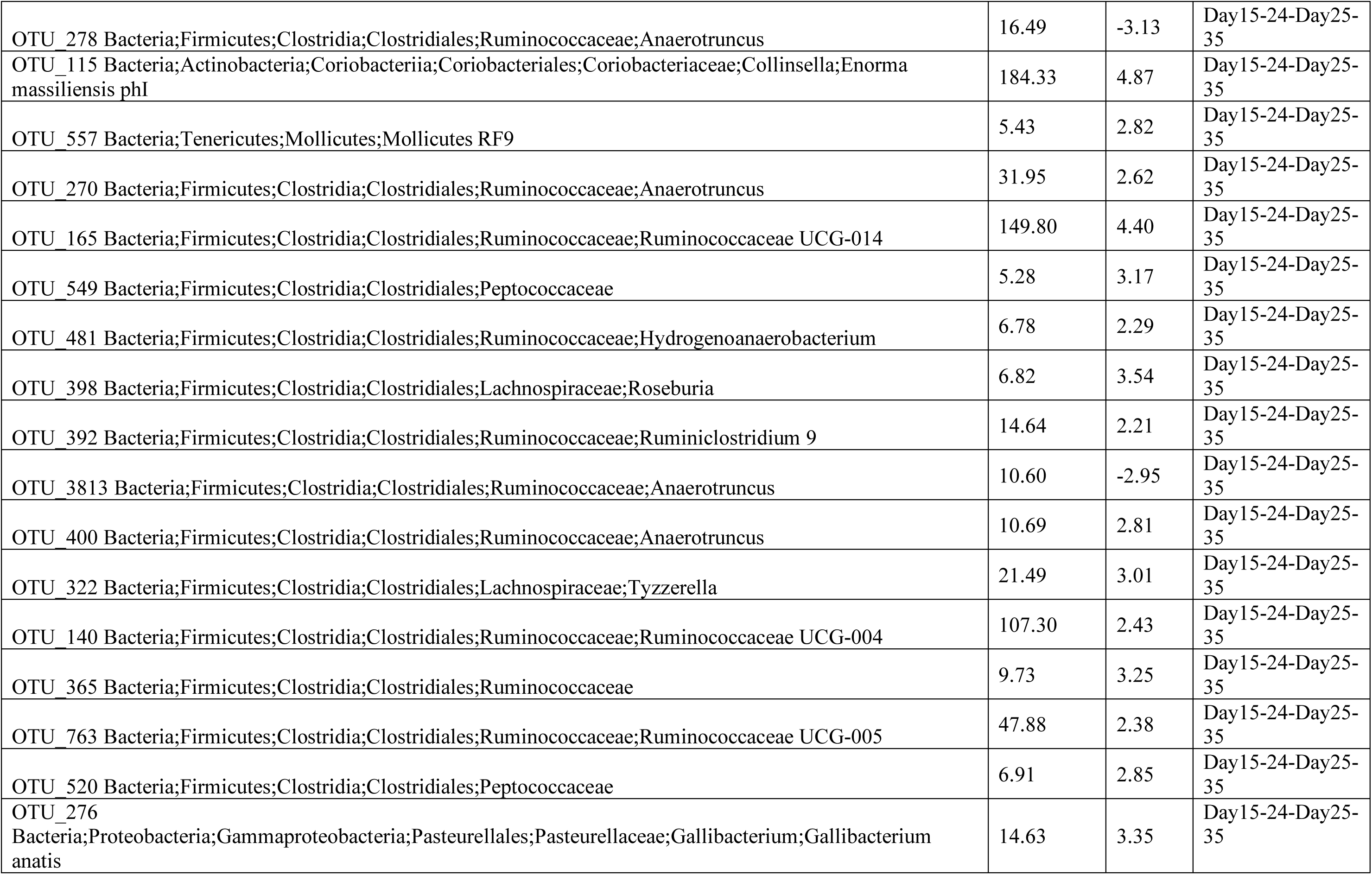
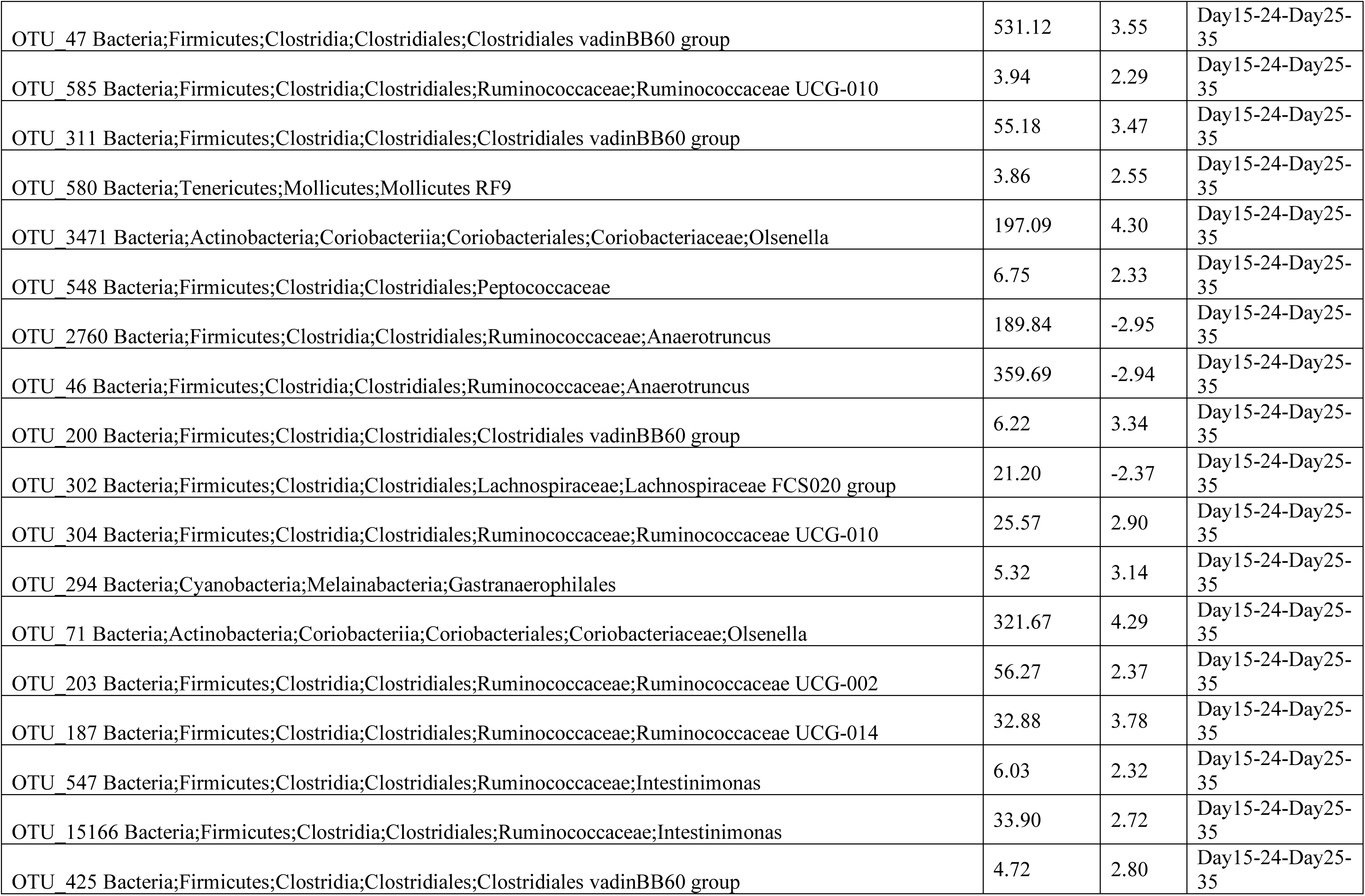
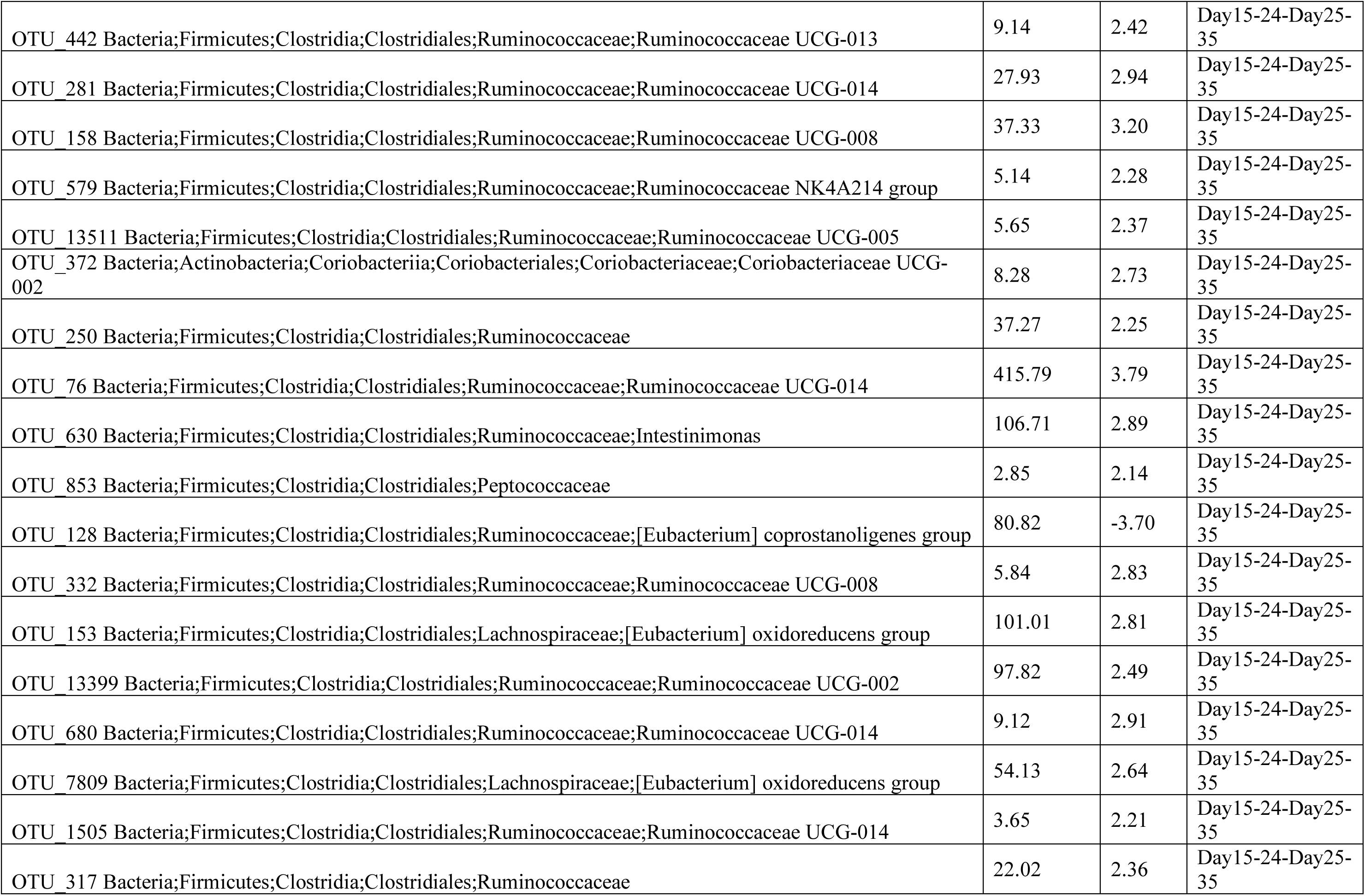
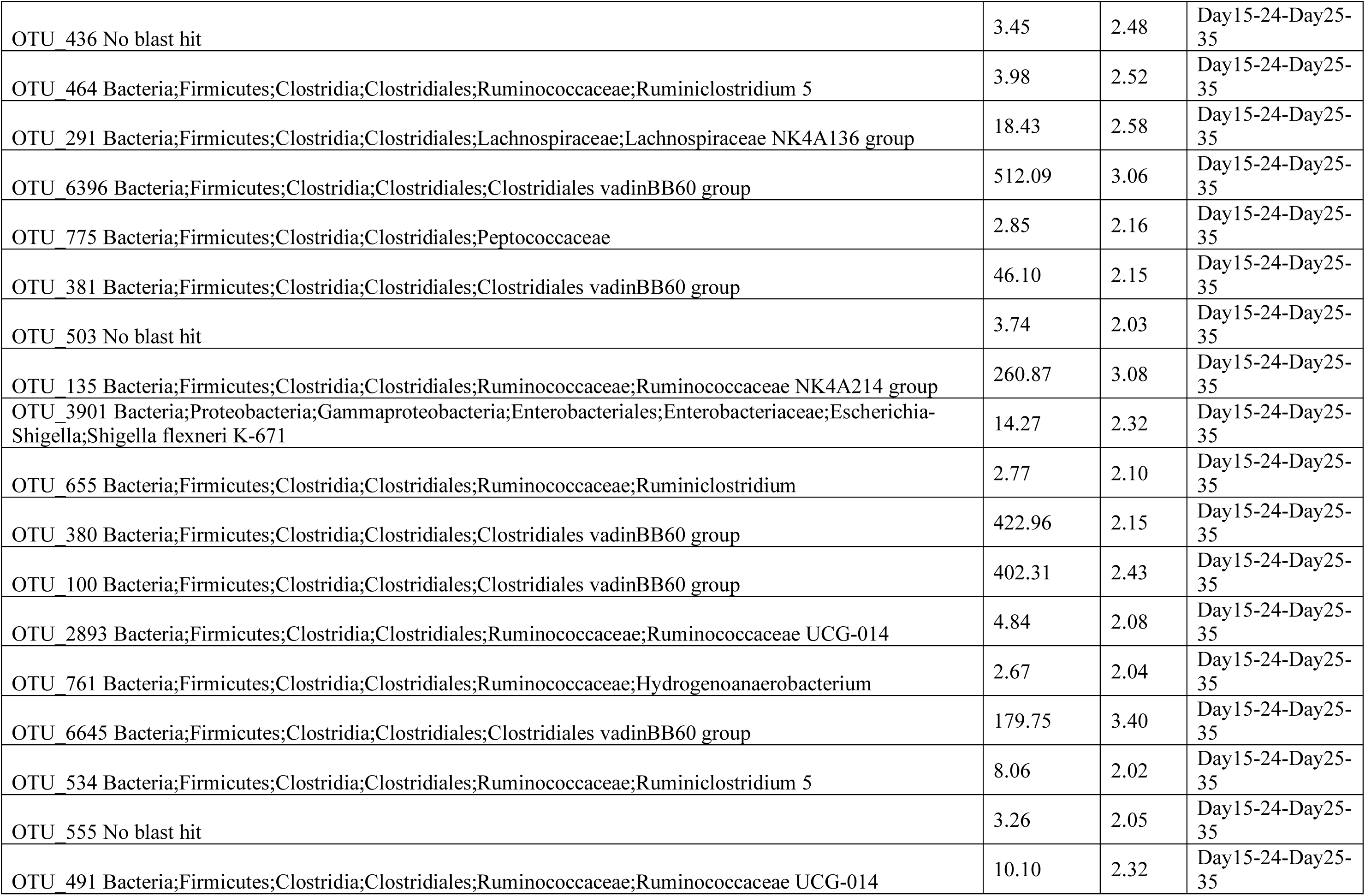
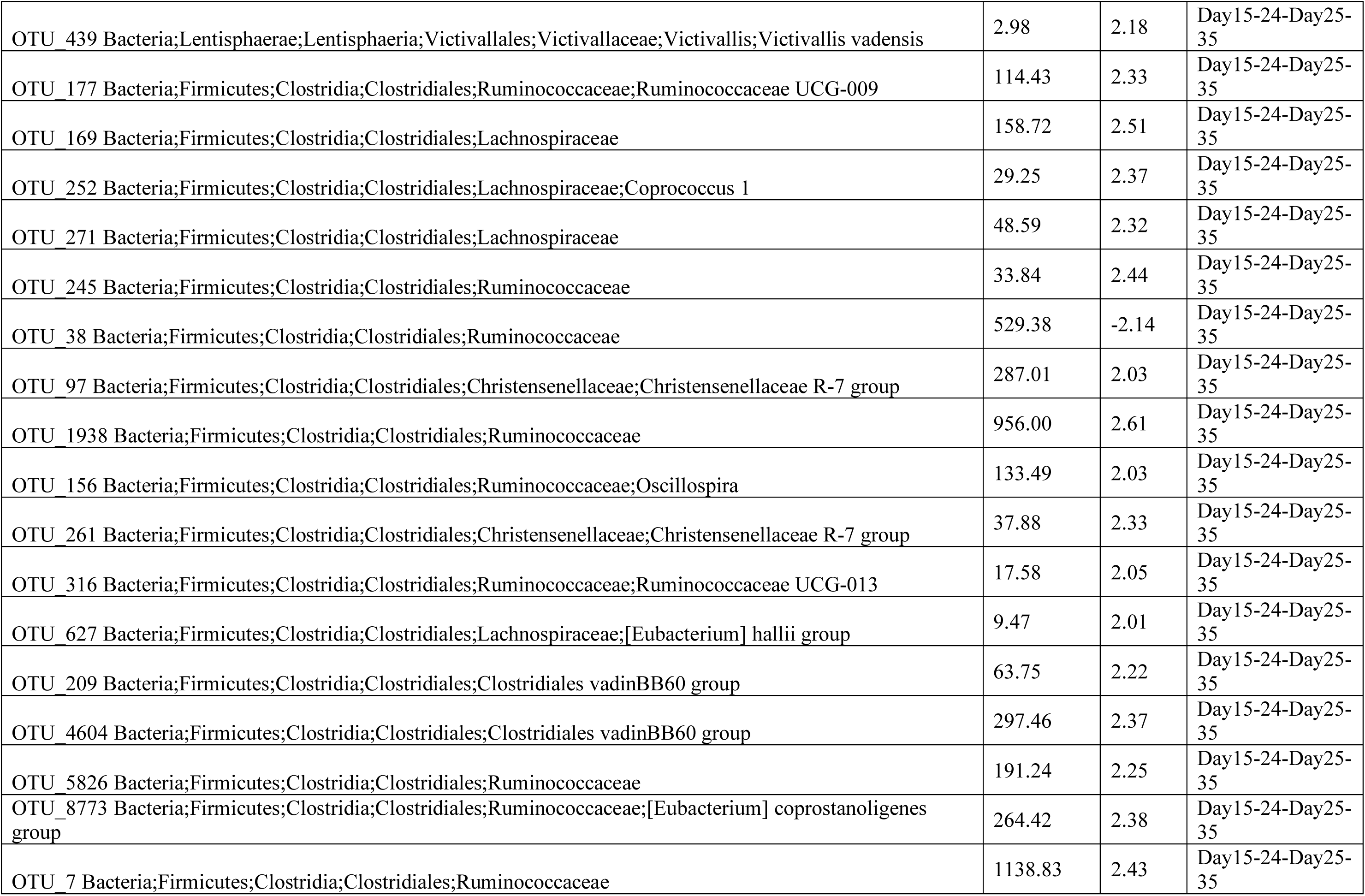
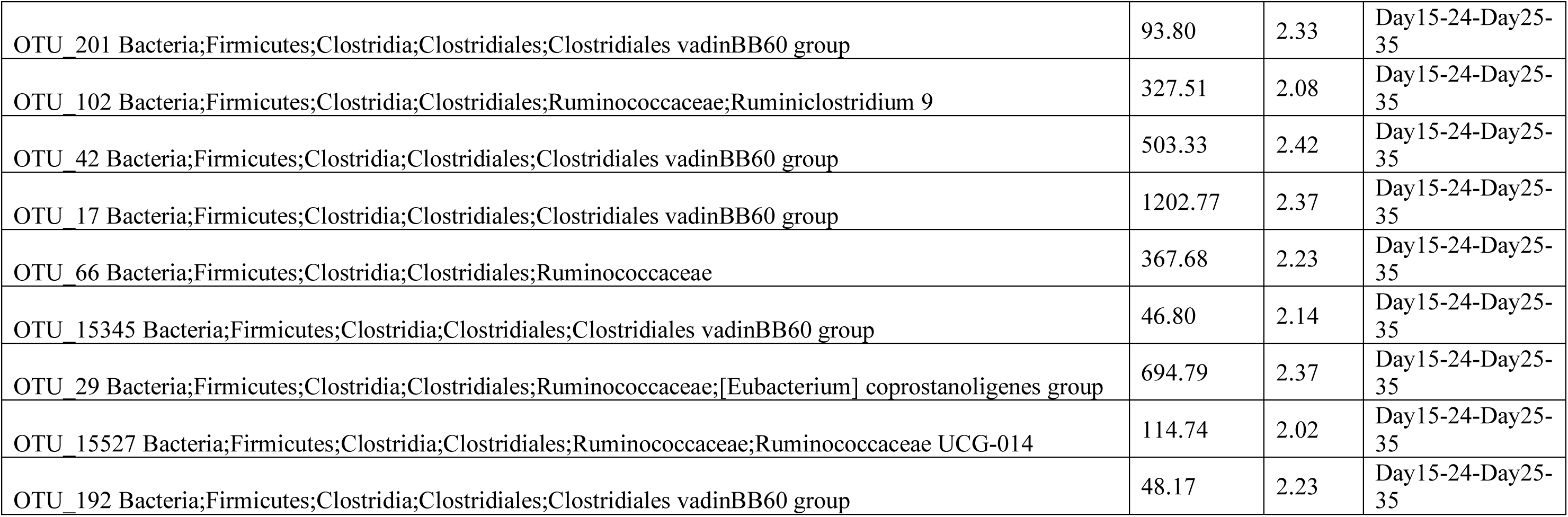
Differential analysis of OTUs that are up/down-regulated between different groups (Adjusted P values ≤ 0.05) where positive log2 fold change represent OTUs becoming abundant as we go forward in time. Here only the significant OTUs are shown for both daily and weekly comparisons.

**Supplementary Table 2:** Differential analysis of genera that are up/down regulated between different groups (Adjusted P values ≤ 0.05) where positive log2 fold change represent genera becoming abundant as we go forward in time. Here only the significant genera are shown for daily comparisons.

| Genus | Base Mean | Log2 Fold | Group |
| --- | --- | --- | --- |
| Ruminococcaceae UCG-002 | 27.48 | 4.73 | 03-04 |
| Ruminococcaceae UCG-014 | 84.78 | 4.17 | 03-04 |
| Ruminococcaceae UCG-005 | 10.12 | 3.50 | 03-04 |
| Lachnospiraceae;Lachnospira | 19.72 | 2.71 | 04-05 |
| [Eubacterium] hallii group | 83.69 | 2.48 | 04-05 |
| Enterobacter | 1975.82 | -2.55 | 04-05 |
| Candidatus Arthromitus | 5.21 | 2.54 | 05-06 |
| Prevotellaceae;Prevotella 7 | 3.27 | 2.13 | 05-06 |
| Tyzzarella | 7.12 | 2.56 | 05-06 |
| Anaerofilum | 4.03 | 2.25 | 05-06 |
| Enterococcus | 65.50 | -3.33 | 05-06 |
| Defluviitaleaceae UCG-011 | 24.57 | -3.43 | 05-06 |
| Lachnospiraceae UCG-010 | 160.54 | 2.97 | 06-07 |
| Coproccoccus 1 | 1124.60 | 2.09 | 06-07 |
| Shuttleworthia | 145.95 | 2.13 | 06-07 |
| Brevibacillus | 4.21 | -2.27 | 06-07 |
| Prevotella 7 | 3.12 | -2.06 | 06-07 |
| Tyzzarella | 73.83 | 2.90 | 07-08 |
| Faecalibacterium | 405.59 | 4.04 | 08-09 |
| Enterobacter | 35.00 | -3.30 | 08-09 |
| Family XIII UCG-001 | 4.18 | 2.19 | 09-10 |
| Roseburia | 10.30 | 2.26 | 09-10 |
| Tyzzarella | 488.55 | -2.38 | 09-10 |
| Corynebacterium 1 | 5.29 | 2.10 | 10-11 |
| Ruminococcaceae UCG-008 | 4.46 | 2.17 | 10-11 |
| Tyzzarella | 492.04 | 2.76 | 10-11 |
| Fusicatenibacter | 694.62 | -3.07 | 10-11 |
| Lachnospiraceae FE2018 group | 507.70 | -2.97 | 10-11 |
| Ruminococcus 1 | 19.47 | -2.54 | 10-11 |
| Defluviitaleaceae UCG-011 | 212.59 | -2.44 | 10-11 |
| Ruminococcaceae UCG-005 | 783.84 | -2.24 | 10-11 |
| Comamonadaceae; Delftia | 5.50 | 2.03 | 12-13 |
| Ruminococcus 1 | 50.97 | 3.13 | 12-13 |
| Butyricicoccus | 26.20 | 3.16 | 12-13 |
| Megamonas | 13.32 | -2.73 | 12-13 |
| Faecalitalea | 37.63 | 3.47 | 13-14 |
| Enterorhabdus | 6.48 | -2.03 | 13-14 |
| Ruminococcus 1 | 64.03 | 2.17 | 14-15 |
| Acetitomaculum | 38.98 | 2.17 | 14-15 |
| Family XIII UCG-001 | 20.64 | 2.39 | 14-15 |
| Bifidobacterium | 83.86 | 2.54 | 14-15 |
| Lachnospiraceae FCS020 group | 303.03 | 2.57 | 14-15 |
| Lactobacillus | 1058.97 | 2.80 | 14-15 |
| Butyricicoccus | 118.45 | 3.01 | 14-15 |
| Faecalitalea | 35.59 | -4.53 | 14-15 |
| Escherichia-Shigella | 3107.38 | -2.08 | 14-15 |
| Enterobacteriaceae;Enterobacter | 3.81 | -2.05 | 14-15 |
| Streptococcus | 8.08 | 2.91 | 15-16 |
| Ruminococcaceae UCG-009 | 334.48 | 2.44 | 16-17 |
| Ruminococcaceae V9D2013 group | 10.84 | 3.39 | 16-17 |
| Erysipelatoclostridium | 423.97 | -2.09 | 16-17 |
| Marvinbryantia | 22.84 | -2.70 | 16-17 |
| Intestinimonas | 538.49 | 2.23 | 16-17 |
| Tyzzzeria 3 | 171.81 | -2.19 | 16-17 |
| Campylobacter | 3.74 | -2.35 | 16-17 |
| Hydrogenoanaerobacterium | 21.04 | 2.82 | 17-18 |
| Ruminococcaceae UCG-008 | 10.47 | 2.52 | 18-19 |
| Ruminococcaceae UCG-004 | 40.26 | 2.15 | 18-19 |
| Oscillospira | 12.08 | 2.56 | 18-19 |
| Intestinimonas | 1028.88 | 2.10 | 19-20 |
| Escherichia-Shigella | 1946.88 | -2.68 | 19-20 |
| Proteus | 4.49 | 2.02 | 20-21 |
| Bacillus | 329.21 | 2.52 | 23-24 |
| Holdemania | 11.81 | 2.29 | 23-24 |
| Faecalibacterium | 8727.64 | 2.15 | 23-24 |
| Oscillibacter | 11.58 | 2.00 | 25-26 |
| Collinsella | 11.63 | -2.31 | 25-26 |
| Lachnospiraceae UCG-006 | 6.87 | 2.46 | 26-27 |
| Erysipelotrichaceae;Dielma | 3.46 | 2.17 | 26-27 |
| Intestinibacter | 22.43 | 2.78 | 27-28 |
| Dielma | 3.57 | -2.30 | 27-28 |
| Lachnospiraceae UCG-006 | 7.16 | -2.41 | 27-28 |
| Faecalitalea | 21.99 | -2.02 | 27-28 |
| Coriobacteriaceae UCG-002 | 25.30 | 3.37 | 28-29 |
| Bacillus | 562.14 | 3.18 | 30-31 |

**Supplementary Table 3:** Differential analysis of pathways becoming significant based on Kruskal-Wallis test (Adjusted P values ≤ 0.05). Here results are shown for both daily and weekly comparisons.

| KEGG KOs | P Values | Adjusted P Values | Upregulated | Group Comparison |
| --- | --- | --- | --- | --- |
| ko00061; Fatty acid biosynthesis | 0.007911789 | 0.037310224 | 03 | 03 - 04 |
| ko00240; Pyrimidine metabolism | 0.009374768 | 0.037310224 | 04 | 03 - 04 |
| ko00250; Alanine, aspartate and glutamate metabolism | 0.006656727 | 0.037310224 | 04 | 03 - 04 |
| ko00290; Valine, leucine and isoleucine biosynthesis | 0.009374768 | 0.037310224 | 04 | 03 - 04 |
| ko00330; Arginine and proline metabolism | 0.009374768 | 0.037310224 | 04 | 03 - 04 |
| ko00350; Tyrosine metabolism | 0.011074438 | 0.037310224 | 04 | 03 - 04 |
| ko00460; Cyanoamino acid metabolism | 0.002213692 | 0.037310224 | 03 | 03 - 04 |
| ko00524; Butirosin and neomycin biosynthesis | 0.009374768 | 0.037310224 | 03 | 03 - 04 |
| ko00562; Inositol phosphate metabolism | 0.006656727 | 0.037310224 | 03 | 03 - 04 |
| ko00624; Polycyclic aromatic hydrocarbon degradation | 0.01304252 | 0.037310224 | 04 | 03 - 04 |
| ko00730; Thiamine metabolism | 0.011074438 | 0.037310224 | 04 | 03 - 04 |
| ko00740; Riboflavin metabolism | 0.01304252 | 0.037310224 | 03 | 03 - 04 |
| ko00750; Vitamin B6 metabolism | 0.007911789 | 0.037310224 | 04 | 03 - 04 |
| ko00770; Pantothenate and CoA biosynthesis | 0.009374768 | 0.037310224 | 04 | 03 - 04 |
| ko00780; Biotin metabolism | 0.01304252 | 0.037310224 | 04 | 03 - 04 |
| ko00906; Carotenoid biosynthesis | 0.01304252 | 0.037310224 | 04 | 03 - 04 |
| ko00960; Tropane, piperidine and pyridine alkaloid biosynthesis | 0.007911789 | 0.037310224 | 03 | 03 - 04 |
| ko01051; Biosynthesis of ansamycins | 0.009374768 | 0.037310224 | 03 | 03 - 04 |
| ko02010; ABC transporters | 0.009374768 | 0.037310224 | 03 | 03 - 04 |
| ko03008; Ribosome biogenesis in eukaryotes | 0.011074438 | 0.037310224 | 03 | 03 - 04 |
| ko03018; RNA degradation | 0.006656727 | 0.037310224 | 03 | 03 - 04 |
| ko03410; Base excision repair | 0.011074438 | 0.037310224 | 03 | 03 - 04 |
| ko04020; Calcium signaling pathway | 0.009374768 | 0.037310224 | 04 | 03 - 04 |
| ko04122; Sulfur relay system | 0.009374768 | 0.037310224 | 04 | 03 - 04 |
| ko04141; Protein processing in endoplasmic reticulum | 0.007911789 | 0.037310224 | 04 | 03 - 04 |
| ko04146; Peroxisome | 0.007911789 | 0.037310224 | 03 | 03 - 04 |
| ko04260; Cardiac muscle contraction | 0.011074438 | 0.037310224 | 03 | 03 - 04 |
| ko04370; VEGF signaling pathway | 0.011074438 | 0.037310224 | 04 | 03 - 04 |
| ko04626; Plant-pathogen interaction | 0.007911789 | 0.037310224 | 03 | 03 - 04 |
| ko04666; Fc gamma R-mediated phagocytosis | 0.009374768 | 0.037310224 | 04 | 03 - 04 |
| ko04728; Dopaminergic synapse | 0.01304252 | 0.037310224 | 03 | 03 - 04 |
| ko04930; Type II diabetes mellitus | 0.011074438 | 0.037310224 | 03 | 03 - 04 |
| ko04974; Protein digestion and absorption | 0.007911789 | 0.037310224 | 03 | 03 - 04 |
| ko05030; Cocaine addiction | 0.005583617 | 0.037310224 | 03 | 03 - 04 |
| ko05031; Amphetamine addiction | 0.005583617 | 0.037310224 | 03 | 03 - 04 |
| ko05034; Alcoholism | 0.01304252 | 0.037310224 | 03 | 03 - 04 |
| ko00190; Oxidative phosphorylation | 0.015313822 | 0.03909974 | 04 | 03 - 04 |
| ko00561; Glycerolipid metabolism | 0.015313822 | 0.03909974 | 03 | 03 - 04 |
| ko04070; Phosphatidylinositol signaling system | 0.015313822 | 0.03909974 | 03 | 03 - 04 |
| ko04112; Cell cycle - Caulobacter | 0.015313822 | 0.03909974 | 04 | 03 - 04 |
| ko04726; Serotonergic synapse | 0.015313822 | 0.03909974 | 03 | 03 - 04 |
| ko05032; Morphine addiction | 0.015313822 | 0.03909974 | 03 | 03 - 04 |
| ko05203; Viral carcinogenesis | 0.015313822 | 0.03909974 | 03 | 03 - 04 |
| ko04144; Endocytosis | 0.000219851 | 0.007285107 | 05 | 05-06 |
| ko04912; GnRH signaling pathway | 0.000219851 | 0.007285107 | 05 | 05-06 |
| ko05012; Parkinsons disease | 0.000219851 | 0.007285107 | 05 | 05-06 |
| ko00030; Pentose phosphate pathway | 0.001822735 | 0.013247142 | 05 | 05-06 |
| ko00053; Ascorbate and aldarate metabolism | 0.002213692 | 0.013247142 | 05 | 05-06 |
| ko00100; Steroid biosynthesis | 0.001822735 | 0.013247142 | 05 | 05-06 |
| ko00196; Photosynthesis - antenna proteins | 0.002208779 | 0.013247142 | 05 | 05-06 |
| ko00540; Lipopolysaccharide biosynthesis | 0.000998686 | 0.013247142 | 05 | 05-06 |
| ko00592; alpha-Linolenic acid metabolism | 0.000275504 | 0.013247142 | 05 | 05-06 |
| ko00623; Toluene degradation | 0.000275504 | 0.013247142 | 05 | 05-06 |
| ko00965; Betalain biosynthesis | 0.000532006 | 0.013247142 | 05 | 05-06 |
| ko00984; Steroid degradation | 0.001496164 | 0.013247142 | 05 | 05-06 |
| ko01057; Biosynthesis of type II polyketide products | 0.000998686 | 0.013247142 | 05 | 05-06 |
| ko02060; Phosphotransferase system (PTS) | 0.001224283 | 0.013247142 | 05 | 05-06 |
| ko04610; Complement and coagulation cascades | 0.001221273 | 0.013247142 | 05 | 05-06 |
| ko05014; Amyotrophic lateral sclerosis (ALS) | 0.000998686 | 0.013247142 | 05 | 05-06 |
| ko05020; Prion diseases | 0.000658337 | 0.013247142 | 05 | 05-06 |
| ko05131; Shigellosis | 0.001221273 | 0.013247142 | 05 | 05-06 |
| ko05142; Chagas disease (American trypanosomiasis) | 0.001822735 | 0.013247142 | 05 | 05-06 |
| ko05204; Chemical carcinogenesis | 0.001224283 | 0.013247142 | 05 | 05-06 |
| ko00363; Bisphenol degradation | 0.002680172 | 0.036672283 | 05 | 05-06 |
| ko00630; Glyoxylate and dicarboxylate metabolism | 0.002680172 | 0.036672283 | 05 | 05-06 |
| ko00901; Indole alkaloid biosynthesis | 0.002680172 | 0.036672283 | 05 | 05-06 |
| ko00920; Sulfur metabolism | 0.002680172 | 0.036672283 | 05 | 05-06 |
| ko00945; Stilbenoid, diarylheptanoid and gingerol biosynthesis | 0.002680172 | 0.036672283 | 05 | 05-06 |
| ko00980; Metabolism of xenobiotics by cytochrome P450 | 0.002680172 | 0.036672283 | 05 | 05-06 |
| ko00982; Drug metabolism - cytochrome P450 | 0.002680172 | 0.036672283 | 05 | 05-06 |
| ko05111; Vibrio cholerae pathogenic cycle | 0.002680172 | 0.036672283 | 05 | 05-06 |
| ko05132; Salmonella infection | 0.002680172 | 0.036672283 | 05 | 05-06 |
| ko04115; p53 signaling pathway | 0.00128048 | 0.028496707 | 10 | 10 -11 |
| ko00909; Sesquiterpenoid and triterpenoid biosynthesis | 0.001702519 | 0.045759699 | 10 | 10 -11 |
| ko05168; Herpes simplex infection | 0.001702519 | 0.045759699 | 10 | 10 -11 |
| ko05210; Colorectal cancer | 0.001702519 | 0.045759699 | 10 | 10 -11 |
| ko05416; Viral myocarditis | 0.001702519 | 0.045759699 | 10 | 10 -11 |
| ko02040; Flagellar assembly | 0.000165522 | 0.012327241 | 14 | 14 - 15 |
| ko05111; Vibrio cholerae pathogenic cycle | 0.000219473 | 0.012327241 | 14 | 14 - 15 |
| ko05130; Pathogenic Escherichia coli infection | 0.000380365 | 0.012327241 | 14 | 14 - 15 |
| ko00364; Fluorobenzoate degradation | 0.001078987 | 0.0146559 | 14 | 14 - 15 |
| ko00592; alpha-Linolenic acid metabolism | 0.001078987 | 0.0146559 | 14 | 14 - 15 |
| ko00633; Nitrotoluene degradation | 0.000646749 | 0.0146559 | 14 | 14 - 15 |
| ko00710; Carbon fixation in photosynthetic organisms | 0.000497167 | 0.0146559 | 15 | 14 - 15 |
| ko03070; Bacterial secretion system | 0.000497167 | 0.0146559 | 14 | 14 - 15 |
| ko05132; Salmonella infection | 0.000497167 | 0.0146559 | 14 | 14 - 15 |
| ko00240; Pyrimidine metabolism | 0.001766337 | 0.01897042 | 15 | 14 - 15 |
| ko00760; Nicotinate and nicotinamide metabolism | 0.001383791 | 0.01897042 | 15 | 14 - 15 |
| ko00920; Sulfur metabolism | 0.001383791 | 0.01897042 | 14 | 14 - 15 |
| ko00960; Tropane, piperidine and pyridine alkaloid biosynthesis | 0.001383791 | 0.01897042 | 14 | 14 - 15 |
| ko02020; Two-component system | 0.001383791 | 0.01897042 | 14 | 14 - 15 |
| ko03440; Homologous recombination | 0.001766337 | 0.01897042 | 15 | 14 - 15 |
| ko05032; Morphine addiction | 0.001383791 | 0.01897042 | 14 | 14 - 15 |
| ko05133; Pertussis | 0.001383791 | 0.01897042 | 14 | 14 - 15 |
| ko00020; Citrate cycle (TCA cycle) | 0.002837545 | 0.022375931 | 14 | 14 - 15 |
| ko00232; Caffeine metabolism | 0.003571236 | 0.022375931 | 14 | 14 - 15 |
| ko00281; Geraniol degradation | 0.003571236 | 0.022375931 | 14 | 14 - 15 |
| ko00310; Lysine degradation | 0.001766337 | 0.022375931 | 14 | 14 - 15 |
| ko00473; D-Alanine metabolism | 0.002837545 | 0.022375931 | 15 | 14 - 15 |
| ko00511; Other glycan degradation | 0.003571236 | 0.022375931 | 15 | 14 - 15 |
| ko00603; Glycosphingolipid biosynthesis - globo series | 0.003571236 | 0.022375931 | 15 | 14 - 15 |
| ko00624; Polycyclic aromatic hydrocarbon degradation | 0.002837545 | 0.022375931 | 15 | 14 - 15 |
| ko00630; Glyoxylate and dicarboxylate metabolism | 0.002837545 | 0.022375931 | 14 | 14 - 15 |
| ko00640; Propanoate metabolism | 0.002244033 | 0.022375931 | 14 | 14 - 15 |
| ko00740; Riboflavin metabolism | 0.003571236 | 0.022375931 | 14 | 14 - 15 |
| ko03030; DNA replication | 0.003571236 | 0.022375931 | 15 | 14 - 15 |
| ko04113; Meiosis - yeast | 0.002244033 | 0.022375931 | 14 | 14 - 15 |
| ko00791; Atrazine degradation | 0.004473649 | 0.041073328 | 14 | 14 - 15 |
| ko00906; Carotenoid biosynthesis | 0.004473649 | 0.041073328 | 15 | 14 - 15 |
| ko00910; Nitrogen metabolism | 0.004473649 | 0.041073328 | 14 | 14 - 15 |
| ko01040; Biosynthesis of unsaturated fatty acids | 0.004473649 | 0.041073328 | 14 | 14 - 15 |
| ko03010; Ribosome | 0.004473649 | 0.041073328 | 15 | 14 - 15 |
| ko05150; Staphylococcus aureus infection | 0.004473649 | 0.041073328 | 15 | 14 - 15 |
| ko00331; Clavulanic acid biosynthesis | 1.16E-09 | 4.38E-09 | Day03-07 | Day03-07 - Day08-14 |
| ko00592; alpha-Linolenic acid metabolism | 1.05E-09 | 4.38E-09 | Day03-07 | Day03-07 - Day08-14 |
| ko00920; Sulfur metabolism | 1.07E-09 | 4.38E-09 | Day03-07 | Day03-07 - Day08-14 |
| ko04011; MAPK signaling pathway - yeast | 1.13E-09 | 4.38E-09 | Day08-14 | Day03-07 - Day08-14 |
| ko04723; Retrograde endocannabinoid signaling | 8.68E-10 | 4.38E-09 | Day03-07 | Day03-07 - Day08-14 |
| ko04726; Serotonergic synapse | 8.91E-10 | 4.38E-09 | Day03-07 | Day03-07 - Day08-14 |
| ko05020; Prion diseases | 4.30E-10 | 4.38E-09 | Day03-07 | Day03-07 - Day08-14 |
| ko05132; Salmonella infection | 2.56E-10 | 4.38E-09 | Day03-07 | Day03-07 - Day08-14 |
| ko05133; Pertussis | 8.01E-10 | 4.38E-09 | Day03-07 | Day03-07 - Day08-14 |
| ko05014; Amyotrophic lateral sclerosis (ALS) | 1.19E-09 | 6.19E-09 | Day03-07 | Day03-07 - Day08-14 |
| ko00380; Tryptophan metabolism | 6.45E-09 | 1.99E-08 | Day03-07 | Day03-07 - Day08-14 |
| ko01053; Biosynthesis of siderophore group nonribosomal peptides | 5.26E-09 | 1.99E-08 | Day03-07 | Day03-07 - Day08-14 |
| ko04144; Endocytosis | 5.98E-09 | 1.99E-08 | Day03-07 | Day03-07 - Day08-14 |
| ko04610; Complement and coagulation cascades | 5.13E-09 | 1.99E-08 | Day03-07 | Day03-07 - Day08-14 |
| ko04912; GnRH signaling pathway | 5.98E-09 | 1.99E-08 | Day03-07 | Day03-07 - Day08-14 |
| ko05034; Alcoholism | 3.15E-09 | 1.99E-08 | Day03-07 | Day03-07 - Day08-14 |
| ko05110; Vibrio cholerae infection | 1.36E-09 | 1.99E-08 | Day03-07 | Day03-07 - Day08-14 |
| ko05111; Vibrio cholerae pathogenic cycle | 4.63E-09 | 1.99E-08 | Day03-07 | Day03-07 - Day08-14 |
| ko05131; Shigellosis | 4.40E-09 | 1.99E-08 | Day03-07 | Day03-07 - Day08-14 |
| ko05219; Bladder cancer | 3.32E-09 | 1.99E-08 | Day03-07 | Day03-07 - Day08-14 |
| ko00030; Pentose phosphate pathway | 2.42E-08 | 8.26E-08 | Day03-07 | Day03-07 - Day08-14 |
| ko00480; Glutathione metabolism | 1.67E-08 | 8.26E-08 | Day03-07 | Day03-07 - Day08-14 |
| ko00540; Lipopolysaccharide biosynthesis | 1.63E-08 | 8.26E-08 | Day03-07 | Day03-07 - Day08-14 |
| ko00562; Inositol phosphate metabolism | 2.73E-08 | 8.26E-08 | Day03-07 | Day03-07 - Day08-14 |
| ko00901; Indole alkaloid biosynthesis | 6.79E-09 | 8.26E-08 | Day03-07 | Day03-07 - Day08-14 |
| ko00965; Betalain biosynthesis | 2.04E-08 | 8.26E-08 | Day03-07 | Day03-07 - Day08-14 |
| ko00982; Drug metabolism - cytochrome P450 | 3.01E-08 | 8.26E-08 | Day03-07 | Day03-07 - Day08-14 |
| ko01040; Biosynthesis of unsaturated fatty acids | 1.15E-08 | 8.26E-08 | Day03-07 | Day03-07 - Day08-14 |
| ko05100; Bacterial invasion of epithelial cells | 2.94E-08 | 8.26E-08 | Day03-07 | Day03-07 - Day08-14 |
| ko05204; Chemical carcinogenesis | 1.21E-08 | 8.26E-08 | Day03-07 | Day03-07 - Day08-14 |
| ko05340; Primary immunodeficiency | 1.84E-08 | 8.26E-08 | Day03-07 | Day03-07 - Day08-14 |
| ko00564; Glycerophospholipid metabolism | 3.16E-08 | 1.95E-07 | Day03-07 | Day03-07 - Day08-14 |
| ko00980; Metabolism of xenobiotics by cytochrome P450 | 4.43E-08 | 1.95E-07 | Day03-07 | Day03-07 - Day08-14 |
| ko03010; Ribosome | 4.02E-08 | 1.95E-07 | Day08-14 | Day03-07 - Day08-14 |
| ko00196; Photosynthesis - antenna proteins | 6.04E-08 | 3.54E-07 | Day03-07 | Day03-07 - Day08-14 |
| ko05142; Chagas disease (American trypanosomiasis) | 4.76E-08 | 3.54E-07 | Day03-07 | Day03-07 - Day08-14 |
| ko05030; Cocaine addiction | 7.66E-08 | 6.61E-07 | Day03-07 | Day03-07 - Day08-14 |
| ko05031; Amphetamine addiction | 7.66E-08 | 6.61E-07 | Day03-07 | Day03-07 - Day08-14 |
| ko00310; Lysine degradation | 1.85E-07 | 1.40E-06 | Day03-07 | Day03-07 - Day08-14 |
| ko00660; C5-Branched dibasic acid metabolism | 2.73E-07 | 1.40E-06 | Day03-07 | Day03-07 - Day08-14 |
| ko00945; Stilbenoid, diarylheptanoid and gingerol biosynthesis | 9.46E-08 | 1.40E-06 | Day03-07 | Day03-07 - Day08-14 |
| ko04728; Dopaminergic synapse | 2.03E-07 | 1.40E-06 | Day03-07 | Day03-07 - Day08-14 |
| ko05012; Parkinsons disease | 2.79E-07 | 1.40E-06 | Day03-07 | Day03-07 - Day08-14 |
| ko00071; Fatty acid metabolism | 1.15E-06 | 4.07E-06 | Day03-07 | Day03-07 - Day08-14 |
| ko00232; Caffeine metabolism | 7.11E-07 | 4.07E-06 | Day03-07 | Day03-07 - Day08-14 |
| ko00363; Bisphenol degradation | 1.68E-06 | 4.07E-06 | Day03-07 | Day03-07 - Day08-14 |
| ko00471; D-Glutamine and D-glutamate metabolism | 1.51E-06 | 4.07E-06 | Day08-14 | Day03-07 - Day08-14 |
| ko00601; Glycosphingolipid biosynthesis - lacto and neolacto series | 9.04E-07 | 4.07E-06 | Day08-14 | Day03-07 - Day08-14 |
| ko00642; Ethylbenzene degradation | 2.86E-07 | 4.07E-06 | Day08-14 | Day03-07 - Day08-14 |
| ko00680; Methane metabolism | 1.42E-06 | 4.07E-06 | Day08-14 | Day03-07 - Day08-14 |
| ko00830; Retinol metabolism | 1.68E-06 | 4.07E-06 | Day03-07 | Day03-07 - Day08-14 |
| ko04727; GABAergic synapse | 1.15E-06 | 4.07E-06 | Day08-14 | Day03-07 - Day08-14 |
| ko04940; Type I diabetes mellitus | 4.09E-07 | 4.07E-06 | Day08-14 | Day03-07 - Day08-14 |
| ko05032; Morphine addiction | 1.05E-06 | 4.07E-06 | Day03-07 | Day03-07 - Day08-14 |
| ko05130; Pathogenic Escherichia coli infection | 9.64E-07 | 4.07E-06 | Day03-07 | Day03-07 - Day08-14 |
| ko05164; Influenza A | 1.76E-06 | 4.07E-06 | Day08-14 | Day03-07 - Day08-14 |
| ko05203; Viral carcinogenesis | 1.83E-06 | 4.07E-06 | Day03-07 | Day03-07 - Day08-14 |
| ko00740; Riboflavin metabolism | 1.95E-06 | 4.65E-06 | Day03-07 | Day03-07 - Day08-14 |
| ko04724; Glutamatergic synapse | 1.95E-06 | 4.65E-06 | Day08-14 | Day03-07 - Day08-14 |
| ko00630; Glyoxylate and dicarboxylate metabolism | 2.08E-06 | 6.01E-06 | Day03-07 | Day03-07 - Day08-14 |
| ko00640; Propanoate metabolism | 2.26E-06 | 6.01E-06 | Day03-07 | Day03-07 - Day08-14 |
| ko04930; Type II diabetes mellitus | 2.41E-06 | 1.82E-05 | Day03-07 | Day03-07 - Day08-14 |
| ko00603; Glycosphingolipid biosynthesis - globo series | 4.56E-06 | 2.31E-05 | Day08-14 | Day03-07 - Day08-14 |
| ko00860; Porphyrin and chlorophyll metabolism | 5.93E-06 | 2.31E-05 | Day03-07 | Day03-07 - Day08-14 |
| ko04146; Peroxisome | 4.65E-06 | 2.31E-05 | Day03-07 | Day03-07 - Day08-14 |
| ko04910; Insulin signaling pathway | 2.90E-06 | 2.31E-05 | Day03-07 | Day03-07 - Day08-14 |
| ko05168; Herpes simplex infection | 5.58E-06 | 2.31E-05 | Day08-14 | Day03-07 - Day08-14 |
| ko00130; Ubiquinone and other terpenoid-quinone biosynthesis | 6.05E-06 | 2.46E-05 | Day03-07 | Day03-07 - Day08-14 |
| ko04070; Phosphatidylinositol signaling system | 6.96E-06 | 2.46E-05 | Day03-07 | Day03-07 - Day08-14 |
| ko00312; beta-Lactam resistance | 1.14E-05 | 2.59E-05 | Day08-14 | Day03-07 - Day08-14 |
| ko00350; Tyrosine metabolism | 9.76E-06 | 2.59E-05 | Day08-14 | Day03-07 - Day08-14 |
| ko00400; Phenylalanine, tyrosine and tryptophan biosynthesis | 1.39E-05 | 2.59E-05 | Day08-14 | Day03-07 - Day08-14 |
| ko00623; Toluene degradation | 1.56E-05 | 2.59E-05 | Day03-07 | Day03-07 - Day08-14 |
| ko00710; Carbon fixation in photosynthetic organisms | 1.44E-05 | 2.59E-05 | Day08-14 | Day03-07 - Day08-14 |
| ko01057; Biosynthesis of type II polyketide products | 1.33E-05 | 2.59E-05 | Day03-07 | Day03-07 - Day08-14 |
| ko02060; Phosphotransferase system (PTS) | 7.39E-06 | 2.59E-05 | Day03-07 | Day03-07 - Day08-14 |
| ko03008; Ribosome biogenesis in eukaryotes | 1.47E-05 | 2.59E-05 | Day03-07 | Day03-07 - Day08-14 |
| ko03018; RNA degradation | 1.50E-05 | 2.59E-05 | Day03-07 | Day03-07 - Day08-14 |
| ko03060; Protein export | 1.44E-05 | 2.59E-05 | Day08-14 | Day03-07 - Day08-14 |
| ko05210; Colorectal cancer | 1.16E-05 | 2.59E-05 | Day08-14 | Day03-07 - Day08-14 |
| ko05416; Viral myocarditis | 1.16E-05 | 2.59E-05 | Day08-14 | Day03-07 - Day08-14 |
| ko00061; Fatty acid biosynthesis | 1.59E-05 | 3.45E-05 | Day03-07 | Day03-07 - Day08-14 |
| ko00100; Steroid biosynthesis | 1.68E-05 | 8.68E-05 | Day03-07 | Day03-07 - Day08-14 |
| ko00340; Histidine metabolism | 1.96E-05 | 8.68E-05 | Day08-14 | Day03-07 - Day08-14 |
| ko00401; Novobiocin biosynthesis | 2.16E-05 | 0.000109709 | Day08-14 | Day03-07 - Day08-14 |
| ko00621; Dioxin degradation | 2.51E-05 | 0.000109709 | Day08-14 | Day03-07 - Day08-14 |
| ko00010; Glycolysis / Gluconeogenesis | 6.52E-05 | 0.000145793 | Day03-07 | Day03-07 - Day08-14 |
| ko00053; Ascorbate and aldarate metabolism | 0.000110879 | 0.000145793 | Day03-07 | Day03-07 - Day08-14 |
| ko00140; Steroid hormone biosynthesis | 2.75E-05 | 0.000145793 | Day03-07 | Day03-07 - Day08-14 |
| ko00361; Chlorocyclohexane and chlorobenzene degradation | 7.13E-05 | 0.000145793 | Day03-07 | Day03-07 - Day08-14 |
| ko00430; Taurine and hypotaurine metabolism | 0.000120947 | 0.000145793 | Day03-07 | Day03-07 - Day08-14 |
| ko00524; Butirosin and neomycin biosynthesis | 9.31E-05 | 0.000145793 | Day03-07 | Day03-07 - Day08-14 |
| ko00565; Ether lipid metabolism | 8.52E-05 | 0.000145793 | Day03-07 | Day03-07 - Day08-14 |
| ko00970; Aminoacyl-tRNA biosynthesis | 0.000114808 | 0.000145793 | Day08-14 | Day03-07 - Day08-14 |
| ko02040; Flagellar assembly | 3.26E-05 | 0.000145793 | Day03-07 | Day03-07 - Day08-14 |
| ko03030; DNA replication | 0.000107078 | 0.000145793 | Day08-14 | Day03-07 - Day08-14 |
| ko03450; Non-homologous end-joining | 7.53E-05 | 0.000145793 | Day03-07 | Day03-07 - Day08-14 |
| ko04310; Wnt signaling pathway | 8.98E-05 | 0.000145793 | Day08-14 | Day03-07 - Day08-14 |
| ko04330; Notch signaling pathway | 8.98E-05 | 0.000145793 | Day08-14 | Day03-07 - Day08-14 |
| ko04514; Cell adhesion molecules (CAMs) | 6.25E-05 | 0.000145793 | Day08-14 | Day03-07 - Day08-14 |
| ko04640; Hematopoietic cell lineage | 6.76E-05 | 0.000145793 | Day08-14 | Day03-07 - Day08-14 |
| ko04721; Synaptic vesicle cycle | 0.000108963 | 0.000145793 | Day08-14 | Day03-07 - Day08-14 |
| ko04725; Cholinergic synapse | 6.41E-05 | 0.000145793 | Day03-07 | Day03-07 - Day08-14 |
| ko04810; Regulation of actin cytoskeleton | 0.000118694 | 0.000145793 | Day08-14 | Day03-07 - Day08-14 |
| ko04962; Vasopressin-regulated water reabsorption | 0.000108963 | 0.000145793 | Day08-14 | Day03-07 - Day08-14 |
| ko04975; Fat digestion and absorption | 8.22E-05 | 0.000145793 | Day08-14 | Day03-07 - Day08-14 |
| ko05202; Transcriptional misregulation in cancer | 9.32E-05 | 0.000145793 | Day08-14 | Day03-07 - Day08-14 |
| ko05220; Chronic myeloid leukemia | 8.98E-05 | 0.000145793 | Day08-14 | Day03-07 - Day08-14 |
| ko05412; Arrhythmogenic right ventricular cardiomyopathy (ARVC) | 6.25E-05 | 0.000145793 | Day08-14 | Day03-07 - Day08-14 |
| ko05414; Dilated cardiomyopathy | 6.25E-05 | 0.000145793 | Day08-14 | Day03-07 - Day08-14 |
| ko00450; Selenocompound metabolism | 0.000131872 | 0.000236604 | Day03-07 | Day03-07 - Day08-14 |
| ko00604; Glycosphingolipid biosynthesis - ganglio series | 0.000123061 | 0.000236604 | Day08-14 | Day03-07 - Day08-14 |
| ko00240; Pyrimidine metabolism | 0.000215926 | 0.000470296 | Day08-14 | Day03-07 - Day08-14 |
| ko01055; Biosynthesis of vancomycin group antibiotics | 0.000219572 | 0.000470296 | Day08-14 | Day03-07 - Day08-14 |
| ko02010; ABC transporters | 0.00017638 | 0.000470296 | Day03-07 | Day03-07 - Day08-14 |
| ko04064; NF-kappa B signaling pathway | 0.000233992 | 0.000470296 | Day08-14 | Day03-07 - Day08-14 |
| ko04112; Cell cycle - Caulobacter | 0.000191949 | 0.000470296 | Day08-14 | Day03-07 - Day08-14 |
| ko04626; Plant-pathogen interaction | 0.000167617 | 0.000470296 | Day03-07 | Day03-07 - Day08-14 |
| ko04916; Melanogenesis | 0.000135979 | 0.000470296 | Day08-14 | Day03-07 - Day08-14 |
| ko05140; Leishmaniasis | 0.000233992 | 0.000470296 | Day08-14 | Day03-07 - Day08-14 |
| ko00253; Tetracycline biosynthesis | 0.000277087 | 0.000568971 | Day03-07 | Day03-07 - Day08-14 |
| ko00785; Lipoic acid metabolism | 0.000300849 | 0.000568971 | Day03-07 | Day03-07 - Day08-14 |
| ko03070; Bacterial secretion system | 0.000268084 | 0.000568971 | Day03-07 | Day03-07 - Day08-14 |
| ko03440; Homologous recombination | 0.000238688 | 0.000568971 | Day08-14 | Day03-07 - Day08-14 |
| ko04080; Neuroactive ligand-receptor interaction | 0.000292503 | 0.000568971 | Day08-14 | Day03-07 - Day08-14 |
| ko04622; RIG-I-like receptor signaling pathway | 0.000321219 | 0.000568971 | Day03-07 | Day03-07 - Day08-14 |
| ko00300; Lysine biosynthesis | 0.000331881 | 0.000834975 | Day08-14 | Day03-07 - Day08-14 |
| ko00364; Fluorobenzoate degradation | 0.00038409 | 0.000834975 | Day03-07 | Day03-07 - Day08-14 |
| ko00720; Carbon fixation pathways in prokaryotes | 0.000360004 | 0.000834975 | Day08-14 | Day03-07 - Day08-14 |
| ko05169; Epstein-Barr virus infection | 0.000419343 | 0.000834975 | Day03-07 | Day03-07 - Day08-14 |
| ko04141; Protein processing in endoplasmic reticulum | 0.000423057 | 0.002396589 | Day08-14 | Day03-07 - Day08-14 |
| ko00460; Cyanoamino acid metabolism | 0.000590446 | 0.002834065 | Day03-07 | Day03-07 - Day08-14 |
| ko00523; Polyketide sugar unit biosynthesis | 0.000700997 | 0.002834065 | Day08-14 | Day03-07 - Day08-14 |
| ko04122; Sulfur relay system | 0.00074576 | 0.002834065 | Day08-14 | Day03-07 - Day08-14 |
| ko05016; Huntingtons disease | 0.000856352 | 0.002834065 | Day03-07 | Day03-07 - Day08-14 |
| ko05134; Legionellosis | 0.000488481 | 0.002834065 | Day08-14 | Day03-07 - Day08-14 |
| ko05200; Pathways in cancer | 0.000896474 | 0.002834065 | Day08-14 | Day03-07 - Day08-14 |
| ko00250; Alanine, aspartate and glutamate metabolism | 0.001324644 | 0.005498727 | Day08-14 | Day03-07 - Day08-14 |
| ko00730; Thiamine metabolism | 0.001364383 | 0.005498727 | Day08-14 | Day03-07 - Day08-14 |
| ko00940; Phenylpropanoid biosynthesis | 0.001826811 | 0.005498727 | Day03-07 | Day03-07 - Day08-14 |
| ko05120; Epithelial cell signaling in Helicobacter pylori infection | 0.001468591 | 0.005498727 | Day03-07 | Day03-07 - Day08-14 |
| ko05211; Renal cell carcinoma | 0.001774802 | 0.005498727 | Day08-14 | Day03-07 - Day08-14 |
| ko05222; Small cell lung cancer | 0.000952682 | 0.005498727 | Day08-14 | Day03-07 - Day08-14 |
| ko00020; Citrate cycle (TCA cycle) | 0.001853337 | 0.005535155 | Day03-07 | Day03-07 - Day08-14 |
| ko00195; Photosynthesis | 0.006251038 | 0.005535155 | Day08-14 | Day03-07 - Day08-14 |
| ko00330; Arginine and proline metabolism | 0.002463965 | 0.005535155 | Day08-14 | Day03-07 - Day08-14 |
| ko00351; DDT degradation | 0.00279471 | 0.005535155 | Day08-14 | Day03-07 - Day08-14 |
| ko00472; D-Arginine and D-ornithine metabolism | 0.003344319 | 0.005535155 | Day03-07 | Day03-07 - Day08-14 |
| ko00561; Glycerolipid metabolism | 0.002913663 | 0.005535155 | Day03-07 | Day03-07 - Day08-14 |
| ko00591; Linoleic acid metabolism | 0.00481191 | 0.005535155 | Day03-07 | Day03-07 - Day08-14 |
| ko00600; Sphingolipid metabolism | 0.003390427 | 0.005535155 | Day08-14 | Day03-07 - Day08-14 |
| ko00620; Pyruvate metabolism | 0.00415511 | 0.005535155 | Day03-07 | Day03-07 - Day08-14 |
| ko00622; Xylene degradation | 0.00514041 | 0.005535155 | Day08-14 | Day03-07 - Day08-14 |
| ko00624; Polycyclic aromatic hydrocarbon degradation | 0.003209399 | 0.005535155 | Day08-14 | Day03-07 - Day08-14 |
| ko00633; Nitrotoluene degradation | 0.006581604 | 0.005535155 | Day03-07 | Day03-07 - Day08-14 |
| ko00750; Vitamin B6 metabolism | 0.004268115 | 0.005535155 | Day08-14 | Day03-07 - Day08-14 |
| ko00770; Pantothenate and CoA biosynthesis | 0.002395511 | 0.005535155 | Day08-14 | Day03-07 - Day08-14 |
| ko00906; Carotenoid biosynthesis | 0.004211269 | 0.005535155 | Day08-14 | Day03-07 - Day08-14 |
| ko01051; Biosynthesis of ansamycins | 0.002498859 | 0.005535155 | Day03-07 | Day03-07 - Day08-14 |
| ko03015; mRNA surveillance pathway | 0.006251038 | 0.005535155 | Day03-07 | Day03-07 - Day08-14 |
| ko03050; Proteasome | 0.003580706 | 0.005535155 | Day08-14 | Day03-07 - Day08-14 |
| ko04020; Calcium signaling pathway | 0.004502523 | 0.005535155 | Day08-14 | Day03-07 - Day08-14 |
| ko04114; Oocyte meiosis | 0.006799255 | 0.005535155 | Day08-14 | Day03-07 - Day08-14 |
| ko04270; Vascular smooth muscle contraction | 0.006799255 | 0.005535155 | Day08-14 | Day03-07 - Day08-14 |
| ko04370; VEGF signaling pathway | 0.002913663 | 0.005535155 | Day08-14 | Day03-07 - Day08-14 |
| ko04666; Fc gamma R-mediated phagocytosis | 0.004099632 | 0.005535155 | Day08-14 | Day03-07 - Day08-14 |
| ko04720; Long-term potentiation | 0.006799255 | 0.005535155 | Day08-14 | Day03-07 - Day08-14 |
| ko04722; Neurotrophin signaling pathway | 0.006799255 | 0.005535155 | Day08-14 | Day03-07 - Day08-14 |
| ko04740; Olfactory transduction | 0.006799255 | 0.005535155 | Day08-14 | Day03-07 - Day08-14 |
| ko04744; Phototransduction | 0.006799255 | 0.005535155 | Day08-14 | Day03-07 - Day08-14 |
| ko04745; Phototransduction - fly | 0.006799255 | 0.005535155 | Day08-14 | Day03-07 - Day08-14 |
| ko05150; Staphylococcus aureus infection | 0.002078632 | 0.005535155 | Day08-14 | Day03-07 - Day08-14 |
| ko05152; Tuberculosis | 0.006332247 | 0.005535155 | Day08-14 | Day03-07 - Day08-14 |
| ko05160; Hepatitis C | 0.005480406 | 0.005535155 | Day08-14 | Day03-07 - Day08-14 |
| ko05162; Measles | 0.005480406 | 0.005535155 | Day08-14 | Day03-07 - Day08-14 |
| ko05214; Glioma | 0.006799255 | 0.005535155 | Day08-14 | Day03-07 - Day08-14 |
| ko00040; Pentose and glucuronate interconversions | 0.007965754 | 0.009152701 | Day03-07 | Day03-07 - Day08-14 |
| ko00473; D-Alanine metabolism | 0.007865947 | 0.009152701 | Day08-14 | Day03-07 - Day08-14 |
| ko00960; Tropane, piperidine and pyridine alkaloid biosynthesis | 0.006839757 | 0.009152701 | Day03-07 | Day03-07 - Day08-14 |
| ko04961; Endocrine and other factor-regulated calcium reabsorption | 0.007965754 | 0.009152701 | Day08-14 | Day03-07 - Day08-14 |
| ko04972; Pancreatic secretion | 0.007290285 | 0.009152701 | Day08-14 | Day03-07 - Day08-14 |
| ko00052; Galactose metabolism | 0.014999488 | 0.030178743 | Day03-07 | Day03-07 - Day08-14 |
| ko00190; Oxidative phosphorylation | 0.012553962 | 0.030178743 | Day08-14 | Day03-07 - Day08-14 |
| ko00670; One carbon pool by folate | 0.016468133 | 0.030178743 | Day08-14 | Day03-07 - Day08-14 |
| ko00780; Biotin metabolism | 0.016660318 | 0.030178743 | Day08-14 | Day03-07 - Day08-14 |
| ko02030; Bacterial chemotaxis | 0.017448893 | 0.030178743 | Day08-14 | Day03-07 - Day08-14 |
| ko03410; Base excision repair | 0.016089595 | 0.030178743 | Day03-07 | Day03-07 - Day08-14 |
| ko04210; Apoptosis | 0.013486272 | 0.030178743 | Day08-14 | Day03-07 - Day08-14 |
| ko04612; Antigen processing and presentation | 0.014309398 | 0.030178743 | Day03-07 | Day03-07 - Day08-14 |
| ko04914; Progesterone-mediated oocyte maturation | 0.014309398 | 0.030178743 | Day03-07 | Day03-07 - Day08-14 |
| ko04960; Aldosterone-regulated sodium reabsorption | 0.008588768 | 0.030178743 | Day08-14 | Day03-07 - Day08-14 |
| ko04970; Salivary secretion | 0.008588768 | 0.030178743 | Day08-14 | Day03-07 - Day08-14 |
| ko04971; Gastric acid secretion | 0.008588768 | 0.030178743 | Day08-14 | Day03-07 - Day08-14 |
| ko05215; Prostate cancer | 0.014309398 | 0.030178743 | Day03-07 | Day03-07 - Day08-14 |
| ko05014; Amyotrophic lateral sclerosis (ALS) | 3.10E-09 | 2.80E-07 | Day08-14 | Day08-14 - Day15-24 |
| ko03050; Proteasome | 4.41E-09 | 3.83E-07 | Day15-24 | Day08-14 - Day15-24 |
| ko00331; Clavulanic acid biosynthesis | 7.43E-09 | 4.28E-07 | Day08-14 | Day08-14 - Day15-24 |
| ko00592; alpha-Linolenic acid metabolism | 1.77E-08 | 4.28E-07 | Day08-14 | Day08-14 - Day15-24 |
| ko04723; Retrograde endocannabinoid signaling | 1.34E-08 | 4.28E-07 | Day08-14 | Day08-14 - Day15-24 |
| ko05020; Prion diseases | 1.56E-08 | 4.28E-07 | Day08-14 | Day08-14 - Day15-24 |
| ko04974; Protein digestion and absorption | 2.09E-08 | 3.33E-06 | Day15-24 | Day08-14 - Day15-24 |
| ko00380; Tryptophan metabolism | 4.72E-08 | 4.96E-06 | Day08-14 | Day08-14 - Day15-24 |
| ko00400; Phenylalanine, tyrosine and tryptophan biosynthesis | 5.90E-07 | 1.25E-05 | Day15-24 | Day08-14 - Day15-24 |
| ko03040; Spliceosome | 7.74E-07 | 1.25E-05 | Day08-14 | Day08-14 - Day15-24 |
| ko04514; Cell adhesion molecules (CAMs) | 7.77E-07 | 1.25E-05 | Day15-24 | Day08-14 - Day15-24 |
| ko04610; Complement and coagulation cascades | 2.49E-07 | 1.25E-05 | Day08-14 | Day08-14 - Day15-24 |
| ko04724; Glutamatergic synapse | 9.61E-07 | 1.25E-05 | Day15-24 | Day08-14 - Day15-24 |
| ko04810; Regulation of actin cytoskeleton | 6.82E-07 | 1.25E-05 | Day15-24 | Day08-14 - Day15-24 |
| ko05131; Shigellosis | 8.63E-08 | 1.25E-05 | Day08-14 | Day08-14 - Day15-24 |
| ko05412; Arrhythmogenic right ventricular cardiomyopathy (ARVC) | 7.77E-07 | 1.25E-05 | Day15-24 | Day08-14 - Day15-24 |
| ko05414; Dilated cardiomyopathy | 7.77E-07 | 1.25E-05 | Day15-24 | Day08-14 - Day15-24 |
| ko00520; Amino sugar and nucleotide sugar metabolism | 9.74E-07 | 7.45E-05 | Day15-24 | Day08-14 - Day15-24 |
| ko00471; D-Glutamine and D-glutamate metabolism | 4.59E-06 | 0.000191679 | Day15-24 | Day08-14 - Day15-24 |
| ko04916; Melanogenesis | 1.56E-06 | 0.000191679 | Day15-24 | Day08-14 - Day15-24 |
| ko05202; Transcriptional misregulation in cancer | 4.37E-06 | 0.000191679 | Day15-24 | Day08-14 - Day15-24 |
| ko04080; Neuroactive ligand-receptor interaction | 1.26E-05 | 0.000351318 | Day15-24 | Day08-14 - Day15-24 |
| ko04975; Fat digestion and absorption | 1.26E-05 | 0.000351318 | Day15-24 | Day08-14 - Day15-24 |
| ko05169; Epstein-Barr virus infection | 6.10E-06 | 0.000351318 | Day15-24 | Day08-14 - Day15-24 |
| ko00232; Caffeine metabolism | 7.33E-05 | 0.000771811 | Day08-14 | Day08-14 - Day15-24 |
| ko00253; Tetracycline biosynthesis | 4.09E-05 | 0.000771811 | Day08-14 | Day08-14 - Day15-24 |
| ko00401; Novobiocin biosynthesis | 7.75E-05 | 0.000771811 | Day15-24 | Day08-14 - Day15-24 |
| ko00500; Starch and sucrose metabolism | 6.78E-05 | 0.000771811 | Day15-24 | Day08-14 - Day15-24 |
| ko00513; Various types of N-glycan biosynthesis | 1.44E-05 | 0.000771811 | Day15-24 | Day08-14 - Day15-24 |
| ko00623; Toluene degradation | 7.75E-05 | 0.000771811 | Day08-14 | Day08-14 - Day15-24 |
| ko01053; Biosynthesis of siderophore group nonribosomal peptides | 6.49E-05 | 0.000771811 | Day08-14 | Day08-14 - Day15-24 |
| ko01057; Biosynthesis of type II polyketide products | 5.80E-05 | 0.000771811 | Day08-14 | Day08-14 - Day15-24 |
| ko03010; Ribosome | 5.13E-05 | 0.000771811 | Day15-24 | Day08-14 - Day15-24 |
| ko04640; Hematopoietic cell lineage | 2.40E-05 | 0.000771811 | Day15-24 | Day08-14 - Day15-24 |
| ko05100; Bacterial invasion of epithelial cells | 2.93E-05 | 0.000771811 | Day08-14 | Day08-14 - Day15-24 |
| ko00310; Lysine degradation | 0.000145273 | 0.001828317 | Day08-14 | Day08-14 - Day15-24 |
| ko00603; Glycosphingolipid biosynthesis - globo series | 8.74E-05 | 0.001828317 | Day15-24 | Day08-14 - Day15-24 |
| ko00984; Steroid degradation | 0.000156489 | 0.001828317 | Day08-14 | Day08-14 - Day15-24 |
| ko01055; Biosynthesis of vancomycin group antibiotics | 8.74E-05 | 0.001828317 | Day15-24 | Day08-14 - Day15-24 |
| ko05130; Pathogenic Escherichia coli infection | 0.000136266 | 0.001828317 | Day08-14 | Day08-14 - Day15-24 |
| ko05164; Influenza A | 0.000161541 | 0.001828317 | Day15-24 | Day08-14 - Day15-24 |
| ko05204; Chemical carcinogenesis | 0.000163258 | 0.001828317 | Day08-14 | Day08-14 - Day15-24 |
| ko00363; Bisphenol degradation | 0.000173932 | 0.002295258 | Day08-14 | Day08-14 - Day15-24 |
| ko00473; D-Alanine metabolism | 0.000221233 | 0.002295258 | Day15-24 | Day08-14 - Day15-24 |
| ko00901; Indole alkaloid biosynthesis | 0.000203556 | 0.002295258 | Day08-14 | Day08-14 - Day15-24 |
| ko00051; Fructose and mannose metabolism | 0.000564731 | 0.009251318 | Day15-24 | Day08-14 - Day15-24 |
| ko00531; Glycosaminoglycan degradation | 0.000228226 | 0.009251318 | Day15-24 | Day08-14 - Day15-24 |
| ko03015; mRNA surveillance pathway | 0.00050698 | 0.009251318 | Day08-14 | Day08-14 - Day15-24 |
| ko04727; GABAergic synapse | 0.000323394 | 0.009251318 | Day15-24 | Day08-14 - Day15-24 |
| ko05111; Vibrio cholerae pathogenic cycle | 0.000383863 | 0.009251318 | Day08-14 | Day08-14 - Day15-24 |
| ko04064; NF-kappa B signaling pathway | 0.001256247 | 0.017059387 | Day15-24 | Day08-14 - Day15-24 |
| ko04940; Type I diabetes mellitus | 0.000887451 | 0.017059387 | Day15-24 | Day08-14 - Day15-24 |
| ko05110; Vibrio cholerae infection | 0.001041361 | 0.017059387 | Day08-14 | Day08-14 - Day15-24 |
| ko05140; Leishmaniasis | 0.001256247 | 0.017059387 | Day15-24 | Day08-14 - Day15-24 |
| ko05145; Toxoplasmosis | 0.000592958 | 0.017059387 | Day15-24 | Day08-14 - Day15-24 |
| ko00982; Drug metabolism - cytochrome P450 | 0.001465349 | 0.017880782 | Day08-14 | Day08-14 - Day15-24 |
| ko05133; Pertussis | 0.001265523 | 0.017880782 | Day08-14 | Day08-14 - Day15-24 |
| ko00590; Arachidonic acid metabolism | 0.001547502 | 0.018567059 | Day08-14 | Day08-14 - Day15-24 |
| ko00071; Fatty acid metabolism | 0.001954847 | 0.020888313 | Day08-14 | Day08-14 - Day15-24 |
| ko00600; Sphingolipid metabolism | 0.0025238 | 0.020888313 | Day15-24 | Day08-14 - Day15-24 |
| ko00627; Aminobenzoate degradation | 0.002501893 | 0.020888313 | Day08-14 | Day08-14 - Day15-24 |
| ko00790; Folate biosynthesis | 0.00248016 | 0.020888313 | Day15-24 | Day08-14 - Day15-24 |
| ko00980; Metabolism of xenobiotics by cytochrome P450 | 0.001693903 | 0.020888313 | Day08-14 | Day08-14 - Day15-24 |
| ko05012; Parkinsons disease | 0.002545882 | 0.020888313 | Day08-14 | Day08-14 - Day15-24 |
| ko00965; Betalain biosynthesis | 0.002682127 | 0.024353449 | Day08-14 | Day08-14 - Day15-24 |
| ko00960; Tropane, piperidine and pyridine alkaloid biosynthesis | 0.003468705 | 0.045274535 | Day08-14 | Day08-14 - Day15-24 |
| ko05132; Salmonella infection | 0.002874153 | 0.045274535 | Day08-14 | Day08-14 - Day15-24 |
| ko00340; Histidine metabolism | 1.31E-09 | 1.21E-07 | Day15-24 | Day15-24 - Day25-35 |
| ko00030; Pentose phosphate pathway | 2.08E-08 | 1.16E-06 | Day25-35 | Day15-24 - Day25-35 |
| ko04622; RIG-I-like receptor signaling pathway | 2.16E-09 | 1.16E-06 | Day25-35 | Day15-24 - Day25-35 |
| ko05168; Herpes simplex infection | 2.35E-08 | 1.16E-06 | Day15-24 | Day15-24 - Day25-35 |
| ko00710; Carbon fixation in photosynthetic organisms | 3.31E-08 | 1.46E-06 | Day25-35 | Day15-24 - Day25-35 |
| ko00790; Folate biosynthesis | 6.64E-08 | 1.46E-06 | Day25-35 | Day15-24 - Day25-35 |
| ko00010; Glycolysis / Gluconeogenesis | 1.08E-07 | 3.74E-06 | Day25-35 | Day15-24 - Day25-35 |
| ko00521; Streptomycin biosynthesis | 2.34E-07 | 3.74E-06 | Day25-35 | Day15-24 - Day25-35 |
| ko04112; Cell cycle - Caulobacter | 1.82E-07 | 3.74E-06 | Day15-24 | Day15-24 - Day25-35 |
| ko04115; p53 signaling pathway | 1.82E-07 | 3.74E-06 | Day15-24 | Day15-24 - Day25-35 |
| ko05164; Influenza A | 2.56E-07 | 3.74E-06 | Day25-35 | Day15-24 - Day25-35 |
| ko05210; Colorectal cancer | 7.21E-08 | 3.74E-06 | Day15-24 | Day15-24 - Day25-35 |
| ko05416; Viral myocarditis | 7.21E-08 | 3.74E-06 | Day15-24 | Day15-24 - Day25-35 |
| ko02030; Bacterial chemotaxis | 2.72E-07 | 1.40E-05 | Day15-24 | Day15-24 - Day25-35 |
| ko04210; Apoptosis | 5.01E-07 | 1.40E-05 | Day15-24 | Day15-24 - Day25-35 |
| ko00472; D-Arginine and D-ornithine metabolism | 5.21E-07 | 3.11E-05 | Day25-35 | Day15-24 - Day25-35 |
| ko00941; Flavonoid biosynthesis | 7.97E-07 | 3.20E-05 | Day15-24 | Day15-24 - Day25-35 |
| ko00312; beta-Lactam resistance | 5.76E-06 | 0.000163818 | Day15-24 | Day15-24 - Day25-35 |
| ko00523; Polyketide sugar unit biosynthesis | 2.65E-06 | 0.000163818 | Day25-35 | Day15-24 - Day25-35 |
| ko02020; Two-component system | 1.25E-05 | 0.000163818 | Day15-24 | Day15-24 - Day25-35 |
| ko02060; Phosphotransferase system (PTS) | 1.08E-06 | 0.000163818 | Day25-35 | Day15-24 - Day25-35 |
| ko03010; Ribosome | 1.17E-05 | 0.000163818 | Day25-35 | Day15-24 - Day25-35 |
| ko04940; Type I diabetes mellitus | 6.13E-06 | 0.000163818 | Day25-35 | Day15-24 - Day25-35 |
| ko05219; Bladder cancer | 1.06E-05 | 0.000163818 | Day25-35 | Day15-24 - Day25-35 |
| ko05222; Small cell lung cancer | 7.00E-06 | 0.000163818 | Day15-24 | Day15-24 - Day25-35 |
| ko00052; Galactose metabolism | 2.38E-05 | 0.000392037 | Day25-35 | Day15-24 - Day25-35 |
| ko00130; Ubiquinone and other terpenoid-quinone biosynthesis | 1.27E-05 | 0.000392037 | Day25-35 | Day15-24 - Day25-35 |
| ko00290; Valine, leucine and isoleucine biosynthesis | 3.03E-05 | 0.000392037 | Day15-24 | Day15-24 - Day25-35 |
| ko03450; Non-homologous end-joining | 2.70E-05 | 0.000392037 | Day25-35 | Day15-24 - Day25-35 |
| ko04070; Phosphatidylinositol signaling system | 1.69E-05 | 0.000392037 | Day25-35 | Day15-24 - Day25-35 |
| ko05146; Amoebiasis | 2.79E-05 | 0.000392037 | Day15-24 | Day15-24 - Day25-35 |
| ko00604; Glycosphingolipid biosynthesis - ganglio series | 5.21E-05 | 0.001884939 | Day15-24 | Day15-24 - Day25-35 |
| ko04930; Type II diabetes mellitus | 3.37E-05 | 0.001884939 | Day25-35 | Day15-24 - Day25-35 |
| ko00053; Ascorbate and aldarate metabolism | 0.000196433 | 0.00197874 | Day25-35 | Day15-24 - Day25-35 |
| ko00072; Synthesis and degradation of ketone bodies | 0.000154622 | 0.00197874 | Day15-24 | Day15-24 - Day25-35 |
| ko00196; Photosynthesis - antenna proteins | 0.000150028 | 0.00197874 | Day25-35 | Day15-24 - Day25-35 |
| ko00350; Tyrosine metabolism | 0.000159347 | 0.00197874 | Day15-24 | Day15-24 - Day25-35 |
| ko00450; Selenocompound metabolism | 0.000179642 | 0.00197874 | Day25-35 | Day15-24 - Day25-35 |
| ko00471; D-Glutamine and D-glutamate metabolism | 8.71E-05 | 0.00197874 | Day25-35 | Day15-24 - Day25-35 |
| ko00909; Sesquiterpenoid and triterpenoid biosynthesis | 0.00021469 | 0.00197874 | Day15-24 | Day15-24 - Day25-35 |
| ko03430; Mismatch repair | 0.000109797 | 0.00197874 | Day15-24 | Day15-24 - Day25-35 |
| ko04974; Protein digestion and absorption | 6.72E-05 | 0.00197874 | Day25-35 | Day15-24 - Day25-35 |
| ko05142; Chagas disease (American trypanosomiasis) | 0.000241519 | 0.00197874 | Day25-35 | Day15-24 - Day25-35 |
| ko00524; Butirosin and neomycin biosynthesis | 0.000243296 | 0.002969885 | Day25-35 | Day15-24 - Day25-35 |
| ko04310; Wnt signaling pathway | 0.000347041 | 0.002969885 | Day15-24 | Day15-24 - Day25-35 |
| ko04330; Notch signaling pathway | 0.000347041 | 0.002969885 | Day15-24 | Day15-24 - Day25-35 |
| ko05203; Viral carcinogenesis | 0.000367481 | 0.002969885 | Day25-35 | Day15-24 - Day25-35 |
| ko05220; Chronic myeloid leukemia | 0.000347041 | 0.002969885 | Day15-24 | Day15-24 - Day25-35 |
| ko00061; Fatty acid biosynthesis | 0.000605033 | 0.004279392 | Day15-24 | Day15-24 - Day25-35 |
| ko00195; Photosynthesis | 0.000661672 | 0.004279392 | Day25-35 | Day15-24 - Day25-35 |
| ko00253; Tetracycline biosynthesis | 0.000375432 | 0.004279392 | Day15-24 | Day15-24 - Day25-35 |
| ko00540; Lipopolysaccharide biosynthesis | 0.00052298 | 0.004279392 | Day25-35 | Day15-24 - Day25-35 |
| ko00630; Glyoxylate and dicarboxylate metabolism | 0.000661672 | 0.004279392 | Day25-35 | Day15-24 - Day25-35 |
| ko01051; Biosynthesis of ansamycins | 0.000657146 | 0.004279392 | Day25-35 | Day15-24 - Day25-35 |
| ko03060; Protein export | 0.000605033 | 0.004279392 | Day25-35 | Day15-24 - Day25-35 |
| ko05322; Systemic lupus erythematosus | 0.000541518 | 0.004279392 | Day15-24 | Day15-24 - Day25-35 |
| ko00051; Fructose and mannose metabolism | 0.001580412 | 0.011322937 | Day25-35 | Day15-24 - Day25-35 |
| ko00230; Purine metabolism | 0.001388759 | 0.011322937 | Day25-35 | Day15-24 - Day25-35 |
| ko00281; Geraniol degradation | 0.000934912 | 0.011322937 | Day15-24 | Day15-24 - Day25-35 |
| ko00331; Clavulanic acid biosynthesis | 0.001663673 | 0.011322937 | Day25-35 | Day15-24 - Day25-35 |
| ko00351; DDT degradation | 0.001251127 | 0.011322937 | Day15-24 | Day15-24 - Day25-35 |
| ko00362; Benzoate degradation | 0.001962938 | 0.011322937 | Day15-24 | Day15-24 - Day25-35 |
| ko00562; Inositol phosphate metabolism | 0.001026433 | 0.011322937 | Day25-35 | Day15-24 - Day25-35 |
| ko00623; Toluene degradation | 0.002000478 | 0.011322937 | Day25-35 | Day15-24 - Day25-35 |
| ko01055; Biosynthesis of vancomycin group antibiotics | 0.000680068 | 0.011322937 | Day25-35 | Day15-24 - Day25-35 |
| ko01056; Biosynthesis of type II polyketide backbone | 0.002064511 | 0.011322937 | Day15-24 | Day15-24 - Day25-35 |
| ko02010; ABC transporters | 0.001111408 | 0.011322937 | Day25-35 | Day15-24 - Day25-35 |
| ko03008; Ribosome biogenesis in eukaryotes | 0.001762155 | 0.011322937 | Day25-35 | Day15-24 - Day25-35 |
| ko03020; RNA polymerase | 0.001621551 | 0.011322937 | Day25-35 | Day15-24 - Day25-35 |
| ko04975; Fat digestion and absorption | 0.001226767 | 0.011322937 | Day15-24 | Day15-24 - Day25-35 |
| ko00300; Lysine biosynthesis | 0.002383987 | 0.019852813 | Day15-24 | Day15-24 - Day25-35 |
| ko00643; Styrene degradation | 0.002090647 | 0.019852813 | Day15-24 | Day15-24 - Day25-35 |
| ko00270; Cysteine and methionine metabolism | 0.002443962 | 0.021979557 | Day15-24 | Day15-24 - Day25-35 |
| ko02040; Flagellar assembly | 0.002697924 | 0.021979557 | Day15-24 | Day15-24 - Day25-35 |
| ko05010; Alzheimers disease | 0.002903976 | 0.021979557 | Day15-24 | Day15-24 - Day25-35 |
| ko00280; Valine, leucine and isoleucine degradation | 0.003030708 | 0.030943195 | Day15-24 | Day15-24 - Day25-35 |
| ko00360; Phenylalanine metabolism | 0.003833813 | 0.030943195 | Day15-24 | Day15-24 - Day25-35 |
| ko00473; D-Alanine metabolism | 0.003743341 | 0.030943195 | Day25-35 | Day15-24 - Day25-35 |

